# An Extended Motif in the SARS-CoV-2 Spike Modulates Binding and Release of Host Coatomer in Retrograde Trafficking

**DOI:** 10.1101/2021.09.03.458953

**Authors:** Debajit Dey, Suruchi Singh, Saif Khan, Matthew Martin, Nicholas J. Schnicker, Lokesh Gakhar, Brian G. Pierce, S. Saif Hasan

## Abstract

β-Coronaviruses such as SARS-CoV-2 hijack coatomer protein-I (COPI) for spike protein retrograde trafficking to the progeny assembly site in endoplasmic reticulum-Golgi intermediate compartment (ERGIC). However, limited residue-level details are available into how the spike interacts with COPI. Here we identify an extended COPI binding motif in the spike that encompasses the canonical K-x-H dibasic sequence. This motif demonstrates selectivity for αCOPI subunit. Guided by an *in silico* analysis of dibasic motifs in the human proteome, we employ mutagenesis and binding assays to show that the spike motif terminal residues are critical modulators of complex dissociation, which is essential for spike release in ERGIC. αCOPI residues critical for spike motif binding are elucidated by mutagenesis and crystallography and found to be conserved in the zoonotic reservoirs, bats, pangolins, camels, and in humans. Collectively, our investigation on the spike motif identifies key COPI binding determinants with implications for retrograde trafficking.

## Introduction

β-Coronaviruses have been responsible for major human respiratory diseases in the last two decades. In 2002, the severe acute respiratory syndrome coronavirus (SARS-CoV) was implicated in an epidemic first reported in China before spreading to 27 countries, which resulted in 774 deaths ^1^. A decade later, Middle East respiratory syndrome (MERS) was reported in Saudi Arabia in 2012 with over 30% fatality in patients ^2,3^. Most recently, the novel SARS-CoV-2 has been implicated in the COVID-19 global pandemic that has claimed over four million lives. Current efforts to contain the pandemic are focused primarily on vaccinations using the viral spike protein that is responsible for SARS-CoV-2 entry into host cells ^4,5^. Fundamental insights into spike biogenesis will advance the understanding of how β-coronaviruses exploit host resources during viral infection and may potentially lead to the development of novel therapeutics.

The trimeric β-coronavirus spike is organized into an ecto-domain, a transmembrane domain, and a cytosolic domain ^6,7^ (**Figure 1**). In infected cells, the newly synthesized spike protein in ER is transported first to the Golgi, and then from Golgi to the ERGIC compartment, which is the site of β-coronavirus progeny assembly ^8–10^. This retrograde trafficking of the post-translationally modified spike from Golgi to ERGIC involves a cytosolic dibasic motif, K-x-H-x-x (Lys-x-His, where x is any amino acid) ^9,10^ (**Figure 1**). Such C-terminal dibasic motifs and variants such as K-x-K-x-x and K-K-x-x are widely reported in the cytosolic tail of host membrane proteins that undergo retrograde trafficking ^11–14^. As such, the β-coronavirus spike demonstrates molecular mimicry of dibasic trafficking motifs ^9,10^. This recycling of spike protein has been suggested to enhance interactions with the viral membrane (M) protein localized in ERGIC during progeny assembly and is crucial for spike maturation ^9,10,15^. These observations establish a key role of the dibasic motif in SARS-CoV and SARS-CoV-2 infection and propagation cycles. Interestingly, the spike dibasic motif and adjacent residues are completely conserved in sarbecoviruses, i.e., SARS-CoV and SARS-CoV-2, although sequence divergence in residues neighboring the dibasic motif is noted in MERS-CoV ^9,10^.

**Figure 1:**
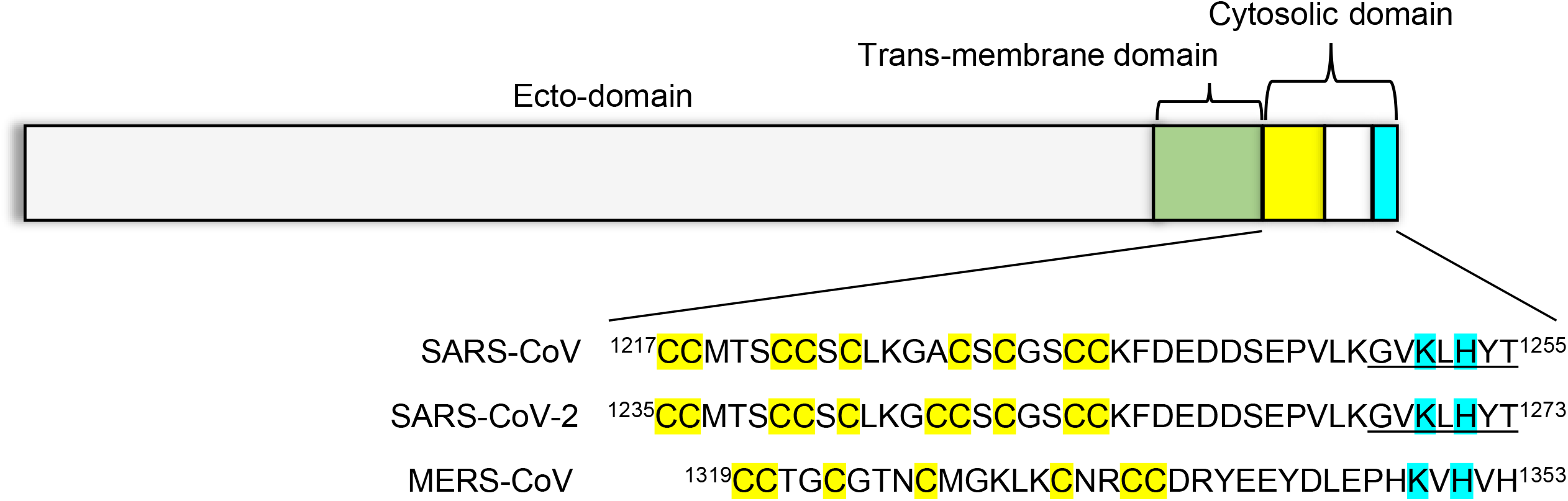
Organization of the coronavirus spike protein. The spike protein is divided into an ecto-domain (gray), a trans-membrane domain (green), and a cytosolic domain (yellow-white-cyan). The cytosolic domain includes a cysteine rich tract (yellow) and a dibasic motif for COPI interactions (cyan). This overall organization and the dibasic motif are conserved in the spike protein of SARS-CoV, SARS-CoV-2, and MERS-CoV, which have been implicated in wide-spread human disease. The underlined residues correspond to the peptide sequence utilized in this manuscript.

On the host side, retrograde trafficking is mediated by the interactions of dibasic motifs with the coatomer protein-I (COPI) complex ^11,16^. Seven subunits, namely, α, β, β’, γ, δ, ε and ζ, assemble into a COPI complex on retrograde trafficking vesicles that carry cargo ^17–24^. Prior genetic, biochemical, biophysical, and structural investigations have shown that the binding site for host cargo dibasic motifs maps to the N-terminal β-propeller WD40 domains of α and β’COPI subunits, which are structural homologs ^11,25–29^. Mutagenesis analyses of α and β’ subunit N-terminal WD40 domains have identified residues critical for binding of host protein dibasic motifs^27^. This and another study ^28^ provided important structural details of how dibasic host and viral peptides bind to αCOPI-WD40 and β’COPI-WD40 domains.

Cellular and biochemical investigations in recent years and during the ongoing COVID-19 pandemic have suggested a role of COPI interactions in sarbecovirus spike trafficking, maturation, glycan processing, and syncytia formation during infection ^10,15,30^. These studies have established a platform to investigate the underlying chemistry of spike-COPI interaction. For instance, it is not known whether the K-x-H motif is sufficient to determine the strength of this interaction, whether adjacent residues in the spike cytosolic tail play a role in this binding, and what are the sequence determinants of spike-COPI disassembly crucial for spike release in ERGIC. On the host side, it is presently not known which COPI residues are critical for spike interactions. As such, key facets of this initial binding event in spike trafficking remain largely unknown for SARS-CoV-2 as well as SARS-CoV. In the present investigation, we address these questions using a combination of bio-layer interferometry (BLI), molecular modeling, mutagenesis, X-ray crystallography, and an *in silico* analysis of the human membrane proteome. Employing a sarbecovirus spike hepta-peptide corresponding to the K-x-H-x-x motif, we identify critical residues in αCOPI-WD40 for hepta-peptide binding and demonstrate structural alterations in an αCOPI-WD40 mutant. Amino acid propensity is described in human dibasic motifs and adjacent downstream residues, and mutagenesis experiments driven by this analysis provide insights into how sarbecovirus spike modulates strength of binding to COPI. Collectively, our study advances the structural and biophysical understanding of how the dibasic motif hijacks COPI for spike retention in endo-membranes and trafficking to the plasma membrane during sarbecovirus infections.

## Results

### Direct binding of sarbecovirus spike hepta-peptide is selective for αCOPI-WD40 domain

In this investigation, we heterologously expressed and purified the N-terminal WD-40 domain of β’COPI-WD40 (residues 1-301) from *Saccharomyces cerevisiae* and αCOPI-WD40 (residues 1-327) from *Schizosaccharomyces pombe* (**Supplementary Figures S1, S2**). Although SARS-CoV-2 infects mammals, we chose these yeast constructs for two reasons. First, the putative interaction interface for dibasic peptides is conserved between these constructs and theαCOPI and β’COPI orthologs in humans, COPA and COPB2, respectively ^27,28^. Second, these domains have been previously crystallized and structurally characterized ^27,28^. This is consistent with our aim of understanding the structural basis of spike-COPI interactions. Although β’COPI-WD40 was expressed in *E. coli* as described previously, αCOPI-WD40 has reportedly presented challenges in protein production ^27,28^. Hence, the αCOPI-WD40 construct was expressed in Expi293 cells. This generated yields of purified αCOPI-WD40 of ~4 mg per 100 ml cell culture volume. For comparison with a previously published construct of αCOPI-WD40 expressed in *E. coli* ^28^, the crystal structure of the purified αCOPI-WD40 domain was determined to 1.8Å resolution (**Supplementary Table T1**). The αCOPI-WD40 domain is organized into a β-propeller and is consistent with previously described structures of αCOPI-WD40 (Cα root-mean-square-deviation is <0.5Å) ^28^. However, a peripheral loop and a short α-helix (Gly^168^-Ala^188^, shown with an arrow in **Figure 2a**) demonstrate substantial differences from previously described αCOPI-WD40 structures likely due to altered crystal packing. An N-terminal acetylation of the αCOPI-WD40 polypeptide was identified in the structure. Importantly, the αCOPI-WD40 domain interface for putative interactions with dibasic motifs is similar between the structure determined here and previously published structures ^28^.

**Figure 2:**
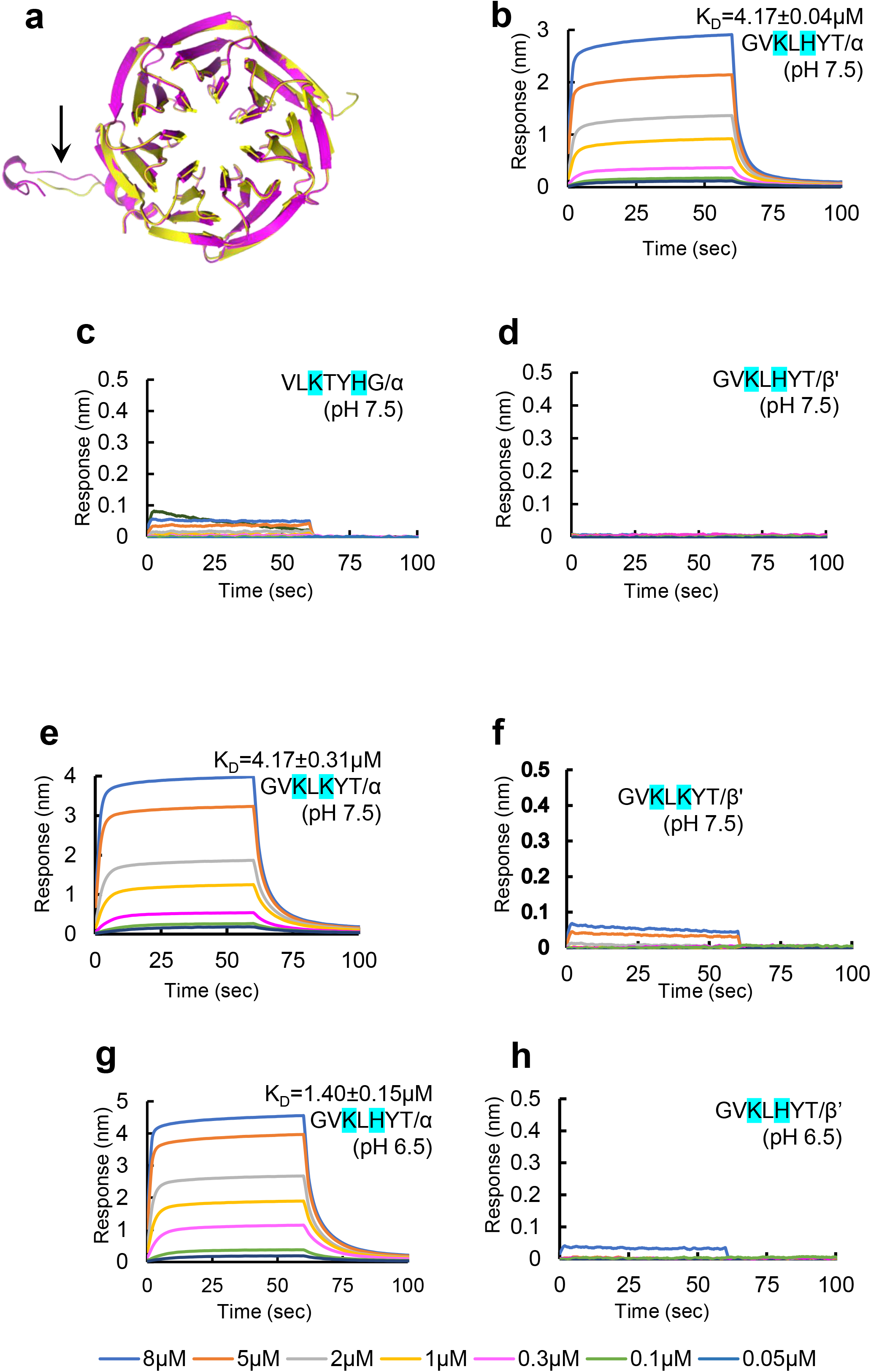
Direct binding interaction of sarbecovirus spike hepta-peptide with αCOPI-WD40. **(a)** Structural conservation of αCOPI-WD40 domain determined in the present investigation (yellow) and a previous structure (magenta). Arrow highlights main chain differences between these two αCOPI-WD40 structures in Gly^168^-Ala^188^. **(b-h)** BLI assay of N-biotinylated spike hepta-peptide with COPI-WD40 domain. One representative experiment of three is shown in panels **(b-h)**. Color code for concentrations is given at the bottom of the figure. The equilibrium K_D_ is provided with each sensorgram for comparison. **(b)** The spike wild type peptide sequence demonstrates dose-dependent binding to αCOPI-WD40 domain. **(c)** Scrambling of the hepta-peptide sequence abolishes binding suggesting sequence-specific interaction. **(d)** β’COPI-Wd40 demonstrates no interaction with the immobilized hepta-peptide. The mutant peptide, Gly-Val-Lys-Leu-Lys-Tyr-Thr, shows dose-dependent binding to **(e)** αCOPI-WD40 but not **(f)** β’COPI-WD40. **(g)** Acidification enhances binding between the wild-type spike hepta-peptide and αCOPI-WD40 domain. **(h)** β’COPI-WD40 shows weakly enhanced binding to the spike hepta-peptide upon acidification. “n.d.” implies not determined for weak interactions.

Recent investigations of SARS-CoV-2 spike and previously of SARS-CoV spike with COPI have employed cellular lysates ^10,15,30^. Hence, we asked if there is direct interaction between the purified components. To address this question, we first established a BLI assay to test this interaction. Monomeric constructs of the spike cytosolic domain demonstrate similar interactions with COPI as the trimeric cytosolic domain ^10^. Hence, a hepta-peptide representing a monomeric dibasic motif in the sarbecovirus spike (^1267^Gly-Val-Lys-Leu-His-Tyr-Thr^1273^) was synthesized with an N-terminal biotin tag attached via a linker. The hepta-peptide C-terminus has a free carboxylate to mimic the C-terminus of a polypeptide. This hepta-peptide (or its sequence variants) was immobilized on a streptavidin biosensor for BLI analysis. The purified α or β’COPI-WD40 domain was provided as the analyte in the BLI assay (**Figure 2b-h, Table 1**). It was observed that the spike hepta-peptide binds directly to the purified αCOPI-WD40 domain with an equilibrium dissociation constant (K_D_)=4.17±0.04 μM and a kinetic K_D_=2.75±0.09 μM at pH 7.5 (**Figure 2b, Table 1**). A scrambled sequence of this hepta-peptide showed no detectable interaction with αCOPI-WD40 (**Figure 2c**) suggesting that the binding of the wild-type hepta-peptide was sequence specific. This binding analysis demonstrates that the C-terminal peptide of the sarbecovirus spike contains sufficient sequence and structural information to interact directly with αCOPI-WD40. This is consistent with prior COPI binding analyses with peptides corresponding to host dibasic motifs ^27,28^. In contrast, this sarbecovirus hepta-peptide demonstrates a lack of binding to β’COPI-WD40 (**Figure 2d**). This selectivity for αCOPI-WD40 is consistent with that reported for a similar spike hepta-peptide (^1377^Phe-Glu-Lys-Val-His-Val-Gln^1383^) from porcine epidemic diarrhea virus (PEDV), an α-coronavirus ^28^. Mutation of the K-x-H motif to K-x-K in the sarbecovirus mutant peptide, i.e., ^1267^Gly-Val-Lys-Leu-Lys-Tyr-Thr^1273^, demonstrates similar selectivity for αCOPI-WD40 over β’COPI-WD40 (**Figure 2e, f**). Cellular studies suggest enhanced interactions of this mutant spike sequence with COPI subunits ^15,30^. It is likely that this mutation affects the local conformation of the full-length spike protein leading to modulation of COPI interactions in a cellular environment.

**Table 1:** Dissociation and rate constants for the interaction of spike hepta-peptides with COPI-WD40 domains.

| Peptide sequence<sup>a</sup> | COPI-WD40 analyte<sup>b</sup> | pH | Equilibrium $K_D$ (μM) | Kinetic $K_D$ (μM) | $k_{on}$ ($10^3$/Ms) | $k_{off}$ ($10^{-1}$/s) |
| --- | --- | --- | --- | --- | --- | --- |
| GVKLHYT | $\alpha$ | 7.5 | $4.17 \pm 0.04$ | $2.75 \pm 0.09$ | $89.96 \pm 2.72$ | $2.50 \pm 0.03$ |
| GVKLHYT | $\beta'$ | 7.5 | n.d. | n.d. | n.d. | n.d. |
| VLKTYHG | $\alpha$ | 7.5 | n.d. | n.d. | n.d. | n.d. |
| GVKLKYT | $\alpha$ | 7.5 | $4.17 \pm 0.31$ | $2.71 \pm 0.05$ | $67.3 \pm 1.45$ | $1.84 \pm 0.01$ |
| GVKLKYT | $\beta'$ | 7.5 | n.d. | n.d. | n.d. | n.d. |
| GVKLHYT | $\alpha$ | 6.5 | $1.40 \pm 0.15$ | $0.95 \pm 0.02$ | $152.13 \pm 3.45$ | $1.45 \pm 0.01$ |
| GVKLHYT | $\beta'$ | 6.5 | n.d. | n.d. | n.d. | n.d. |
| GVALHYT | $\alpha$ | 6.5 | n.d. | n.d. | n.d. | n.d. |
| GVKLAYT | $\alpha$ | 6.5 | n.d. | n.d. | n.d. | n.d. |
| GVALAYT | $\alpha$ | 6.5 | n.d. | n.d. | n.d. | n.d. |
| GVKLHAT | $\alpha$ | 6.5 | $3.77 \pm 0.34$ | $2.32 \pm 0.07$ | $120.87 \pm 3.39$ | $2.80 \pm 0.03$ |
| GVKLHYA | $\alpha$ | 6.5 | $1.13 \pm 0.05$ | $1.14 \pm 0.02$ | $66.67 \pm 0.80$ | $0.76 \pm 0.01$ |
| GVKLHYV | $\alpha$ | 6.5 | $3.20 \pm 0.04$ | $2.06 \pm 0.05$ | $102.67 \pm 2.61$ | $2.12 \pm 0.02$ |
| GVKLHYE | $\alpha$ | 6.5 | $0.31 \pm 0.01$ | $0.38 \pm 0.01$ | $58.96 \pm 0.54$ | $0.24 \pm 0.003$ |
| GVKLHYQ | $\alpha$ | 6.5 | $0.78 \pm 0.04$ | $1.15 \pm 0.01$ | $60.05 \pm 0.75$ | $0.66 \pm 0.01$ |
| GVKLHYR | $\alpha$ | 6.5 | $1.73 \pm 0.24$ | $1.29 \pm 0.04$ | $57.60 \pm 0.61$ | $1.77 \pm 0.01$ |
| KEVYLHG | $\alpha$ | 6.5 | n.d. | n.d. | n.d. | n.d. |
| GVKLHYT | α-Ala<sup>57</sup> | 6.5 | n.d. | n.d. | n.d. | n.d. |
| GVKLHYT | α-Ala<sup>115</sup> | 6.5 | n.d. | n.d. | n.d. | n.d. |
| GVKLHYT | α-Ala<sup>139</sup> | 6.5 | n.d. | n.d. | n.d. | n.d. |

<sup>a</sup>The sequences of immobilized sarbecovirus spike peptides are given in the left column. The dibasic motif residues are highlighted in bold. Mutations in the peptides relative to the wild-type sequence are highlighted in bold and are underlined.
<sup>b</sup> $\alpha$ and $\beta'$ represent wild-type αCOPI-WD40 and $\beta'$ COPI-WD40, respectively. $\alpha$ -Ala<sup>57</sup>, $\alpha$ -Ala<sup>115</sup>, and $\alpha$ -Ala<sup>139</sup>, represent the Arg<sup>57</sup>→Ala, Asp<sup>115</sup>→Ala, and Tyr<sup>139</sup>→Ala mutants of αCOPI-WD40, respectively.

One of the first analyses of COPI involvement in SARS-CoV spike trafficking employed cellular lysate pull-downs to show enhanced spike-COPI interactions under acidification to pH 6.5 ^10^. As the sarbecovirus hepta-peptide contains His^1271^ in the K-x-H motif, we asked if this acidification would affect hepta-peptide interaction with αCOPI-WD40. A nearly 3-fold enhancement in binding between the wild-type hepta-peptide and αCOPI-WD40 (equilibrium K_D_=1.40±0.15 μM) was observed upon acidification to pH 6.5, likely due to partial protonation of the His residue in the hepta-peptide K-x-H motif (**Figure 2g, Table 1**). Relative to pH 7.5, this lower pH accelerated the association rate of the hepta-peptide with αCOPI-WD40 by a factor of 1.7 while concomitantly suppressing complex dissociation by another 1.7-fold (**Table 1**). As such, acidification was inferred to be a key factor in stabilizing the hepta-peptide complex with αCOPI-WD40. All subsequent BLI assays were performed at pH 6.5.

The binding of β’COPI-WD40 to the wild type spike peptide is still substantially weaker than that with αCOPI-WD40 in low pH (**Figure 2h**).

### The terminal residues in the spike hepta-peptide are key modulators of αCOPI-WD40 binding and dissociation

Homology modeling was employed to analyze the structural basis of interaction between SARS-CoV-2 spike hepta-peptide and αCOPI-WD40 domain. This modeling was based on a prior co-crystal structure of αCOPI-WD40 domain with a dibasic peptide ^28^. Apart from the Lys^1269^ and His^1271^ residues in the K-x-H motif, the terminal Tyr^1272^-Thr^1273^ residues in the hepta-peptide are within interaction distance of αCOPI-WD40 surface residues (**Figure 3a**). This is intriguingly suggestive of a role of these two spike residues in binding αCOPI. Interestingly, the two N-terminal residues in the hepta-peptide, i.e., Gly^1267^-Val^1268^, make no contact with the αCOPI-WD40 surface. To evaluate the role of the spike residues in binding αCOPI-WD40, *in silico* alanine scanning mutagenesis of the modeled spike hepta-peptide was performed (**Table 2**). The spike Lys^1269^ and His^1271^ residues that constitute the K-x-H dibasic motif are predicted to be most crucial for binding αCOPI-WD40 domain. The *in silico* mutations of these residues to Ala yield highly unfavorable free energy changes suggestive of substantially weakened binding to αCOPI-WD40 (**Table 2**). The Ala mutation of Tyr^1272^ in the spike peptide implies a substantial role of this residue in stabilization of the spike-αCOPI-WD40 complex. This is likely due to the side chain interaction between the oxygen atom in Tyr^1272^ side-chain hydroxyl group with the αCOPI-WD40 His^31^ side-chain NE2 atom, along with main chain interactions of Tyr^1272^. The terminal residue in the spike, i.e., Thr^1273^, is predicted to contribute modestly to the stabilization of the complex with αCOPI-WD40 (**Table 2**).

**Figure 3:**
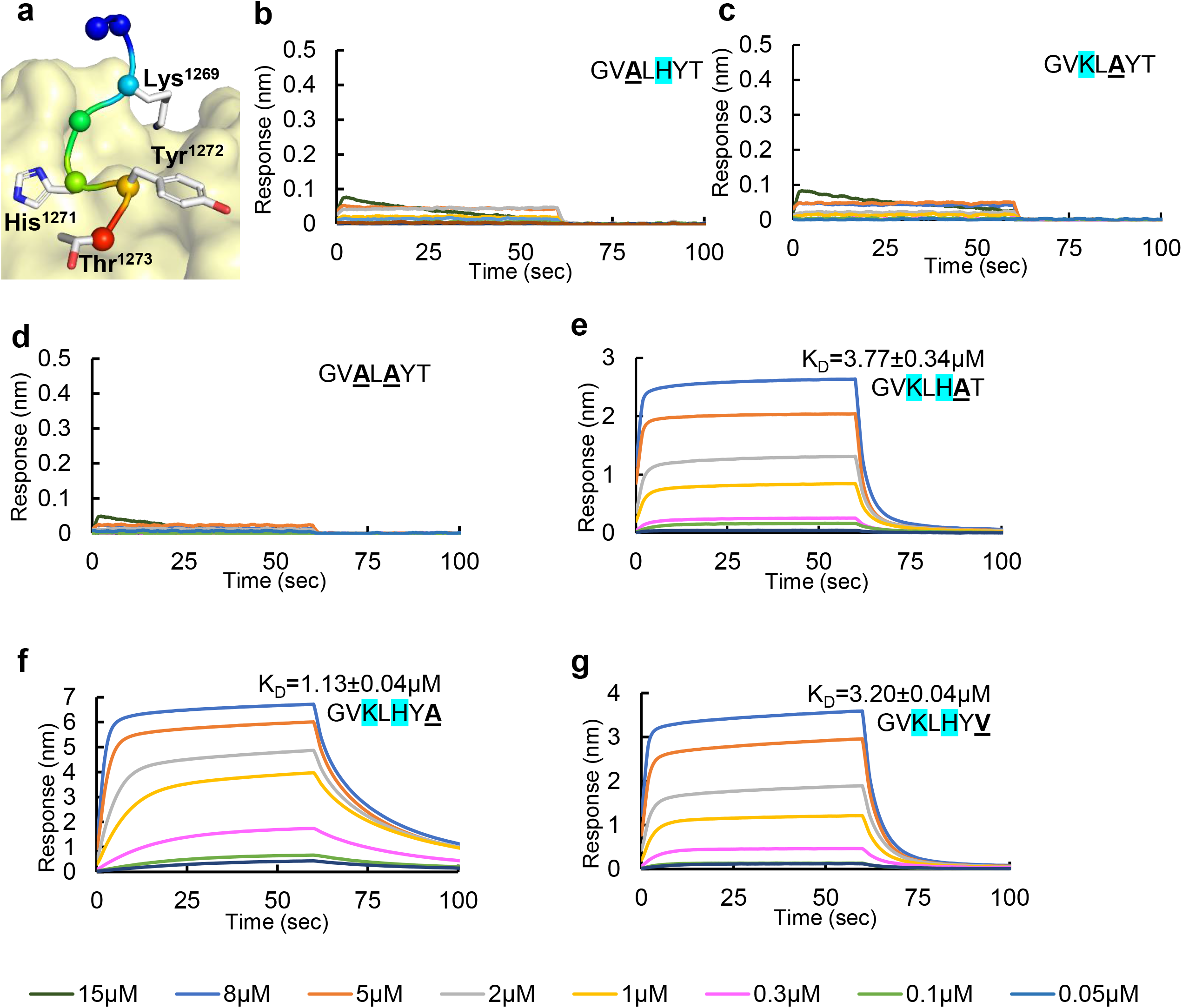
Structure-guided mutagenesis of spike hepta-peptide and binding analysis with αCOPI-WD40 domain. **(a)** *In silico* model of the spike hepta-peptide complexed with αCOPI-WD40 domain (yellow surface). The hepta-peptide is shown as a ribbon in rainbow colors from N (blue) to C (red) terminus. The Cα-atoms in the hepta-peptide are shown as spheres. The side chains of residues that interact with αCOPI-WD40 are shown as a stick. **(b-g)** BLI analyses of αCOPI-WD40 binding to spike hepta-peptide mutants. The color code of BLI traces is given at the bottom of the figure. One representative experiment of three is shown. Color code for concentrations is given at the bottom of the figure. The mutation in the spike hepta-peptide sequence is highlighted in bold and is underlined. The equilibrium K_D_ is provided with each sensorgram for comparison. Mutagenesis of, **(b)** Lys^1269^, **(c)** His^1271^, or **(d)** both abolishes binding to αCOPI-WD40. **(e)** In contrast, Tyr^1272^→Ala mutation only weakens binding to αCOPI-WD40. The middle panel shows weak binding of αCOPI-WD40 domain with a hepta-peptide wherein Lys^1269^ has been mutated to Ala. **(f)** Mutagenesis of Thr^1273^ to Ala in the spike hepta-peptide leads to moderately enhanced binding to αCOPI-WD40 whereas mutagenesis to Val^1273^ weakens binding **(g)**.

**Table 2:**
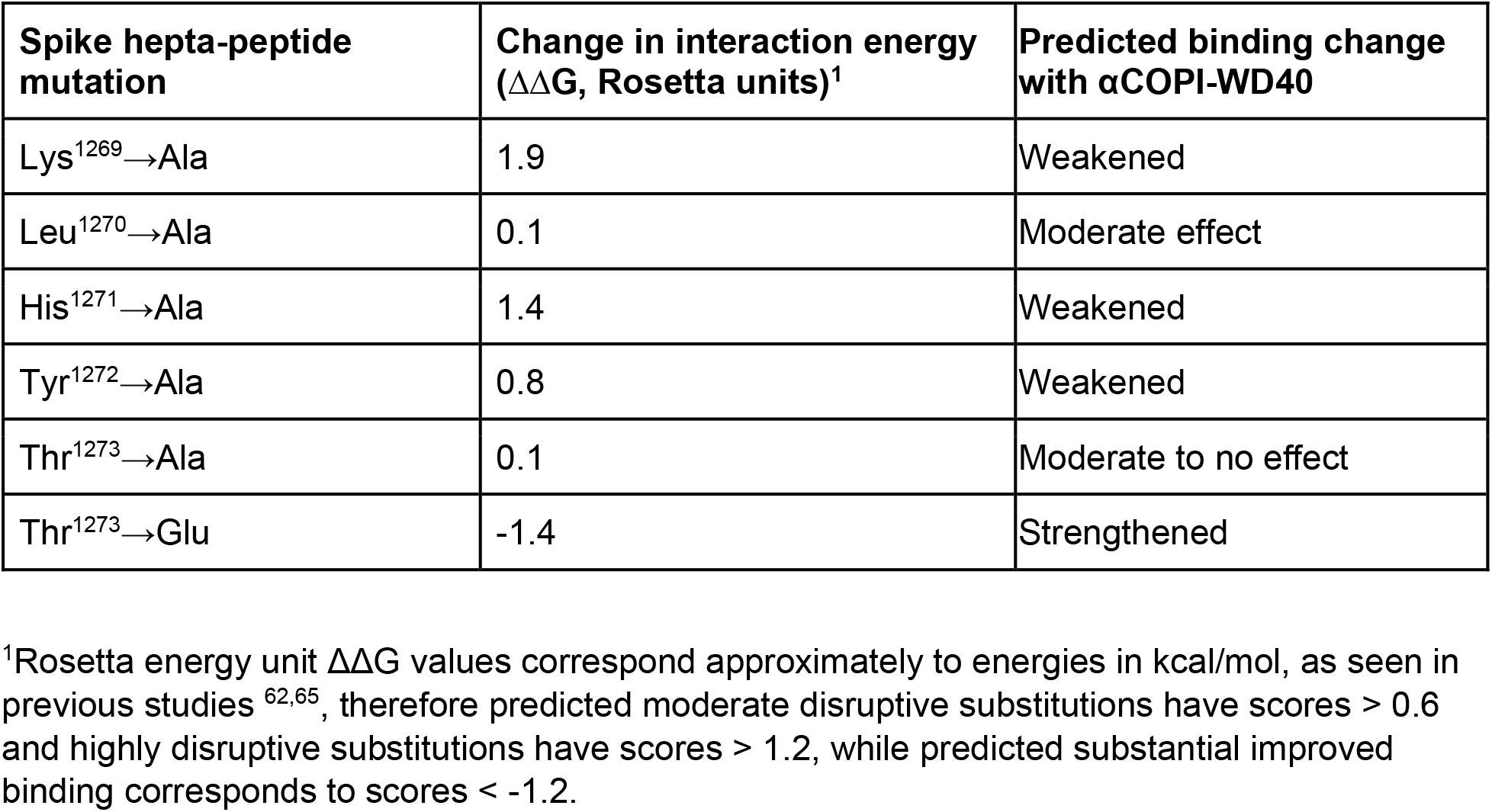
*In silico* mutagenesis of extended dibasic motif in SARS-CoV-2 spike.

| **Spike hepta-peptide mutation** | **Change in interaction energy ($\Delta\Delta G$, Rosetta units)<sup>1</sup>** | **Predicted binding change with αCOPI-WD40** |
| --- | --- | --- |
| Lys<sup>1269</sup>→Ala | 1.9 | Weakened |
| Leu<sup>1270</sup>→Ala | 0.1 | Moderate effect |
| His<sup>1271</sup>→Ala | 1.4 | Weakened |
| Tyr<sup>1272</sup>→Ala | 0.8 | Weakened |
| Thr<sup>1273</sup>→Ala | 0.1 | Moderate to no effect |
| Thr<sup>1273</sup>→Glu | -1.4 | Strengthened |

<sup>1</sup>Rosetta energy unit $\Delta\Delta G$ values correspond approximately to energies in kcal/mol, as seen in previous studies <sup>62,65</sup>, therefore predicted moderate disruptive substitutions have scores > 0.6 and highly disruptive substitutions have scores > 1.2, while predicted substantial improved binding corresponds to scores < -1.2.

Next, we tested this *in silico* model of interactions between the spike hepta-peptide and αCOPI-WD40 using BLI assays (**Figure 3b-g**). The mutagenesis of Lys^1269^ or His^1271^ in the spike K-x-H motif to Ala residues abolished binding to αCOPI-WD40 (**Figure 3b, c**). As expected, the dual Ala mutation of the K-x-H motif does not demonstrate any substantial binding to αCOPI-WD40 (**Figure 3d**). As such, both basic residues in SARS-CoV-2 spike K-x-H motif are individually and concomitantly required for αCOPI-WD40 binding. Replacement of either residue is sufficient to disrupt αCOPI-WD40 binding to the spike hepta-peptide. These data are consistent with the *in silico* predictions described above as well as with cellular assays on SARS-CoV and SARS-CoV-2 spike trafficking ^10,15,30^. We next tested the contribution of spike hepta-peptide Tyr^1272^ residue to αCOPI-WD40 binding. A BLI assay of a mutant Tyr^1272^→Ala spike hepta-peptide (^1267^Gly-Val-Lys-Leu-His-<u>Ala</u>-Thr^1273^) yielded an equilibrium K_D_=3.77±0.34 μM, which is 2.7-fold weaker than the wild-type spike peptide (**Figure 3e, Table 1)**. Although this mutation only reduced the rate of complex formation by 1.3-fold relative to the wild-type hepta-peptide (**Table 1**), it accelerated complex dissociation by 1.9-fold (**Table 1**). This suggests weakened interactions of the spike hepta-peptide with αCOPI-WD40 when the aromatic side chain interactions of Tyr^1272^ are abrogated. Collectively, this BLI analysis indicates that Tyr^1272^ is important for complex stability. These experimental results are consistent with the above described *in silico* model (**Table 3**). Next, we evaluated the C-terminal position Thr^1273^ in the spike. A BLI assay of a mutant Thr^1273^→Ala hepta-peptide (^1267^Gly-Val-Lys-Leu-His-Tyr-<u>Ala</u>^1273^) yielded an equilibrium KD=1.13±0.05 μM, which is similar to the wild type hepta-peptide (KD=1.40±0.15 μM) (**Figure 3f, Table 1**). This Thr^1273^→Ala mutation caused a slowing down of αCOPI-WD40 association-dissociation kinetics by 2.3 and 1.9-fold respectively (**Table 1**). This suggested that a β-branched residue at the C-terminus may be an important determinant in complex formation kinetics. To probe further, we generated a Thr^1273^→Val mutant hepta-peptide, which maintains a β-branched residue at the C-terminus. Val has methyl groups at the two side chain γ positions, which replace a methyl and a hydroxyl group at equivalent γ positions in Thr. A BLI assay of this mutant hepta-peptide showed a 2.3-fold weakened interaction with αCOPI-WD40 relative to the wild type, with an equilibrium K_D_=3.20±0.04 μM (**Figure 3g, Table 1**). Interestingly, this mutant demonstrated 1.5-fold slower association kinetics and 1.5-fold more rapid dissociation than for the wild type hepta-peptide (**Table 1**). Compared to the Thr^1273^→Ala hepta-peptide, this Thr^1273^→Val mutant weakened binding by 2.8-fold while accelerating αCOPI-WD40 complex association and dissociation kinetics by 1.5 and 2.8-fold respectively (**Table 1**). These data suggest that the spike C-terminal Thr residue side-chain provides interactions that modulate dissociation of the complex with αCOPI-WD40. Interestingly, a prior analysis implicated β-branched residues at the penultimate position in the PEDV spike sequence in modulating interactions with COPI-WD40 domains ^28^.

**Table 3:** Interaction residues in *in silico* model of spike hepta-peptide with αCOPI-WD40.

| Residue in SARS-CoV-2 spike hepta-peptide | αCOPI-WD40 contact residue within 3.5Å |
| --- | --- |
| Lys<sup>1269</sup> | Asp<sup>96</sup>, Asp<sup>115</sup>, Tyr<sup>139</sup> |
| Leu<sup>1270</sup> | Arg<sup>99</sup> |
| His<sup>1271</sup> | Arg<sup>57</sup>, Arg<sup>99</sup>, Met<sup>141</sup>, Leu<sup>157</sup>, Asn<sup>216</sup> |
| Tyr<sup>1272</sup> | His<sup>31</sup>, Arg<sup>57</sup> |
| Thr<sup>1273</sup> | Arg<sup>13</sup>, Lys<sup>15</sup>, His<sup>31</sup>, Arg<sup>57</sup>, Arg<sup>300</sup>, Trp<sup>302</sup> |

**Table 4:** Effects of *in silico* αCOPI-WD40 mutagenesis on spike hepta-peptide binding.

| αCOPI-WD40 mutation | Change in interaction energy ($\Delta\Delta G$, Rosetta Units) | Predicted binding change with αCOPI-WD40 |
| --- | --- | --- |
| Lys<sup>15</sup>→Ala | 0.8 | Weakened |
| His<sup>31</sup>→Ala | 0.8 | Weakened |
| Arg<sup>57</sup>→Ala | 1.2 | Weakened |
| Asp<sup>115</sup>→Ala | 0.9 | Weakened |
| Tyr<sup>139</sup>→Ala | 1.1 | Weakened |

### Electrostatics of spike hepta-peptide C-terminus drive dissociation from αCOPI-WD40

We performed an *in silico* analysis of the human proteome to gain insights into whether the spike extended dibasic motif demonstrates consistency with host dibasic motifs and their environment. We identified 119 sequences predicted to be membrane proteins that terminate with K-x-H-x-x and K-x-K-x-x dibasic motifs (**Supplementary Table T2**). These sequences were aligned and analyzed for the frequency of 20 amino acids at each of the positions in the dibasic motif and the two terminal residues following the motif (**Figure 4a, Supplementary Table T3**). This analysis revealed novel details about the dibasic motif. First, it was inferred that the predominant dibasic motif is K-x-K-x-x rather than K-x-H-x-x by nearly an order of magnitude. Second, only a low frequency (0.07) of the sequences has an aromatic residue at the penultimate position, which corresponds to Tyr^1272^ in the SARS-CoV-2 spike. β-Branched residues Leu, Ile, Val, Ser, and Thr are found at a high frequency of 0.38 at this penultimate position. Third, acidic residues at the C-terminus are observed in nearly a quarter (frequency=0.24) of the sequences. Overall, with a frequency of 0.42, the C-terminal position has a strong tendency to be occupied by charged residues such as Arg, Asp, Glu, His, and Lys. Hydroxyl side chain containing Thr, which corresponds to Thr^1273^ in SARS-CoV-2 spike, is a low frequency residue (0.05). In our *in silico* model of the hepta-peptide complexed with αCOPI-WD40, the side chain of this Thr^1273^ residue is within interaction distance of a cluster of basic residues in αCOPI-WD40 (Arg^13^, Lys^15^, and Arg^300^, **Figure 4b**). Hence, we hypothesized that the presence of a charged residue at this spike position would modulate interactions with αCOPI-WD40. This was supported by our *in silico* analysis, which predicted stabilization of the complex when an acidic Glu residue replaced Thr^1273^ in the spike hepta-peptide (**Table 2**).

**Figure 4:**
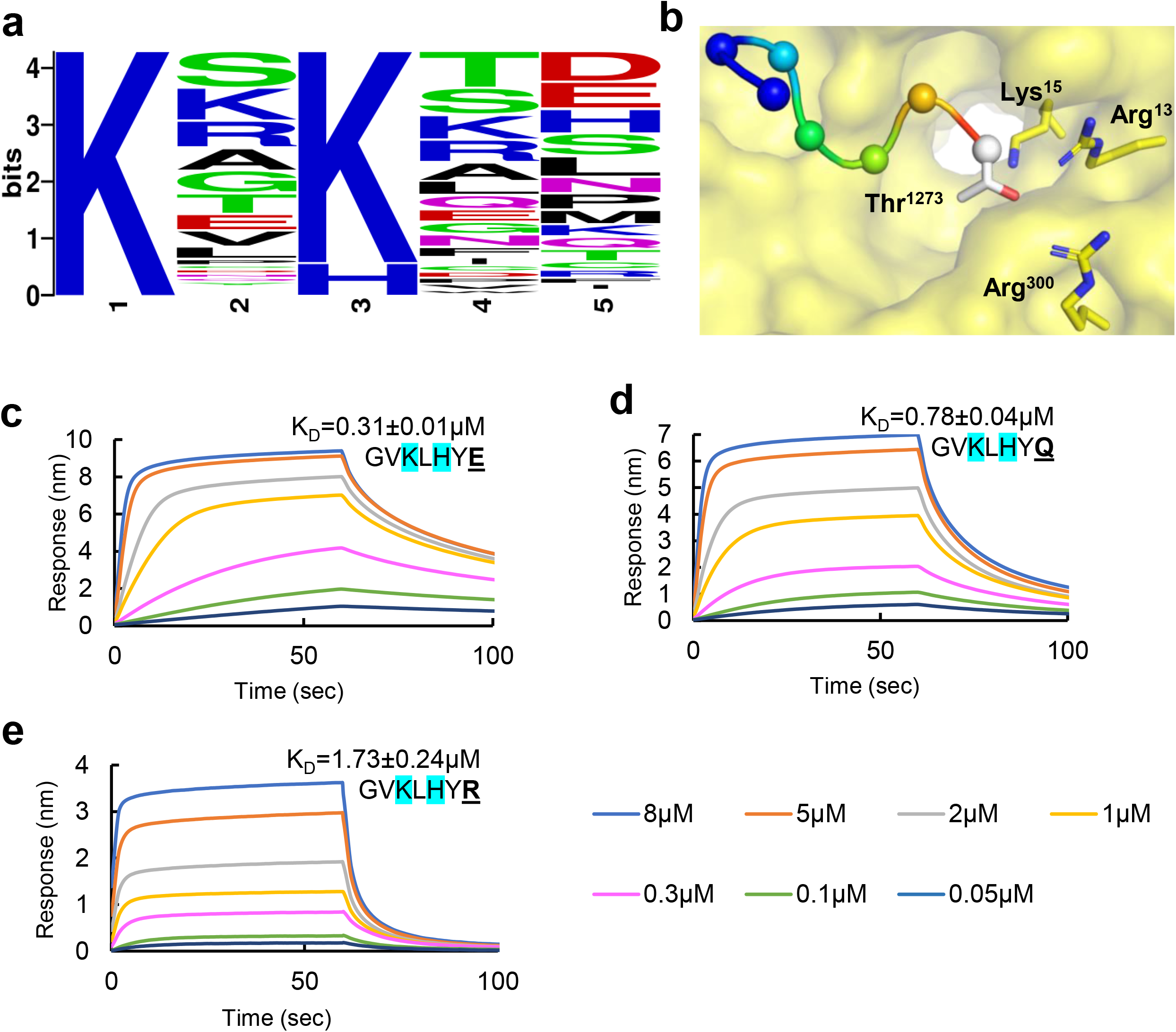
*In silico* and biophysical analysis of spike C-terminal position in αCOPI-WD40 binding. **(a)** Sequence logo generated from the alignment of K-x-H(K)-x-x sequence in 119 proteins predicted to be in the human membrane proteome. This shows the abundance of Lys in the first and third positions, low frequency of aromatic residues in the penultimate position, and the abundance of Asp and Glu in the C-terminal position. **(b)** An *in silico* model of the spike hepta-peptide on αCOPI-WD40 (yellow surface) shows an abundance of basic residues in the vicinity of the terminal Thr^1273^ spike residue. Panels **(c-e)** show results of a BLI analysis of binding between spike hepta-peptide mutants and αCOPI-WD40. The equilibrium K_D_ is provided with each sensorgram for comparison. Color code for concentrations is given at the bottom of the figure. Stabilization of the spike hepta-peptide complexed with αCOPI-WD40 is observed when the terminal position contains either, **(c)** acidic Glu^1273^, or **(d)** neutral Gln^1273^ residue. **(e)** In contrast, basic Arg^1273^ in the spike hepta-peptide does not favor enhanced binding. These data show a role of this terminal hepta-peptide position in modulating tight binding to αCOPI-WD40. In **(c-e)**, one representative experiment of three is shown.

We next tested the role of this spike C-terminal residue in modulating αCOPI-WD40 binding using a BLI assay (**Figure 4c-e**). We employed three distinct mutations of the spike hepta-peptide at this position, i.e., acidic (Glu), basic (Arg), and neutral (Gln) (**Figure 4c-e**). The presence of an acidic Glu residue at the C-terminus was found to substantially strengthen binding of the hepta-peptide to αCOPI-WD40 with an equilibrium K_D_=0.31±0.01 μM, which is 4.5-fold tighter than the binding of the wild type hepta-peptide sequence (**Figure 4c, Table 2**). This is consistent with our *in silico* model and is strongly suggestive of an electrostatic interaction between the Glu^1273^ side chain and αCOPI Arg^13^, Lys^15^, and Arg^299^ side chains to stabilize the complex. Furthermore, the rate of dissociation of αCOPI-WD40 domain from the Glu^1273^ hepta-peptide is 6-fold slower than that of the wild-type spike hepta-peptide (**Table 1**). In fact, during the time course of our experiment, we did not observe complete dissociation of this complex with the Glu^1273^ containing hepta-peptide. To eliminate the possibility of non-specific interactions, we employed a hepta-peptide with a scrambled sequence (**Supplementary Figure S3**). Next, we tested whether modifying side-chain charge at the C-terminal position of the spike hepta-peptide affects complex formation with αCOPI-WD40. Relative to Glu^1273^, the binding between αCOPI-WD40 domain was weakened when neutral Gln^1273^ was substituted into the hepta-peptide (equilibrium K_D_=0.78±0.04 μM, **Figure 4d**). However, the binding of the hepta-peptide with Gln^1273^ was still 1.8-fold tighter than that of the wild-type hepta-peptide (**Table 1**). In contrast, basic Arg^1273^ in the spike hepta-peptide (equilibrium K_D_=1.73±0.24 μM) yielded an interaction strength similar to the wild type sequence (equilibrium K_D_=1.40±0.15 μM, **Figure 4e**). The amide carbonyl group in the Gln^1273^ side-chain likely interacts with the basic residue cluster on αCOPI-WD40 through hydrogen bonding. This stabilizing interaction is disrupted when Gln is replaced by Arg^1273^ in the spike hepta-peptide. Intriguingly, the rate of association with αCOPI-WD40 was slowed down relative to the wild type hepta-peptide by a factor of 2.6 for Glu^1273^ and Gln^1273^, whereas it was similar to that of Arg^1273^ containing hepta-peptide (**Table 2**). Overall, these data establish a critical role of the C-terminal position in the SARS-CoV-2 spike in modulating binding to αCOPI-WD40.

### Low frequency of acidic residues at the spike C-terminus

Given the inference that a C-terminal acidic residue strengthens binding to αCOPI-WD40, which likely interferes with spike release, we asked if a typical coronavirus spike demonstrates a paucity for such C-terminal acidic residues. A sequence and phylogenetic analysis of the five C-terminal residues in the spike protein of coronaviruses was performed to determine the frequency of Asp and Glu at the C-terminus (**Figure 5**). None of the coronavirus spike proteins that demonstrate the K-x-H-x-x, K-x-K-x-x, or K-x-R-x-x dibasic motif has a C-terminal acidic residue. It has been suggested that bats are the likely genetic source of human β-coronaviruses ^31–33^. Apart from bats, during the SARS-CoV epidemic and the ongoing SARS-CoV-2 pandemic, zoonotic reservoirs such as civets and pangolins have been suggested to be involved in coronavirus transmission ^33-38^. Our phylogenetic analysis showed that the extended coatomer binding motif in the spike, i.e., ^1269^Lys-Leu-His-Tyr-Thr^1273^, is conserved in coronavirus isolates from these animals (**Figure 5**). Moreover, this conservation of the extended motif is seen in the WHO deemed emerging variants of concern for SARS-CoV-2, i.e., α (Pango lineage B.1.1.7), β (B.1.351), γ (P.1), δ (B.1.617.2), and the latest Ο (B.1.1.529) ^39^. This is indicative of a strong selection pressure to maintain this COPI-interacting sequence in the spike protein.

**Figure 5:**
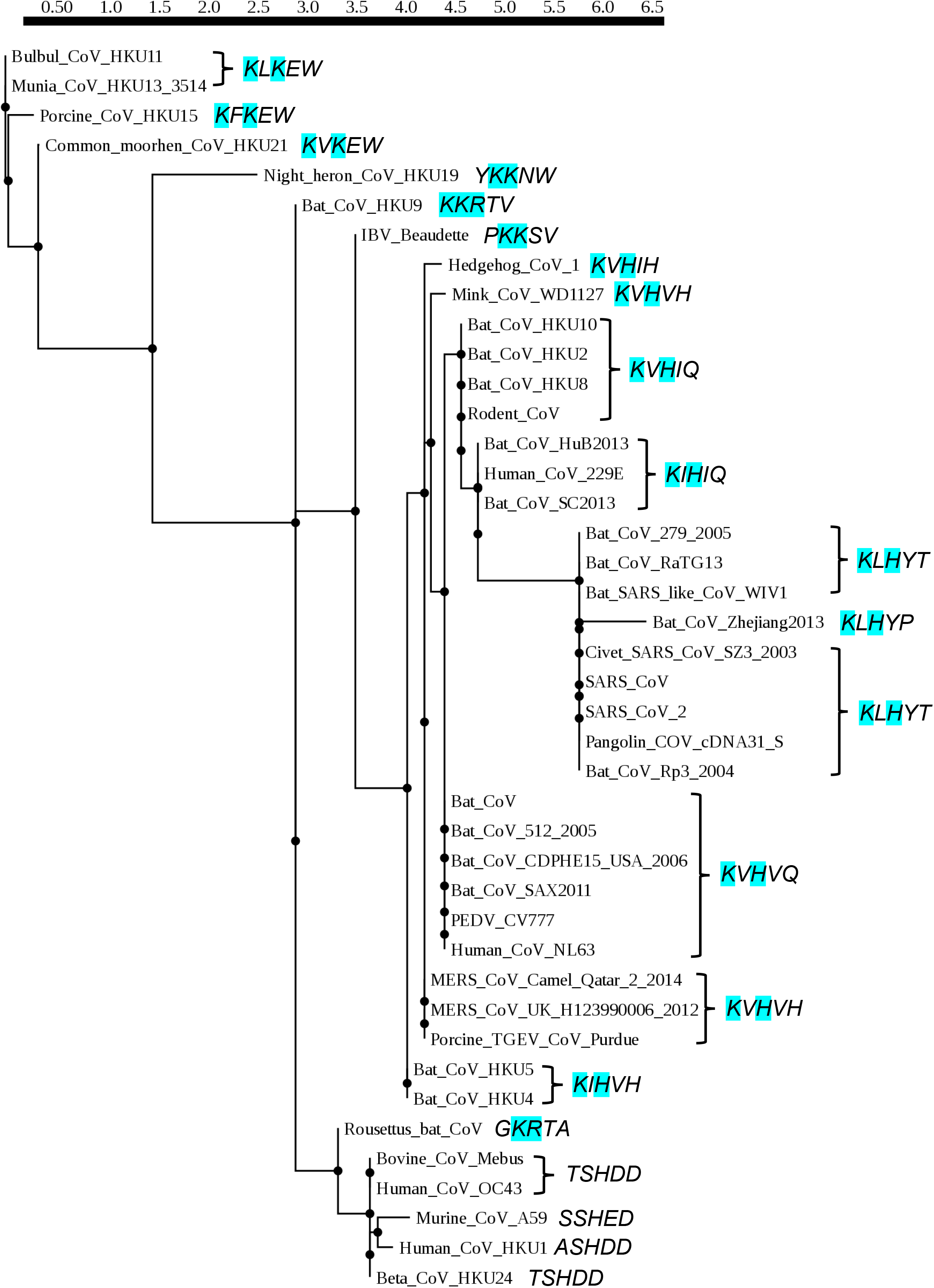
Phylogenetic analysis of coronavirus spike C-terminal sequence. This phylogram was generated from the alignment of five C-terminal residues in the spike protein. This penta-residue spike sequence is shown in italics to the right of each coronavirus. The residues in the dibasic motif are highlighted in cyan. The C-terminal acidic residues are highlighted in red. An acidic Asp is seen at the C-terminus of the spike proteins of only those coronaviruses that lack a dibasic motif.

### A polar αCOPI-WD40 interface for spike hepta-peptide binding

We subsequently focused our attention on the spike binding residues in αCOPI-WD40. The *in silico* modeling of SARS-CoV-2 spike hepta-peptide shows that interaction with αCOPI-WD40 domain involves predominantly polar residues (**Table 3**). Amongst these residues, Arg^57^, Asp^115^, and Tyr^139^ provide the highest level of stabilization to spike hepta-peptide binding. The Arg^57^ side-chain interacts with the main chain carbonyl of spike His^1271^, Tyr^1272^, and Thr^1273^ (**Figure 6a**). The Asp^115^ side-chain forms a bond with the terminal NZ atom in the spike Lys^1269^ side-chain (**Figure 6b**). This side-chain of spike Lys^1269^ is further stabilized by an interaction with the hydroxyl oxygen in Tyr^139^ side-chain (**Figure 6c**). Hence, the side chains of αCOPI-WD40 Arg^57^, Asp^115^, and Tyr^139^ residues provide an extensive and polar interaction network for binding of the spike hepta-peptide. Therefore, mutagenesis of these three αCOPI-WD40 residues to Ala is predicted to disrupt interactions with the spike hepta-peptide as suggested by our *in silico* analysis (**Table 3**).

**Figure 6:**
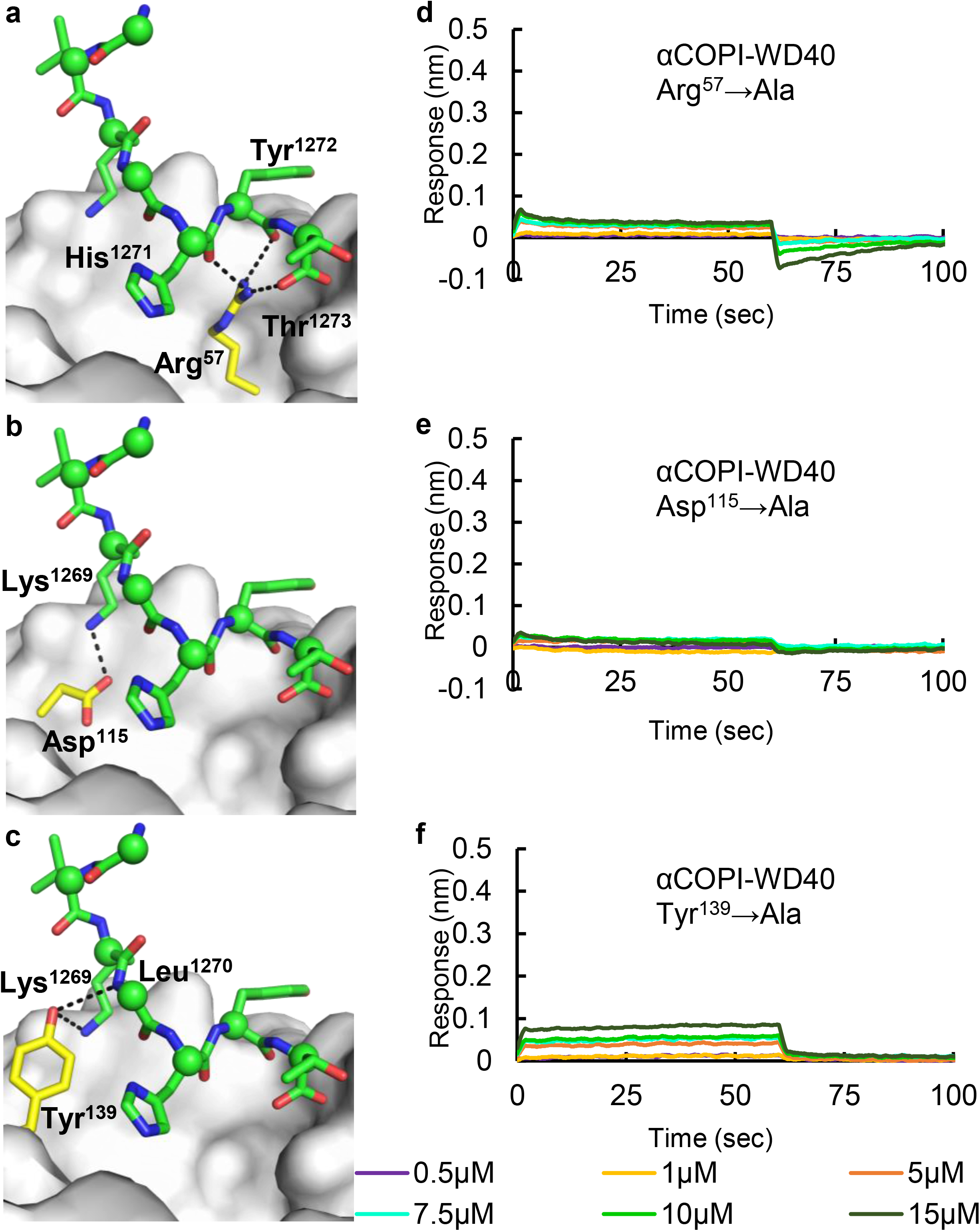
Structure-guided mutagenesis of αCOPI-WD40 domain and binding analysis with spike hepta-peptide. Panels **(a-c)** highlight the hepta-peptide interactions (within 4Å) of **(a)** Arg^57^, Asp^115^, and **(c)** Tyr^139^ residues in αCOPI-WD40 in an *in silico* model. These three interacting αCOPI-WD40 residues are shown as yellow-red-blue sticks whereas the other residues are shown as a yellow surface for simplicity. The corresponding interacting residues in the spike hepta-peptide are labelled and shown as green-red-blue sticks and spheres for Cα atoms. The BLI analysis of Arg^57^→Ala, Asp^115^→Ala, and Tyr^139^→Ala mutants with the wild type spike hepta-peptide is shown in panels **(d)**, **(e)**, and **(f)** respectively. All three mutants demonstrate no substantial binding of the spike hepta-peptide. One representative experiment of three is shown in panels **(d-f)**.

Next, the role of αCOPI-WD40 Arg^57^, Asp^115^, and Tyr^139^ residues in binding the spike hepta-peptide was tested. We generated three single-site mutants of αCOPI-WD40 wherein Arg^57^, Asp^115^, and Tyr^139^ residues were individually mutated to Ala. These mutants were expressed heterologously and purified from Expi293 cells. Analysis by size-exclusion chromatography (SEC) suggested an overall similarity in hydrodynamic radius with the wild-type αCOPI-WD40 domain (**Supplementary Figure S2**). These three mutants were analyzed for binding to the wild-type spike hepta-peptide by BLI assays. None of the three mutants demonstrates any binding to the wild type sequence of the spike hepta-peptide (**Figure 6d-f**). This demonstrated that αCOPI-WD40 residues Arg^57^, Asp^115^, and Tyr^139^ are individually critical for binding the spike hepta-peptide. Disruption of even one of these interactions is likely sufficient to destabilize the spike-COPI complex.

### Structural basis of conformational changes in an αCOPI-WD40 mutant

The results of αCOPI-WD40 Arg^57^, Asp^115^, and Tyr^139^ mutagenesis led us to ask if the loss of binding to the spike hepta-peptide was due to disruption of a single critical interaction or due to larger alterations in the protein structure. This is because the surface of αCOPI-WD40 contains an intricate network of residues with charged and polar side-chains. Mis-sense mutations that alter charge balance could modify αCOPI-WD40 conformations. To address this question, we crystallized αCOPI-WD40 Arg^57^→Ala and Tyr^139^→Ala mutants. The αCOPI-WD40 Asp^115^→Ala mutant did not yield crystals in the conditions we tested. The crystal structures of αCOPI-WD40 Arg^57^→Ala and Tyr^139^→Ala mutants were determined by X-ray diffraction to a resolution of 1.2Å and 1.5Å, respectively (**Table T1**).

The αCOPI-WD40 Arg^57^→Ala mutant structure demonstrated novel structural alterations that had previously not been reported in the crystal structures of wild-type αCOPI-WD40 or the related ꞵ’COPI-WD40 (**Supplementary Figure S4a**). The mutation of Arg^57^ to an Ala residue generated a cavity in the spike hepta-peptide binding site. This change led to a 62° rotation of a nearby Tyr^97^ residue side chain into the newly generated cavity in αCOPI-WD40 (**Supplementary Figure S4b**). In parallel, the residue Asp^73^ underwent a substantial conformational change. This residue interacts with the side chain of Arg^57^ in the wild-type αCOPI-WD40 structure. However, the loss of stabilizing interactions from the Arg^57^ side chain and the reorientation of Tyr^97^ caused the Asp^73^ side chain to rotate away by 73° from its initial position (**Supplementary Figure S4b**). These conformational changes are accompanied by a 1.1Å and 0.8Å movement of Tyr^139^ and His^31^ side chains respectively closer towards the spike hepta-peptide as inferred from our *in silico* model. In contrast, the side chain of Lys^15^ moves 1.7Å away from the inferred hepta-peptide position. As such, the binding site and its vicinity demonstrate a substantially modified interaction network in Arg^57^→Ala mutant.

Next, we asked if Arg^57^→Ala mutation and the associated rotameric changes caused any main chain reorganization in αCOPI-WD40. To obtain a global overview of changes in the main chain geometry, the differences in Ramachandran angles were calculated between corresponding residues in the wild type and Arg^57^→Ala crystal structures (**Supplementary Figure S4c**). The top peak in this difference Ramachandran plot, i.e., peak 1, in this analysis corresponds to a substantial main chain twist at Gly^72^, which is near the Arg^57^→Ala mutation site. This conformational change is associated with the Asp^73^ side chain rotation and repositioning of the main chain atoms from Gly^72^ to Val^77^, which are pushed away from the domain core consistent with the reorientation of the Asp^73^ side chain (**Supplementary Figure S4d**). Peaks 2 and 4 in this difference Ramachandran plot correspond to changes in surface loops that are 19Å and 25Å from the mutation site and are likely due to crystal contacts. Peaks 3 and 6 correspond to main chain rearrangement in the mutation site, i.e., Arg^57^→Ala. This is likely a combination of the mutation and modifications to side chain rearrangements in the neighborhood of Ala^57^ described above. Peak 5 is associated with a surface loop 31Å from the mutation site. This loop demonstrates weak electron density and is only partly ordered. Hence, three of the top six peaks, i.e., 1, 3, and 6, in this analysis are associated with considerable rearrangement of the αCOPI-WD40 surface upon the mutation of basic Arg^57^ to neutral Ala including the site for spike hepta-peptide binding.

In contrast to the Arg^57^→Ala substitution, the crystal structure of αCOPI-WD40 Tyr^139^→Ala mutant demonstrated no substantial changes as compared to the wild-type structure (**Supplementary Figure S4c, e**). No major rearrangements of side chains or main chains were observed. This lack of conformational rearrangement contrasts with the structural changes in the αCOPI-WD40 Arg^57^ →Ala mutant structure. It is likely that the electroneutral change from Tyr^139^ to Ala does not perturb the local electrostatic surface sufficiently to alter protein conformation. Hence, disruption of spike hepta-peptide binding in this Tyr^139^→Ala mutant is due to the loss of a single side chain hydroxyl group. Collectively, the crystal structures of αCOPI-WD40 Arg^57^→Ala and Tyr^139^→Ala mutants reveal distinct and contrasting structural principles by which spike hepta-peptide binding is disrupted.

An analysis of crystal packing in the αCOPI-WD40 structures reported here showed that the peptide binding site residues are in contact with symmetry related chains. We subsequently asked if distinct crystal packing may have contributed to these different structural consequences of Arg^57^→Ala and Tyr^139^→Ala mutations. However, similar crystal packing interactions are provided by residues His^267^ and Lys^309^ from a symmetry-related αCOPI-WD40 chain to the spike hepta-peptide binding site. As such, conformational differences in these two αCOPI-WD40 mutants are due to altered interaction chemistry in the hepta-peptide binding site.

### Conservation of αCOPI-WD40 residues critical for spike hepta-peptide binding

The mutagenesis, BLI, and crystallographic analyses described here are focused on αCOPI. Hence, we asked if αCOPI Arg^57^, Asp^115^, and Tyr^139^ residues are conserved in bats, pangolins, camels, and humans, which have been implicated as zoonotic reservoirs and hosts for β-, coronaviruses ^40-44^. Overall, αCOPI is 96.5-99.8% identical in these multicellular higher organisms (**Supplementary Table T4**). In contrast, αCOPI conservation is relatively moderate between these organisms and yeast at 46.8-47.1% sequence identity. An analysis of αCOPI sequence conservation across 150 species using the CONSURF server ^45^ demonstrated that Arg^57^ and Asp^115^ are completely conserved whereas Tyr^139^ is replaced by Phe or Trp in 5.3% and 0.7% of the sequences, respectively (**Supplementary Data**). Importantly, all these three αCOPI residues are found to be 100% identical in yeast, bat, pangolin, camel, and human αCOPI. This suggests an evolutionary pressure on these three residues in binding dibasic motifs in host proteins, which is exploited by the sarbecovirus spike to hijack the host COPI machinery. This conservation of the three αCOPI residues extends to chicken, which is has been suggested to be a host for γ- and δ-coronaviruses ^31^. αCOPI residues such as Lys^15^ and His^31^ that are suggested to be involved in spike hepta-peptide binding by our *in silico* analysis demonstrate complete conservation. Interestingly, this conservation extends to *S. cerevisiae* β’COPI wherein αCOPI residues Arg^57^ and Asp^115^ are replaced by Arg^59^ and Asp^117^ in β’COPI, respectively. However, αCOPI Tyr^139^ is semi-conserved and is replaced by Phe^142^ in β’COPI.

## Discussion

Once the sarbecovirus spike is delivered from ER to Golgi, its trafficking to the progeny virus assembly site consists of three distinct steps, i.e., spike-COPI binding in donor membranes such as cis-Golgi, inter-organelle trafficking, and dissociation of spike-COPI at the destination, which is ERGIC ^8^. This trafficking pathway can be disrupted by either weakening of spike-COPI binding leading to premature complex dissociation or enhanced stability of this complex, which interferes with spike release. This is supported by recent cellular imaging and biochemical analysis of the SARS-CoV-2 spike protein ^15^. Hence, elucidating the determinants of spike-COPI interactions is fundamental to understanding sarbecovirus assembly.

Employing a spike hepta-peptide and a purified aCOPI-WD40 domain, the present investigation expounds on the biophysical and structural bases of spike-COPI interactions. We demonstrate that direct binding of purified αCOPI-WD40 domain to the SARS-CoV-2 spike hepta-peptide is modulated by an extended coatomer binding motif that stretches beyond the spike K-x-H residues. Our data show that residues such as acidic Glu in the C-terminal position in the spike likely interact with complementary charged basic residues in αCOPI-WD40. This interaction strengthens spike binding to the host αCOPI. This analysis is consistent with a recent preprint that shows a key role of this SARS-CoV-2 spike C-terminal position in pull-down assays of the spike cytosolic domain with COPI subunits ^30^. A second cellular investigation has recently shown that the inferred stabilization of the spike-COPI complex by a Lys^1269^-x-His^1271^→Lys^1269^-x-Lys^1271^ spike mutation has dramatic effects on SARS-CoV-2 spike processing and trafficking (Jennings et al., 2021). This functional analysis suggests a key role of spike-COPI complex dissociation in modulating spike trafficking and function. Hence, it is likely that residues that strengthen spike-COPI complex stability beyond that from wild type interactions are avoided in the spike C-terminus. This includes acidic Glu and unbranched Ala residue that stabilize the αCOPI-WD40 domain as demonstrated in the present investigation. Interestingly, our analysis of the human membrane proteome suggests that the occurrence of a charged residue such as Glu and β-branched residues is a high probability event at the C-terminus of dibasic motifs. This raises an intriguing question of whether such charged residues present a structural and biophysical disadvantage to spike-COPI interactions, and hence, are selected against in sarbecoviruses. Such a highly stabilized complex may not undergo dissociation in ERGIC to release the spike for processing and downstream virion assembly. Interestingly, acidic residues are absent from the C-terminus of coronavirus spike proteins that have a dibasic motif for COPI dependent trafficking. We note that complex formation of Glu^1273^ or Gln^1283^ containing spike hepta-peptide is substantially slower than in the wild type. Even Arg^1273^, which lacks charge complementarity with αCOPI-WD40 residues shows slower association kinetics. Glu, Gln, and Arg have long side chains unlike Thr^1273^, which is suggestive of a role of side chain size in modulating interactions with COPI.

On the coronavirus side, the spike dibasic motif and adjacent residues demonstrate complete conservation in β-coronavirus isolates from bats, pangolins, civets, camels, and humans ^33–38,46^. However, variations in K-x-H motif as well the absence of a dibasic motif in the spike in animal isolates of coronaviruses suggest that this mechanism of direct interactions between COPI and the spike may not be universal. It has been suggested that dibasic motifs in coronavirus proteins other than the spike may be involved in modulating COPI dependent trafficking by oligomerization with the spike protein ^9^. An example is bovine coronavirus, which demonstrates a dibasic motif in the enzyme hemagglutinin esterase but not in the spike protein ^47^. The interaction of this enzyme with the spike protein ^48^ offers a potential route for COPI dependent trafficking of the spike.

our investigation identifies three αCOPI residues Arg^57^, Asp^115^, and Tyr^139^, as essential for spike hepta-peptide binding. These αCOPI residues are completely conserved across organisms associated with β-coronavirus infections such as bats, pangolins, camels, and humans and in chicken, which is infected by γ- and δ-coronaviruses. This suggests a likely conserved mechanism for COPI dependent spike trafficking. Interestingly, the critical αCOPI residues identified in the present analysis are broadly consistent with a prior genetic and biophysical study that implicated αCOPI Arg^57^ and Lys^15^, and β’COPI Arg^59^ and Asp^117^ (equivalent to αCOPI Asp^115^) as critical for dibasic motif binding, retrograde trafficking, and growth of yeast cells ^27^.

Building on this prior investigation, our crystallographic analysis of αCOPI Arg^57^→Ala and Tyr ^139^→Ala mutants presents two complementary structural results to substantially advance the understanding of how these residues are critical for COPI architecture. The αCOPI Arg^57^→Ala mutant demonstrates a rearrangement of the spike hepta-peptide binding site and of neighboring residues whereas the Tyr^139^→Ala mutant structure is largely similar to the wild type αCOPI structure. Yet, both mutants demonstrate the same functional outcome, i.e., loss of spike hepta-peptide binding. Given this structural sensitivity of αCOPI, and presumably β’COPI, to changes in electrostatics, this raises an interesting question about the structural basis of how mutations in these subunits alter normal retrograde trafficking. It is relevant to note that the *COPA* gene, which encodes the human homolog of αCOPI, has been implicated in a set of clinical disorders collectively known as the COPA syndrome ^49,50^. Here, mis-sense mutations including ones that modify side chain charge in the WD40 domains compromise COPA protein function in retrograde trafficking ^49^. Based on data presented here, it would be of interest to investigate the structural basis of this dysfunction to gain deeper insights into COPI biology.

In conclusion, our present analysis and supporting prior investigations demonstrate that the extended dibasic motif in the sarbecovirus spike functions as an effective tool to hijack the COPI complex involved in retrograde trafficking. In broader terms, our structural analysis provides a basis to further investigate the structural and functional consequences of αCOPI and β’COPI mutations in disrupting retrograde trafficking.

## Methods

### Protein and peptide production

The *S. pombe* αCOPI-WD40 domain was synthesized by TOPGENE and cloned in pcDNA3.1(+) with a C-terminal strep-tag for affinity purification. Five mutations (Leu^181^→Lys, Leu^185^→Lys, Ile^192^→Lys, Leu^196^→Lys and Phe^197^→Lys) were incorporated in the gene to improve solubility as suggested previously ^28^. Expression was performed in Expi293 mammalian cells using the Thermo Fisher ExpiFectamine expression kit. Protein purification was performed by affinity chromatography of the clarified cellular lysate followed by SEC in a Superdex 75 chromatography column. Arg^57^→Ala, Asp^115^→Ala, and Tyr^139^→Ala mutants of αCOPI-WD40 domain were expressed and purified as described for the wild type protein. The purified αCOPI-WD40 domain in 150 mM NaCl, 5 mM dithiotreitol (DTT), 10% glycerol, and either 20 mM Tris-HCl (pH 7.5) or 50 mM MES-NaOH (pH 6.5) was flash-frozen in liquid nitrogen until further experimentation. All mutations in pcDNA3.1(+)-αCOPI-WD40 were made GenScript.

β’COPI-WD40 (residues 1-301) from *S. cerevisiae* was cloned in pSUMO vector with an N-terminal strep-tag, a Hisx8 tag, and a Ulp1 protease cleavage site, and expressed overnight in *E. coli* pLysS cells at 18°C. This fusion protein was purified by affinity chromatography and SEC in 150 mM NaCl, 5 mM dithiotreitol (DTT), 10% glycerol, and 20 mM Tris-HCl (pH 7.5) followed by overnight digestion with Ulp1 protease. The digested β’COPI-WD40 domain was subjected to negative purification by Ni-NTA and SEC and was flash-frozen in liquid nitrogen until further experimentation.

Peptide synthesis was performed by Biomatik (USA) with an N-terminal biotin tag and a (PEG)_4_ linker between the tag and the peptide. No modification was performed at the C-terminus of the peptides thereby leaving a free terminal carboxylate group.

### BLI assay

Biotinylated spike hepta-peptides were tethered to streptavidin (SA) biosensors (FortéBio) in a 96-well plate format. Purified αCOPI-WD40 domain was provided as the analyte. Kinetics measurements for determination of binding affinity were performed on an Octet RED96 system (FortéBio). Data acquisition was carried out using the Data Acquisition 11.1 suite. Briefly, SA biosensors were hydrated in 200 μL of kinetics buffer (20mM Tris-HCl (pH 7.5) or 50 mM MES-NaOH (pH 6.5), 150 mM NaCl, 5 mM DTT, 10% glycerol, 0.2 mg/ml bovine serum albumin (BSA), and 0.002% Tween 20) for 10 minutes prior to binding. The spike hepta-peptide (5 μg/ml) was loaded on the biosensors for 15 seconds. A baseline was established by rinsing the biosensor tips in the kinetic buffer for 30 seconds. This was followed by association with αCOP in varying concentrations over 60 seconds and dissociation in the baseline well for 90 seconds. A temperature of 25°C and a shake speed of 1000 rpm was maintained during acquisition. All experiments were carried out in triplicates. A new sensor was used for each replicate. Data processing and analysis were performed in the FortéBio Data Analysis 11.1 software suite. Raw data was subtracted from the 0 μM αCOP signal as a reference. The baseline step immediately before the association step was used for the alignment of the y-axis. An inter-step correction between the association and dissociation steps was performed. Reference subtracted curves were processed with the Savitzky-Golay filtering method and subjected to global fitting using a 1:1 binding model. All fits to BLI data had R^2^ value (goodness of fit) >0.9.

### Crystallization and structure determination

Purified αCOPI-WD40 domain was concentrated to 2 mg/ml in 20 mM Tris-HCl (pH 7.5), 150 mM NaCl, 5 mM DTT and 10% glycerol buffer. Crystal trays were set up with the hanging drop vapor diffusion method with 0.5 μL of αCOPI-WD40 mixed with an equal volume of reservoir buffer. Crystals were within 48 hours at 22°C in 20% PEG3350 and 0.25 M sodium citrate tribasic dihydrate. Crystals were cryo-protected in mother liquor supplemented with 20% ethylene glycol and flash-frozen in liquid nitrogen. Purified Arg^57^→Ala and Tyr^139^→Ala mutants of αCOP were concentrated to ~2.2 mg/ml and crystallized as described for the wild type protein. Crystals for Arg^57^→Ala were obtained in 22% PEG3350 and 0.2 M trisodium citrate and for Tyr^139^→Ala in 18% PEG3350 and 0.2M potassium-sodium tartrate. The crystals for Arg^57^→Ala and Tyr^139^→Ala αCOPI-WD40 mutants were cryoprotected in 20% glycerol and 20% ethylene glycol, respectively. X-ray diffraction data for wild-type αCOPI-WD40 was collected at the beamline GM/CA 23-ID-D of the Advanced Photon Source at the Argonne National Laboratory and at the National Synchrotron Light Source II (NSLS II) beamline 17-ID-1 AMX at the Brookhaven National Laboratory for the mutants. The X-ray diffraction data for the wild type protein crystals was indexed, integrated and scaled using HKL3000 ^51^ whereas those for the mutants were processed using XDS ^52^ as part of the data acquisition and processing pipeline at the beamline. The data processing statistics are given in **Supplementary Table T1**. The scaled data were merged using AIMLESS in CCP4 suite ^53^. Molecular replacement was performed in Phenix using a previously determined αCOPI-WD40 domain structure (PDB ID 4J87) as the search model ^54,55^. Iterative model building and refinement were performed in Phenix.refine ^56^ and Coot ^57^. Figures were generated in PyMol. Part of the software used here was curated by SBGrid ^58^.

### Analysis of Ramachandran angles

The crystal structures of wild type, Arg^57^→Ala, and Tyr^139^→Ala αCOPI-WD40 were analyzed in Molprobity ^59^. For each structure pair, i.e., wild type with Arg^57^→Ala or wild type with Tyr^139^→Ala, per residue difference in Ramachandran angles was determined using equation [1]-

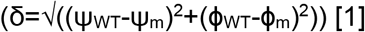

Here, Ramachandran angles for wild type and mutant structures are represented as (ψ_WT_, ϕ_WT_) and (ψ_m_, ϕ_m_) respectively. Each structure-pair was superimposed in PyMol and inspected to ensure consistency with the results of the Ramachandran angle analysis. We identified three instances of surface exposed residues (Asp^96^, Asn^257^ in the wild type coordinates, and Ser^11^ in Arg^57^→Ala coordinates) where the structures were highly similar between corresponding main chain atoms in the wild type and mutant but the sign of a dihedral angle close to 180° had been flipped. The Cα rmsd of short penta-residue stretches of the polypeptide chain centered at each of these residues was 0.14Å, 0.06Å, and 0.18Å, respectively. The signs of the Ramachandran angles for these residues were corrected manually.

### *In silico* analysis of sarbecovirus spike hepta-peptide with αCOPI-WD40

Structural modeling of the SARS-CoV-2 spike C-terminus peptide (sequence: GVKLHYT) in complex with the αCOPI-WD40 domain was performed using homology modeling in Modeller ^60^ and the structure of the αCOP-WD40 complexed with Emp47p peptide (PDB ID 4J8B) as a template. Prior to computational mutagenesis, models were processed with FastRelax ^61^ in Rosetta (v. 3.5), with backbone and side chain atoms constrained to the input coordinates. The command line parameter settings for FastRelax execution (“relax” executable) used were:

-relax:constrain_relax_to_start_coords

-relax:coord_constrain_sidechains

-relax:ramp_constraints false

-ex1

-ex2

-use_input_sc

-correct

-no_his_his_pairE

-no_optH false

-flip_HNQ

-nstruct 1

Computational mutagenesis simulations to predict effects on binding affinities (ΔΔGs) for point substitutions were performed using a previously described protocol implemented in Rosetta (v. 2.3) ^62^. Default parameters were used, with the exception of extra rotamers allowed during packing of modeled side chains, specified by command line parameters:

-extrachi_cutoff 1 -ex1 -ex2 -ex3

### Sequence analysis of dibasic motifs in the human membrane proteome

UNIPROT identifiers of secreted and membrane-bound human proteins, as well as secreted/membrane-bound protein isoforms, were downloaded from the Human Protein Atlas (http://www.proteinatlas.org) ^63^. The corresponding protein sequences were obtained from UNIPROT, leading to approximately 6800 sequences, which were parsed using an in-house Perl script to identify C-terminal motif residues. Of these, 119 sequences that demonstrated a C-terminal dibasic motif were analyzed further for amino acid propensities in the dibasic motif and neighboring residues.

### Accession numbers of protein sequences used in this investigation

Spike protein (bat coronavirus CDPHE15/USA/2006, S5YNL4; bat coronavirus HKU10, K4JZD2; bat *Rhinolopus* alpha-coronavirus/HuB2013, A0A0U1WJW2; human coronavirus 229E, P15423; rodent coronavirus, A0A2H4MWY1; mink coronavirus strain WD1127, D9J1Z4; bat coronavirus HKU8, B1PHK2; bat alpha-coronavirus SAX2011, A0A0U1WHD7; bat alpha-coronavirus SC2013, A0A0U1UZD0; porcine epidemic diarrhea virus strain CV777, Q91AV1; bat coronavirus 512/2005, Q0Q466; bat coronavirus HKU2, A8JP08; human coronavirus NL63, Q6Q1S2; Bat alphacoronavirus, A0A4Y6A7L9; human coronavirus OC43, P36334; beta-coronavirus HKU24, A0A0A7UZR7; human coronavirus HKU1 isolate N5, Q0ZME7; murine coronavirus strain A59, P11224; bat coronavirus/Zhejiang2013, A0A088DJY6; hedgehog coronavirus 1, A0A4D6G1A4; Middle East respiratory syndrome-related coronavirus isolate United Kingdom/H123990006/2012, K9N5Q8; bat coronavirus HKU5, A3EXD0; bat coronavirus HKU4, A3EX94; Rousettus bat coronavirus, A0A1B3Q5V4; bat coronavirus HKU9, A3EXG6; SARS-CoV-2, P0DTC2; SARS-CoV, P59594; avian infectious bronchitis virus strain Beaudette, P11223; bulbul coronavirus HKU11-934, B6VDW0; common moorhen coronavirus HKU21, H9BR35; porcine coronavirus HKU15, X2G836; munia coronavirus HKU13-3514, B6VDY7; night heron coronavirus HKU19, H9BR17; bat coronavirus RaTG13, A0A6B9WHD3; Middle East respiratory syndrome coronavirus isolate Camel/Qatar_2_2014, KJ650098.1; Civet SARS-CoV SZ3/2003, AY304486.1; Pangolin coronavirus cDNA31-S, MT799526.1; bat coronavirus 279/2005, Q0Q475; bat coronavirus Rp3/2004, Q3I5J5; bat SARS-like coronavirus WIV1, AGZ48828.1; porcine transmissible gastroenteritis coronavirus strain Purdue, P07946; bovine coronavirus strain Mebus, P15777), αCOPI (bat, XP_032949522.1; camel, XP_031291824.1; chicken, H9L3L2; civet, XP_036778629.1; human, P53621; yeast, NP_595279.1), and β’COPI (yeast, Q96WV5). The phylogenetic analysis of these sequences was performed in PhyML 3.0 ^64^. A list of αCOPI sequences and their multiple sequence alignment from the CONSURF server ^45^ is given in **Supplementary Data**.

### Statistics and Reproducibility

All BLI experiments were performed in independent triplicates. A statistical correlation coefficient (CC1/2) was used for crystallographic resolution estimation between half datasets. 5% of the crystallographic reflections were omitted from refinement to calculate R_free_ and to avoid over-fitting.

## Acknowledgements

We thank Prof. David Owen (University of Cambridge, Cambridge, England) for critical comments, Dr. Jonathan Goldberg (Memorial Sloan Kettering Cancer Center, New York City, USA), Dr. Elena Goldberg (Memorial Sloan Kettering Cancer Center, New York City, USA), and Dr. Lauren Jackson (Vanderbilt University, Nashville USA) for advice on protein purification, Dr. Travis Gallagher (National Institute of Standards and Technology, Rockville USA) for advice on crystallographic data collection, Dr. Andrey Galkin (University of Maryland Baltimore USA) for advice on BLI assays, and Dr. Jean Jakoncic (NSLSII beamline 17-ID-1 AMX), Dr. Stephan Corcoran (APS beamline 23-ID-D), and Dr. Darren Sherrell (APS beamline 19-ID) for advice during crystallographic data collection and processing, and Ms. Corrinne Wilson for assistance in manuscript preparation. SSH acknowledges support from the University of Maryland School of Medicine, University of Maryland MPower, and MPower COVID-19 Response Fund Award. This article was supported by funds through the Maryland Department of Health’s Cigarette Restitution Fund Program, University of Maryland Marlene and Stewart Greenebaum Comprehensive Cancer Center (National Cancer Institute - Cancer Center Support Grant (CCSG) - P30CA134274), The Holden Comprehensive Cancer Center at The University of Iowa and its National Cancer Institute Award P30CA086862. This research used resources of the Advanced Photon Source, a U.S. Department of Energy (DOE) Office of Science User Facility, operated for the DOE Office of Science by Argonne National Laboratory under Contract No. DE-AC02-06CH11357. GM/CA@APS has been funded by the National Cancer Institute (ACB-12002) and the National Institute of General Medical Sciences (AGM-12006, P30GM138396). Access to Sector 84 laboratories at the Advanced Protein Characterization Facility (APCF) was possible through funding provided by NIH grant GM115586 and DOE contract DE-AC02-06CH11357. This research used resources AMX of the National Synchrotron Light Source II, a U.S. DOE Office of Science User Facility operated for the DOE Office of Science by Brookhaven National Laboratory under contract no. DE-SC0012704.

## Author contributions

DD: Data acquisition, analysis, interpretation, and manuscript preparation.

SS: Data acquisition, analysis, interpretation, and manuscript preparation.

SK: Data acquisition, analysis, and interpretation.

MM: Data acquisition.

NJS: Data acquisition, analysis, interpretation, and manuscript preparation.

LG: Data acquisition, analysis, interpretation, and manuscript preparation.

BGP: Data acquisition, analysis, interpretation, and manuscript preparation.

SSH: Conception, data acquisition, analysis, interpretation, and manuscript preparation.

## Competing interests

The authors declare no competing interests.

## Data availability

Coordinates for the crystal structures have been deposited in the Protein Data Bank with IDs: 7S22 (wild type), 7S16 (Arg^57^→Ala), and 7S23 (Tyr^139^→Ala). The plasmids for constructs pSUMO-β′-COPI-WD40, pcDNA3.1(+)-αCOPI-WD40, pcDNA3.1(+)-αCOPI-WD40 mutant Arg^57^→Ala, pcDNA3.1(+)-αCOPI-WD40 mutant Asp^115^→Ala, and pcDNA3.1(+)-αCOPI-WD40 mutant Tyr^139^→Ala will be deposited in www.addgene.org.

## Notes

### Competing Interest Statement

The authors have declared no competing interest.

## References

1 Stadler, K. et al. SARS-beginning to understand a new virus. Nat Rev Microbiol 1, 209–218, doi:10.1038/nrmicro775 (2003).

2 Ahmed, A. E. The predictors of 3- and 30-day mortality in 660 MERS-CoV patients. BMC Infect Dis 17, 615, doi:10.1186/s12879-017-2712-2 (2017).

3 Zaki, A. M., van Boheemen, S., Bestebroer, T. M., Osterhaus, A. D. & Fouchier, R. A. Isolation of a novel coronavirus from a man with pneumonia in Saudi Arabia. N Engl J Med 367, 1814–1820, doi:10.1056/NEJMoa1211721 (2012).

4 Polack, F. P. et al. Safety and efficacy of the BNT162b2 mRNA COVID-19 vaccine. N Engl J Med 383, 2603–2615, doi:10.1056/NEJMoa2034577 (2020).

5 USFDA. U.S. Food and Drug Administration. Moderna COVID-19 vaccine [FDA briefing document]. Silver Spring, MD: U.S. Food and Drug Administration, Vaccines and Related Biological Products Advisory Committee; 2020., doi:https://www.fda.gov/media/144434/download (2020).

6 Beniac, D. R., Andonov, A., Grudeski, E. & Booth, T. F. Architecture of the SARS coronavirus prefusion spike. Nat Struct Mol Biol 13, 751–752, doi:10.1038/nsmb1123 (2006).

7 Song, H. C. et al. Synthesis and characterization of a native, oligomeric form of recombinant severe acute respiratory syndrome coronavirus spike glycoprotein. J Virol 78, 10328–10335, doi:10.1128/JVI.78.19.10328-10335.2004 (2004).

8 Klumperman, J. et al. Coronavirus M proteins accumulate in the Golgi complex beyond the site of virion budding. J Virol 68, 6523–6534, doi:10.1128/JVI.68.10.6523-6534.1994 (1994).

9 Lontok, E., Corse, E. & Machamer, C. E. Intracellular targeting signals contribute to localization of coronavirus spike proteins near the virus assembly site. J Virol 78, 5913–5922, doi:10.1128/JVI.78.11.5913-5922.2004 (2004).

10 McBride, C. E., Li, J. & Machamer, C. E. The cytoplasmic tail of the severe acute respiratory syndrome coronavirus spike protein contains a novel endoplasmic reticulum retrieval signal that binds COPI and promotes interaction with membrane protein. J Virol 81, 2418–2428, doi:10.1128/JVI.02146-06 (2007).

11 Cosson, P. & Letourneur, F. Coatomer interaction with di-lysine endoplasmic reticulum retention motifs. Science 263, 1629–1631, doi:10.1126/science.8128252 (1994).

12 Gaynor, E. C., te Heesen, S., Graham, T. R., Aebi, M. & Emr, S. D. Signal-mediated retrieval of a membrane protein from the Golgi to the ER in yeast. J Cell Biol 127, 653–665, doi:10.1083/jcb.127.3.653 (1994).

13 Jackson, M. R., Nilsson, T. & Peterson, P. A. Retrieval of transmembrane proteins to the endoplasmic reticulum. J Cell Biol 121, 317–333, doi:10.1083/jcb.121.2.317 (1993).

14 Townsley, F. M. & Pelham, H. R. The KKXX signal mediates retrieval of membrane proteins from the Golgi to the ER in yeast. Eur J Cell Biol 64, 211–216 (1994).

15 Jennings, B. C., Kornfeld, S. & Doray, B. A weak COPI binding motif in the cytoplasmic tail of SARS-CoV-2 spike glycoprotein is necessary for its cleavage, glycosylation, and localization. FEBS Lett 595, 1758–1767, doi:10.1002/1873-3468.14109 (2021).

16 Letourneur, F. et al. Coatomer is essential for retrieval of dilysine-tagged proteins to the endoplasmic reticulum. Cell 79, 1199–1207, doi:10.1016/0092-8674(94)90011-6 (1994).

17 Dodonova, S. O. et al. A structure of the COPI coat and the role of coat proteins in membrane vesicle assembly. Science 349, 195–198, doi:10.1126/science.aab1121 (2015).

18 Duden, R., Griffiths, G., Frank, R., Argos, P. & Kreis, T. E. β-COP, a 110 Kd protein associated with non-clathrin-coated vesicles and the Golgi complex, shows homology to β-adaptin. Cell 64, 649–665, doi:10.1016/0092-8674(91)90248-w (1991).

19 Hara-Kuge, S. et al. En bloc incorporation of coatomer subunits during the assembly of COP-coated vesicles. J Cell Biol 124, 883–892, doi:10.1083/jcb.124.6.883 (1994).

20 Harrison-Lavoie, K. J. et al. A 102 KDa subunit of a Golgi-associated particle has homology to beta subunits of trimeric G proteins. EMBO J 12, 2847–2853 (1993).

21 Malhotra, V., Serafini, T., Orci, L., Shepherd, J. C. & Rothman, J. E. Purification of a novel class of coated vesicles mediating biosynthetic protein transport through the Golgi stack. Cell 58, 329–336, doi:10.1016/0092-8674(89)90847-7 (1989).

22 Serafini, T. et al. A coat subunit of Golgi-derived non-clathrin-coated vesicles with homology to the clathrin-coated vesicle coat protein β-adaptin. Nature 349, 215–220, doi:10.1038/349215a0 (1991).

23 Waters, M. G., Serafini, T. & Rothman, J. E. ’Coatomer’: A cytosolic protein complex containing subunits of non-clathrin-coated Golgi transport vesicles. Nature 349, 248–251, doi:10.1038/349248a0 (1991).

24 Stenbeck, G. et al. beta’-COP, a novel subunit of coatomer. EMBO J 12, 2841–2845 (1993).

25 Eugster, A., Frigerio, G., Dale, M. & Duden, R. The α- and β′-COP WD40 domains mediate cargo-selective interactions with distinct di-lysine motifs. Mol Biol Cell 15, 1011–1023, doi:10.1091/mbc.e03-10-0724 (2004).

26 Fiedler, K., Veit, M., Stamnes, M. A. & Rothman, J. E. Bimodal interaction of coatomer with the p24 family of putative cargo receptors. Science 273, 1396–1399, doi:10.1126/science.273.5280.1396 (1996).

27 Jackson, L. P. et al. Molecular basis for recognition of dilysine trafficking motifs by COPI. Dev Cell 23, 1255–1262, doi:10.1016/j.devcel.2012.10.017 (2012).

28 Ma, W. & Goldberg, J. Rules for the recognition of dilysine retrieval motifs by coatomer. EMBO J 32, 926–937, doi:10.1038/emboj.2013.41 (2013).

29 Schroder-Kohne, S., Letourneur, F. & Riezman, H. α-COP can discriminate between distinct, functional di-lysine signals in vitro and regulates access into retrograde transport. J Cell Sci 111 (Pt 23), 3459–3470 (1998).

30 Cattin-Ortolá, J. et al. Sequences in the cytoplasmic tail of SARS-CoV-2 spike facilitate expression at the cell surface and syncytia formation. Nature Communications 12, 5333 (2021).

31 Woo, P. C. et al. Discovery of seven novel mammalian and avian coronaviruses in the genus deltacoronavirus supports bat coronaviruses as the gene source of alphacoronavirus and betacoronavirus and avian coronaviruses as the gene source of gammacoronavirus and deltacoronavirus. J Virol 86, 3995–4008, doi:10.1128/JVI.06540-11 (2012).

32 Drexler, J. F., Corman, V. M. & Drosten, C. Ecology, evolution and classification of bat coronaviruses in the aftermath of SARS. Antiviral Res 101, 45–56, doi:10.1016/j.antiviral.2013.10.013 (2014).

33 Li, W. et al. Bats are natural reservoirs of SARS-like coronaviruses. Science 310, 676–679, doi:10.1126/science.1118391 (2005).

34 Ge, X. Y. et al. Isolation and characterization of a bat SARS-like coronavirus that uses the ACE2 receptor. Nature 503, 535–538, doi:10.1038/nature12711 (2013).

35 Guan, Y. et al. Isolation and characterization of viruses related to the SARS coronavirus from animals in southern China. Science 302, 276–278, doi:10.1126/science.1087139 (2003).

36 Lau, S. K. et al. Severe acute respiratory syndrome coronavirus-like virus in Chinese horseshoe bats. Proc Natl Acad Sci U S A 102, 14040–14045, doi:10.1073/pnas.0506735102 (2005).

37 Tang, X. C. et al. Prevalence and genetic diversity of coronaviruses in bats from China. J Virol 80, 7481–7490, doi:10.1128/JVI.00697-06 (2006).

38 Xiao, K. et al. Isolation of SARS-CoV-2-related coronavirus from Malayan pangolins. Nature 583, 286–289, doi:10.1038/s41586-020-2313-x (2020).

39 WHO-2021. https://www.who.int/en/activities/tracking-SARS-CoV-2-variants/, accessed on 12/03/2021.

40 Bolles, M., Donaldson, E. & Baric, R. SARS-CoV and emergent coronaviruses: Viral determinants of interspecies transmission. Curr Opin Virol 1, 624–634, doi:10.1016/j.coviro.2011.10.012 (2011).

41 Han, H. J., Yu, H. & Yu, X. J. Evidence for zoonotic origins of Middle East respiratory syndrome coronavirus. J Gen Virol 97, 274–280, doi:10.1099/jgv.0.000342 (2016).

42 Latinne, A. et al. Origin and cross-species transmission of bat coronaviruses in China. Nat Commun 11, 4235, doi:10.1038/s41467-020-17687-3 (2020).

43 Roess, A., Carruth, L., Lahm, S. & Salman, M. Camels, MERS-CoV, and other emerging infections in east Africa. Lancet Infect Dis 16, 14–15, doi:10.1016/S1473-3099(15)00471-5 (2016).

44 Zhang, T., Wu, Q. & Zhang, Z. Probable pangolin origin of SARS-CoV-2 associated with the COVID-19 outbreak. Curr Biol 30, 1346–1351 e1342, doi:10.1016/j.cub.2020.03.022 (2020).

45 Ashkenazy, H. et al. ConSurf 2016: An improved methodology to estimate and visualize evolutionary conservation in macromolecules. Nucleic Acids Res 44, W344–350, doi:10.1093/nar/gkw408 (2016).

46 Raj, V. S. et al. Isolation of MERS coronavirus from a dromedary camel, Qatar, 2014. Emerg Infect Dis 20, 1339–1342, doi:10.3201/eid2008.140663 (2014).

47 Kienzle, T. E., Abraham, S., Hogue, B. G. & Brian, D. A. Structure and orientation of expressed bovine coronavirus hemagglutinin-esterase protein. J Virol 64, 1834–1838, doi:10.1128/JVI.64.4.1834-1838.1990 (1990).

48 Nguyen, V. P. & Hogue, B. G. Protein interactions during coronavirus assembly. J Virol 71, 9278–9284, doi:10.1128/JVI.71.12.9278-9284.1997 (1997).

49 Watkin, L. B. et al. COPA mutations impair ER-Golgi transport and cause hereditary autoimmune-mediated lung disease and arthritis. Nat Genet 47, 654–660, doi:10.1038/ng.3279 (2015).

50 Vece, T. J. et al. COPA syndrome: A novel autosomal dominant immune dysregulatory disease. J Clin Immunol 36, 377–387, doi:10.1007/s10875-016-0271-8 (2016).

51 Minor, W., Cymborowski, M., Otwinowski, Z. & Chruszcz, M. HKL-3000: The integration of data reduction and structure solution-from diffraction images to an initial model in minutes. Acta Crystallogr D Biol Crystallogr 62, 859–866, doi:10.1107/S0907444906019949 (2006).

52 Kabsch, W. XDS. Acta Crystallogr D Biol Crystallogr 66, 125–132, doi:10.1107/S0907444909047337 (2010).

53 Evans, P. R. & Murshudov, G. N. How good are my data and what is the resolution? Acta Crystallogr D Biol Crystallogr 69, 1204–1214, doi:10.1107/S0907444913000061 (2013).

54 McCoy, A. J. et al. Phaser crystallographic software. J Appl Crystallogr 40, 658–674, doi:10.1107/S0021889807021206 (2007).

55 Rossmann, M. G. & Blow, D. M. The detection of sub-units within the crystallographic asymmetric unit. Acta Crystallographica 15, 24–31, doi:doi:10.1107/S0365110X62000067 (1962).

56 Afonine, P. V. et al. Towards automated crystallographic structure refinement with phenix.refine. Acta Crystallogr D Biol Crystallogr 68, 352–367, doi:10.1107/S0907444912001308 (2012).

57 Emsley, P. & Cowtan, K. Coot: Model-building tools for molecular graphics. Acta Crystallogr D Biol Crystallogr 60, 2126–2132, doi:10.1107/S0907444904019158 (2004).

58 Morin, A. et al. Collaboration gets the most out of software. Elife 2, e01456, doi:10.7554/eLife.01456 (2013).

59 Williams, C. J. et al. MolProbity: More and better reference data for improved all-atom structure validation. Protein Sci 27, 293–315, doi:10.1002/pro.3330 (2018).

60 Webb, B. & Sali, A. Protein structure modeling with MODELLER. Methods Mol Biol 1137, 1–15, doi:10.1007/978-1-4939-0366-5_1 (2014).

61 Conway, P., Tyka, M. D., DiMaio, F., Konerding, D. E. & Baker, D. Relaxation of backbone bond geometry improves protein energy landscape modeling. Protein Sci 23, 47–55, doi:10.1002/pro.2389 (2014).

62 Kortemme, T. & Baker, D. A simple physical model for binding energy hot spots in protein-protein complexes. Proc Natl Acad Sci U S A 99, 14116–14121, doi:10.1073/pnas.202485799 (2002).

63 Thul, P. J. et al. A subcellular map of the human proteome. Science 356, doi:10.1126/science.aal3321 (2017).

64 Guindon, S. et al. New algorithms and methods to estimate maximum-likelihood phylogenies: assessing the performance of PhyML 3.0. Syst Biol 59, 307–321, doi:10.1093/sysbio/syq010 (2010).

65 Pierce, B. G. et al. Computational design of the affinity and specificity of a therapeutic T cell receptor. PLoS Comput Biol 10, e1003478, doi:10.1371/journal.pcbi.1003478 (2014).

