## Supplementary Table 1 for "An Extended Motif in the SARS-CoV-2 Spike Modulates Binding and Release of Host Coatomer in Retrograde Trafficking"

**Supplementary Table 1: Crystallographic data and refinement statistics**

| **Structure** | **αCOPI-WD40 WT** | **αCOPI-WD40 $Arg^{57}$→Ala** | **αCOPI-WD40 $Tyr^{139}$→Ala** |
|---|---|---|---|
| PDB ID | 7S22 | 7S16 | 7S23 |
| X-ray Source | 23-ID-D | AMX 17-ID-1 | AMX 17-ID-1 |
| Wavelength (Å) | 1.03 | 0.92 | 0.92 |
| Temperature (K) | 100 | 100 | 100 |
| Space group | $P\ 1\ 2_1\ 1$ | $P\ 1\ 2_1\ 1$ | $P1\ 2_1\ 1$ |
| Unit cell (Å, º) | a= 36.99, b=171.82, c=71.42; α=γ=90, β=99.57 | a=35.87, b=56.71, c=70.55; α=γ=90, β=99.37 | a=37.31, b=171.55, c=71.28; α=γ=90, β=99.71 |
| Resolution (Å) | 44.4-1.8 (1.78-1.75) | 69.6-1.2 (1.26– 1.24) | 171.6 -1.5 (1.52-1.49) |
| $^{a}R_{merge}$ (%) | 9.2 (54.7) | 5.4 (57.9) | 8.6(71.9) |
| <I/σ(I)> | 8.3 (2.2) | 11.5 (1.6) | 5.0 (1.1) |
| CC1/2 (%) | 98.1 (65.3) | 99.8 (66.3) | 99.6 (67.0) |
| No. of reflections | 196291 (10446) | 319698 (6270) | 502939 (25265) |
| No. of unique reflections | 85435 (4461) | 74665(2292) | 140251 (6840) |
| Completeness (%) | 97.0 (95.4) | 94.2 (58.9) | 97.9 (96.8) |
| Redundancy | 2.3 (2.3) | 4.3 (2.7) | 3.6 (3.7) |
| **Refinement Statistics** | | | |
| Resolution (Å) | 44.48-1.75 (1.79-1.75) | 43.97-1.24 (1.26-1.24) | 65.02-1.49 (1.51-1.49) |
| No. of reflections (F>0) used in refinement | 81123 (2645) | 74567 (1653) | 140153 (4335) |
| $^{b}R_{work}$ (%) | 17.3 | 13.5 | 14.0 |
| $^{c}R_{free}$ (%) | 21.4 | 15.9 | 18.4 |
| RMS bond length (Å) | 0.007 | 0.005 | 0.005 |
| RMS bond angle (°) | 0.938 | 0.910 | 0.696 |

| | | | |
|---|---|---|---|
| Overall B value ($Å^2$) | 21.1 | 21.8 | 21.7 |
| **Ramachandran Plot Statistics[d]** | | | |
| Residues | 924 | 321 | 923 |
| Favored (%) | 96.1 | 95.6 | 95.8 |
| Allowed (%) | 3.8 | 4.4 | 4.2 |
| Disallowed (%) | 0.1 | 0.0 | 0.0 |

[a]$R_{merge}$ = [∑h∑i|Ih – Ihi|/∑h∑iIhi] where Ih is the mean of Ihi observations of reflection h. Numbers in parenthesis represent highest resolution shell. [b]$R_{work}$ and [c]$R_{free}$ = ∑||Fobs| - |Fcalc|| / ∑|Fobs| x 100 for 95% of recorded data ($R_{work}$) or 5% data ($R_{free}$). [d]From MolProbity(Williams et al., 2018).
