## Supplementary Table 2 for "An Extended Motif in the SARS-CoV-2 Spike Modulates Binding and Release of Host Coatomer in Retrograde Trafficking"

**Supplementary Table 2: C-terminal sequence of predicted human membrane proteins with K-x-H-x-x or K-x-K-x-x motif**

| UNIPROT ID | Protein Name | C-terminal Sequence |
|---|---|---|
| Q9NVV5 | Androgen-induced gene 1 protein | KPKLE |
| Q9BVK2 | Probable dolichyl pyrophosphate Glc1Man9GlcNAc2 alpha-1,3-glucosyltransferase | KTKKQ |
| Q9NW15 | Anoctamin-10 | KEKAT |
| Q9P241 | Phospholipid-transporting ATPase VD | KGKES |
| O15342 | V-type proton ATPase subunit e 1 | KYHWP |
| Q6UW56 | All-trans retinoic acid-induced differentiation factor | KAKTS |
| Q5SY80 | Cation channel sperm-associated protein subunit epsilon | KRKKN |
| Q96A33 | PAT complex subunit CCDC47 | KVKAM |
| P27544 | Ceramide synthase 1 | KDKRF |
| Q99675 | Cell growth regulator with RING finger domain protein 1 | KPKTL |
| Q9NZ45 | CDGSH iron-sulfur domain-containing protein 1 | KKKET |
| Q8N5K1 | CDGSH iron-sulfur domain-containing protein 2 | KKKEV |
| Q9ULY5 | C-type lectin domain family 4 member E | KGKSL |
| Q6UXB4 | C-type lectin domain family 4 member G | KRHNC |
| P39656 | Dolichyl-diphosphooligosaccharide--protein glycosyltransferase 48 kDa subunit | KEKSD |
| Q6IAN0 | Dehydrogenase/reductase SDR family member 7B | KSKNS |
| Q8NFT8 | Delta and Notch-like epidermal growth factor-related receptor | KTKDL |
| Q15125 | 3-beta-hydroxysteroid-Delta(8),Delta(7)-isomerase | KSKKN |
| Q9BW60 | Elongation of very long chain fatty acids protein 1 | KVKAN |
| Q9HB03 | Elongation of very long chain fatty acids protein 3 | KTKSQ |
| Q9GZR5 | Elongation of very long chain fatty acids protein 4 | KAKGD |
| A1L3X0 | Elongation of very long chain fatty acids protein 7 | KNKDN |
| Q96A26 | Protein FAM162A | KAKTE |
| Q96ND0 | Protein FAM210A | KKKVE |
| Q9NYL4 | Peptidyl-prolyl cis-trans isomerase FKBP11 | KSKKK |
| Q96MZ0 | Ganglioside-induced differentiation-associated protein 1-like 1 | KKKYI |
| Q9P035 | Very-long-chain (3R)-3-hydroxyacyl-CoA dehydratase 3 | KKKIH |
| Q5VWC8 | Very-long-chain (3R)-3-hydroxyacyl-CoA dehydratase 4 | KKKKM |
| Q9HCP6 | Protein-cysteine N-palmitoyltransferase HHAT-like protein | KEKPE |
| Q53GQ0 | Very-long-chain 3-oxoacyl-CoA reductase | KTKKN |
| P37059 | 17-beta-hydroxysteroid dehydrogenase type 2 | KKKAT |
| P14060 | 3 beta-hydroxysteroid dehydrogenase/Delta 5-->4-isomerase type 1 | KSKTQ |
| P26439 | 3 beta-hydroxysteroid dehydrogenase/Delta 5-->4-isomerase type 2 | KSKTQ |
| O15503 | Insulin-induced gene 1 protein | KPHSD |
| Q9Y5U4 | Insulin-induced gene 2 protein | KSHQE |
| Q96N16 | Janus kinase and microtubule-interacting protein 1 | KLKFM |
| Q86W47 | Calcium-activated potassium channel subunit beta-4 | KRKFS |
| Q643R3 | Lysophospholipid acyltransferase LPCAT4 | KQKGD |
| Q6ZNC8 | Lysophospholipid acyltransferase 1 | KRKTD |
| Q96T53 | Ghrelin O-acyltransferase | KHKCN |
| Q8NBP5 | Major facilitator superfamily domain-containing protein 9 | KLKSE |
| Q53F39 | Metallophosphoesterase 1 | KRKTR |
| P39210 | Protein Mpv17 | KAHRL |
| Q969V3 | Nicalin | KAKTQ |
| Q9Y266 | Nuclear migration protein nudC | KAKFN |
| P47890 | Olfactory receptor 1G1 | KIHSP |
| Q96R27 | Olfactory receptor 2M4 | KRKLI |

| | | |
|---|---|---|
| Q8NGI8 | Olfactory receptor 5AN1 | KRKCC |
| A6NM76 | Olfactory receptor 6C76 | KKKKH |
| Q8NGZ6 | Olfactory receptor 6F1 | KWKGK |
| Q9HC56 | Protocadherin-9 | KEHQL |
| O75192 | Peroxisomal membrane protein 11A | KLKTR |
| Q96FM1 | Post-GPI attachment to proteins factor 3 | KFKLD |
| O95427 | GPI ethanolamine phosphate transferase 1 | KSHFM |
| O60486 | Plexin-C1 | KCKWM |
| Q969W9 | Protein TMEPAI | KGHPL |
| Q16799 | Reticulon-1 | KRHAE |
| O75298 | Reticulon-2 | KAKAE |
| O95197 | Reticulon-3 | KKKAE |
| Q9NQC3 | Reticulon-4 | KRKAE |
| Q9NTJ5 | Phosphatidylinositol-3-phosphatase SAC1 | KEKID |
| Q9UI33 | Sodium channel protein type 11 subunit alpha | KVHCD |
| P35498 | Sodium channel protein type 1 subunit alpha | KAKGK |
| Q96BI1 | Solute carrier family 22 member 18 | KDKVR |
| P78381 | UDP-galactose translocator | KVKGS |
| Q9NXE4 | Sphingomyelin phosphodiesterase 4 | KLHQP |
| P43308 | Translocon-associated protein subunit beta | KTKKN |
| Q9P246 | Stromal interaction molecule 2 | KKKSK |
| O15260 | Surfeit locus protein 4 | KKKEW |
| O15533 | Tapasin | KKKAE |
| P57738 | T-cell leukemia translocation-altered gene protein | KTHRE |
| Q6UX40 | Transmembrane protein 107 | KKKPF |
| A0PK00 | Transmembrane protein 120B | KTKQP |
| Q9H6L2 | Transmembrane protein 231 | KEHLS |
| Q53FP2 | Novel acetylcholine receptor chaperone | KVKVS |
| Q5BJD5 | Transmembrane protein 41B | KQKFE |
| Q5BJF2 | Sigma intracellular receptor 2 | KRKKK |
| Q15629 | Translocating chain-associated membrane protein 1 | KEKSS |
| Q8N609 | Translocating chain-associated membrane protein 1-like 1 | KEKSS |
| Q15035 | Translocating chain-associated membrane protein 2 | KLKSP |
| O60858 | E3 ubiquitin-protein ligase TRIM13 | KYKLL |
| Q9HCX4 | Short transient receptor potential channel 7 | KGKDI |
| Q9GZZ9 | Ubiquitin-like modifier-activating enzyme 5 | KMKNM |
| P22309 | UDP-glucuronosyltransferase 1A1 | KSKTH |
| Q9HAW8 | UDP-glucuronosyltransferase 1A10 | KSKTH |
| P35503 | UDP-glucuronosyltransferase 1A3 | KSKTH |
| P22310 | UDP-glucuronosyltransferase 1A4 | KSKTH |
| P35504 | UDP-glucuronosyltransferase 1A5 | KSKTH |
| P19224 | UDP-glucuronosyltransferase 1-6 | KSKTH |
| Q9HAW7 | UDP-glucuronosyltransferase 1A7 | KSKTH |
| Q9HAW9 | UDP-glucuronosyltransferase 1A8 | KSKTH |
| O60656 | UDP-glucuronosyltransferase 1A9 | KSKTH |
| P36537 | UDP-glucuronosyltransferase 2B10 | KGKRD |
| O75310 | UDP-glucuronosyltransferase 2B11 | KGKRD |
| P54855 | UDP-glucuronosyltransferase 2B15 | KKKRD |
| O75795 | UDP-glucuronosyltransferase 2B17 | KKKRD |
| Q9BY64 | UDP-glucuronosyltransferase 2B28 | KGKRD |
| P06133 | UDP-glucuronosyltransferase 2B4 | KGKRD |
| P16662 | UDP-glucuronosyltransferase 2B7 | KGKND |
| Q6NUS8 | UDP-glucuronosyltransferase 3A1 | KVKKT |
| Q3SY77 | UDP-glucuronosyltransferase 3A2 | KVKET |
| Q9BQB6 | Vitamin K epoxide reductase complex subunit 1 | KAKRH |

| | | |
|---|---|---|
| Q8NB15 | Zinc finger protein 511 | KTKQC |
| P58397 | A disintegrin and metalloproteinase with thrombospondin motifs 12 | KSKEL |
| P20851 | C4b-binding protein beta chain | KAKLL |
| P78556 | C-C motif chemokine 20 | KVKNM |
| Q8WUJ3 | Cell migration-inducing and hyaluronan-binding protein | KKKKL |
| O43927 | C-X-C motif chemokine 13 | KRKIP |
| Q30KQ7 | Beta-defensin 113 | KLHQK |
| P0DP74 | Beta-defensin 130A | KGKSP |
| P0DP73 | Beta-defensin 130B | KGKSP |
| Q13609 | Deoxyribonuclease gamma | KSKRS |
| P00740 | Coagulation factor IX | KTKLT |
| P11150 | Hepatic triacylglycerol lipase | KRKIR |
| Q8N1E2 | Lysozyme g-like protein 1 | KRHGF |
| O95631 | Netrin-1 | KCKKA |
| Q9UKZ9 | Procollagen C-endopeptidase enhancer 2 | KNKQC |
| P01270 | Parathyroid hormone | KAKSQ |
| P07225 | Vitamin K-dependent protein S | KTKNS |
