## Supplementary Table 3 for "An Extended Motif in the SARS-CoV-2 Spike Modulates Binding and Release of Host Coatomer in Retrograde Trafficking"

**Supplementary Table 3: Frequency of residues in K-x-H(K)-x-x**

| | Residue position in dibasic motif K-x-H(K)-x-x | | | | |
|---|---|---|---|---|---|
| | Position 1 | Position 2 | Position 3 | Position 4 | Position 5 |
| Amino Acid | | | | | |
| Ala | 0 | 9 | 0 | 8 | 1 |
| Arg | 0 | 11 | 0 | 9 | 3 |
| Asn | 0 | 2 | 0 | 5 | 7 |
| Asp | 0 | 2 | 0 | 3 | 13 |
| Cys | 0 | 2 | 0 | 3 | 3 |
| Glu | 0 | 7 | 0 | 5 | 11 |
| Gln | 0 | 2 | 0 | 6 | 5 |
| Gly | 0 | 9 | 0 | 5 | 0 |
| His | 0 | 1 | 13 | 0 | 10 |
| Ile | 0 | 1 | 0 | 3 | 3 |
| Leu | 0 | 5 | 0 | 6 | 8 |
| Lys | 100 | 13 | 87 | 10 | 5 |
| Met | 0 | 1 | 0 | 0 | 6 |
| Phe | 0 | 1 | 0 | 4 | 3 |
| Pro | 0 | 3 | 0 | 3 | 7 |
| Ser | 0 | 15 | 0 | 11 | 9 |
| Thr | 0 | 1 | 0 | 15 | 5 |
| Trp | 0 | 1 | 0 | 2 | 1 |
| Tyr | 0 | 2 | 0 | 1 | 0 |
| Val | 0 | 7 | 0 | 3 | 1 |
