## Supplementary Table 4 for "An Extended Motif in the SARS-CoV-2 Spike Modulates Binding and Release of Host Coatomer in Retrograde Trafficking"

**Supplementary Table 4: Conservation of αCOPI in coronavirus zoonotic reservoirs and humans[1]**

| Organism | Yeast | Bat | Chicken | Pangolin | Camel | Human |
|---|---|---|---|---|---|---|
| **Yeast** | -- | 47.0/63.4 | 47.6/63.4 | 47.1/63.7 | 47.0/63.5 | 46.8/63.5 |
| **Bat** | -- | -- | 96.7/98.8 | 98.5/99.3 | 98.9/99.4 | 98.7/99.3 |
| **Chicken** | -- | -- | -- | 96.5/98.7 | 96.7/98.9 | 96.7/98.6 |
| **Pangolin** | -- | -- | -- | -- | 99.1/99.8 | 98.5/99.6 |
| **Camel** | -- | -- | -- | -- | -- | 98.9/99.8 |

[1]Conservation is represented as %identity/%similarity.
