## Supplementary Data for "An Extended Motif in the SARS-CoV-2 Spike Modulates Binding and Release of Host Coatomer in Retrograde Trafficking"

**Multiple sequence alignment of αCOPI orthologs from the CONSURF server.**

>Input_pdb_SEQRES_A

MAMLTKFESRSSRAKGVAFH--PTQPWILTSLHNGRIQLWDYRMGTLLD-------RFDG

HDG-PVRGIAFHPT-QPIFVSGGDDYKVNVWNYK--SR-------------KLL-FSLCG

HMDYVRVCTFHHE---YPWILSCSDDQTIRIWNWQ--SRNCIAILTGHSHYVMCAAFHPS

-EDLIVSASLDQTVRVWDI--------S---------GL-RMKN-A-A-P------V---

-----SM---------SK-E--------D-QKAQ-------------AHN-S--------

-ISNDLF----------------GSAD--AIVKFVLEGHDRGVNWCAFHPTLPLILSAGD

DRLVKLWRM------------T----------------ASKAWEVDTCRGHFNNVSCCLF

HPHQ-ELILSASEDKTIRVWDLNRRTAVQTFRR-AN----------DRFWFITVHPKLNL

FAAAHDSGVMVFKLE

>UniRef90_S9Q578_1_327 | Coatomer subunit alpha n=2 Tax=Schizosaccharomyces TaxID=4895 RepID=S9Q578_SCHOY | E_val=1.3e-214

MEMLIKFESKSSRAKGVAFH--PTLPWVLTSLHNGRIQLWDYRMGTLID-------RFDG

HDG-PVRGIAFHPT-QPLFVSGGDDYKVNVWNYK--AK-------------KLL-FSLCG

HMDYVRVCAFHHE---YPWILSCSDDQTIRIWNWQ--SRNCIAILTGHSHYVMCAAFHPS

-EDLIVSASLDQTVRVWDI--------S---------GL-RMKN-A-A-P------V---

-----SM---------TI-E--------E-QLAQ-------------AQN-N--------

-ITNDLF----------------GSTD--AVVKFVLEGHDRGANWCAFHPTLPLILSSGD

DRLIKLWRM------------T----------------ASKAWEVDTCRGHFNNVLCCLF

HPHQ-DLIVSASEDKSIRIWDLNRRTAIQTFRR-DN----------DRFWFVTVHPKLNL

FAAAHDSGVMVFKLE

>UniRef90_A0A1Y2F6H9_12_337 | Coatomer subunit alpha n=1 Tax=Protomyces lactucaedebilis TaxID=56484 RepID=A0A1Y2F6H9_PROLT | E_val=2.2e-176

-QMLTKFESKSSRVKGIAFH--PKRPWLLTSLHNGCIQLWDYRMGSLLD-------RFEE

HDG-AVRAVDFHKT-QPLFVSGGDDYKIRVWNYK--TR-------------KCL-FTLSG

HLDYVRTVFFHHE---YPWILSCSDDQTIRIWNWQ--SRNCIALISGHSHYVMCAQFHPK

-EDLIVSASLDQTVRVWDI--------S---------GL-RKKN-A-A-P------Q---

-----SM---------SL-E--------E-QLQR--------------SQ-A--------

GQGADLF----------------GNTD--CVVKYVLEGHDRGVNWASFHPTKPLIISAGD

DRNVKLWRM------------S----------------DTKAWEVDTCRGHFNNASCAIF

HPTL-DLMLSVGEDKMIRIWDLNKRTAVQAFRR-EN----------DRFWVITAHPTFNL

FAVGHDAGVMVFRLE

>UniRef90_A0A1X2HXL8_1_325 | Coatomer subunit alpha n=1 Tax=Syncephalastrum racemosum TaxID=13706 RepID=A0A1X2HXL8_SYNRA | E_val=1.5e-174

MQMLTKFESKSNRVKGIAFH--AKRPWILASLHNGCIQLWDYRMGTLLE-------RFDE

HDG-PVRGIAFHPT-QPLFVSGGDDYKIKVWNYK--TR-------------RCL-FTLNG

HLDYVRTVFFHHE---YPWILSASDDQTIRIWNWQ--SRNCIAILTGHNHYVMCAQFHPK

-NDLVVSASMDQTVRVWDI--------S---------GL-RKKN-Q-A-P------T---

-----PM---------SF-E--------D-SFA---------------RQ-S--------

-SQADLF----------------GSTD--VMVKYVLEGHDHGVNWASFHPTLPLIVSAGD

DRQVKLWRM------------S----------------ETKAWEVDSCRGHYNNVSSAIF

HPRQ-ELILSDGEDKSVRVWDMTKRTALATFRR-DH----------DRFWVLTAHPELNL

FAAGHDSGLIVFKLE

>UniRef90_F0XN66_7_330 | Coatomer subunit alpha n=6 Tax=Ophiostomataceae TaxID=5152 RepID=F0XN66_GROCL | E_val=1.8e-173

--MLTKFESKSSRAKGLAFH--PKRPWILVSLHSSTIQLWDYRMGTLID-------RFEE

HDG-PVRAVAFHKT-QPLFVSGGDDYKIKVWSYQ--TR-------------RCL-FTLNG

HLDYVRTVFFHDE---LPWIVSCSDDQTIRIWNWQ--NRSLICTMTGHNHYVMCAQFHPK

-DDLIVSASLDQSVRVWDI--------S---------GL-RKKH-S-A-P------T---

-----SM---------SF-E--------D-QMAR--------------SN-Q--------

-NQADMF----------------GNTD--AVVKFVLEGHDRGVNWVAFHPTMPLIVSAGD

DRLVKLWRM------------S----------------ETKAWEVDTCRGHFQNALGCLF

HPHQ-DLILSAGEDKTIRVWDLNKRTPVQTFKR-EN----------DRFWVLAAHPEINL

FAAGHDNGVMVFKLE

>UniRef90_A0A0E9NE57_59_391 | Coatomer subunit alpha n=2 Tax=Saitoella complicata (strain BCRC 22490 / CBS 7301 / JCM 7358 / N | E_val=5.9e-173

MQLLTKFESKSSRAKGIAFHPNPNRPWILTSLHNGTIQLWDYRMGTLLD-------RFEE

HDG-PVRGIAFHPT-KEMFVSGGDDYKIKVWNYK--TR-------------KCL-FTLNG

HLDYVRTVFFHHE---YPWILSASDDQTIRIWNWQ--SRTCIAILTGHNHYVMCAQFHPK

-EDLIVSASLDQTVRVWDI--------S---------NL-RKKS-AVA-P------Q---

-----TM---------TF-E--------E-QMAR--------------AN-S-----AAA

GAGGDLF----------------GNQD--AVCRFVLDGHDRGVNWVSFHPTLPLIVSAGD

DRLVKLWRM------------S----------------ETKAWEVDTCRGHFHNACAALF

HPRQ-ELLLSCGEDKTLRVWNLNTRSSVASFRR-EN----------DRFWVVVAHPEINL

FAAGHDSGVMVFKLE

>UniRef90_A0A197JND9_1_325 | Coatomer subunit alpha n=2 Tax=Linnemannia elongata AG-77 TaxID=1314771 RepID=A0A197JND9_9FUNG | E_val=6.7e-172

MQMLTKFESKSNRVKGIAFH--PKRPWILAALHNGSIQLWDYRMGTLLE-------RFDE

HDG-PVRGIAFHPS-QPLFVSAGDDYKIKVWNYK--TR-------------RCQ-FTLTG

HLDYVRTVYFHHE---CPWIISCSDDQTIRIWNWQ--SRSCIAILTGHNHYVYSAQFHPK

-EDLVVSASLDQTVRVWDI--------S---------GL-RKKN-A-A-P------G---

-----TV---------AH-D--------E-SAH---------------RA-S--------

-QQADIF----------------GSTD--AIVKFVLEGHDRSVNWASFHPTLPLIVSAGD

DRQVKLWRM------------N----------------DSRAWEVDSCRGHTNNVSCTIF

HHRQ-ELILSVGEDKTIRVWDMSKRTAVQTFRR-EH----------DRFWALTAHPELNL

FAAGHDNGLIVFKLE

>UniRef90_A0A063C2H5_6_330 | Coatomer subunit alpha n=2 Tax=Ustilaginoidea virens TaxID=1159556 RepID=A0A063C2H5_USTVR | E_val=1.9e-171

-LTLSQFESKSSRAKGIAFH--PKRPWILVALHSSTIQLWDYRMGTLID-------RFEE

HDG-PVRGVDFHKT-QPLFVSGGDDYKIKVWSYQ--TR-------------RCL-FTLNG

HLDYVRTVFFHHE---LPWILSSSDDQTIRIWNWQ--NRSLICTMTGHNHYTMCAQFHPK

-DDLVVSASLDQSVRVWDI--------S---------GL-RKKH-S-A-P------T---

-----SM---------SY-E--------D-QMVR--------------AN-Q--------

-NQADMF----------------GNTD--AVVKFVLEGHDRGVNWVAFHPTMPLIVSAGD

DRLVKLWRM------------S----------------ETKAWEVDTCRGHFHNASGCLF

HPHQ-DLIISAGEDKTIRVWDLNKRTAVQSFKR-EN----------DRIWVIAAHPEINL

FAAGHDNGVMVFKLE

>UniRef90_A0A6P8BGE8_6_331 | Coatomer subunit alpha n=6 Tax=Pyricularia TaxID=48558 RepID=A0A6P8BGE8_MAGGR | E_val=2.7e-171

-GMLTKFESKSSRAKGIAFH--PKRPWILVSLHSSTIQLWDYRMGTLID-------RFEE

HDG-PVRAIDFHKT-QPLFVSGGDDYKIKVWSYQ--TR-------------RCL-FTLNG

HLDYVRTVFFHHE---LPWIVSASDDQTIRIWNWQ--NRSLICTMTGHNHYAMCAQFHPK

-EDLVVSASLDQSVRVWDI--------S---------GL-RKKH-S-A-P------TS--

-----SL---------SF-E--------D-QMAR--------------NN-A--------

-NQTDMF----------------GNTD--AVVKFVLEGHDRGVNWVAFHPTMPLIVSAGD

DRLVKLWRM------------S----------------ETKAWEVDTCRGHFQNASGCLF

HPHQ-DLILSVGEDKTIRVWDLNKRTGVQSFKR-EN----------DRFWVIAAHPEINL

FAAGHDNGVMVFKLE

>UniRef90_A0A4Q4YG96_6_330 | Coatomer subunit alpha n=12 Tax=Xylariales TaxID=37989 RepID=A0A4Q4YG96_9PEZI | E_val=5.8e-171

-GMLTKFESKSSRAKGIAFH--PKRPWILVSLHSSTIQLWDYRMGTLID-------RFEE

HDG-PVRGIDFHKT-QPLFVSGGDDYKIKVWSYQ--TR-------------RCL-FTLNG

HLDYVRTVFFHHE---LPWIISSSDDQTIRIWNWQ--NRSLICTMTGHNHYVMCAQFHPK

-DDLVVSASLDQSVRVWDI--------S---------GL-RKKH-S-A-P------T---

-----SM---------SF-E--------D-QVAR--------------SN-A--------

-QQADMF----------------GNTD--AVVKFVLEGHDRGVNWVAFHPTAPHIVSAGD

DRLIKLWRF------------S----------------ETKAWEVDTCRGHFQNASGCLF

HPHQ-DLILSCGEDKTIRVWDLQKRTAVQSFKR-EN----------DRFWVIAAHPEINL

FAAGHDNGVMVFKLE

>UniRef90_A0A517LRA5_8_330 | Coatomer subunit alpha n=2 Tax=Venturia TaxID=5024 RepID=A0A517LRA5_9PEZI | E_val=4.3e-170

--MLTKFESKSSRAKGIAFH--SSRPWILVSLHSSTIQLWDYRMGTLID-------RFEE

HDG-PVRGVDFHKT-QPMFVSGGDDYKIKIWSLQ--TR-------------RSI-ATLVG

HLDYVRTVFFHHE---QPWILSCSDDQTIRIWNWQ--NRSLICTMTGHNHYVMCAQWHPS

-QDLVVSASLDQSVRVWDI--------S---------GL-RKKH-S-A-P------T---

-----TM---------SF-E--------D-QMAR--------------AN----------

-NQADVF----------------GNTD--AVVKFVLEGHDRGVNWVAFHPTLPLIVSGGD

DRLVKLWRM------------S----------------ETKAWEVDTCRGHFQNVSACLF

HPHQ-DLIISVGEDKTIRCFDLNKRTSVQTFKR-EN----------DRFWVIAAHPEINL

FASGHDTGVMVFKLE

>UniRef90_A0A1Y2BBK2_1_327 | Coatomer subunit alpha n=1 Tax=Naematelia encephala TaxID=71784 RepID=A0A1Y2BBK2_9TREE | E_val=8e-170

MQMLTKFESKSPRVKGIAFH--PKTPLLAASLHNGTIQLWNYQMGTLVD-------RFDE

HDG-PVRGICFHPT-QPIFCSGGDDYKIKVWNYK--QR-------------KCL-FTLTG

HLDYVRTVFFHRE---YPWIISASDDQTIRIWNWQ--SRTCIAILTGHNHYIMCAQFHPW

-DDLVVSASMDLTVRVWDI--------S---------GL-RKKN-Q-A-H------QA--

-----PM---------SF-E--------E-QMAR--------------AN-Q--------

-GQADLF----------------GNTD--AVVKYVLEGHDRGVNWASFHPTLPLIVSCGD

DRQIKLWRM------------S----------------ETKAWEVDSCRGHFNNVSMTLF

HPRH-ELILSASEDKTIRVWDMTKRTAVQTFRR-EH----------DRFWVLTAHPELNL

FAAGHDNGLIVFKLE

>UniRef90_A0A1Y2E6P5_6_331 | Coatomer subunit alpha n=1 Tax=Pseudomassariella vexata TaxID=1141098 RepID=A0A1Y2E6P5_9PEZI | E_val=7.8e-169

-GMLTKFESKSSRAKGIAFH--PVRPWILVSLHSSTIQLWDYRMGTLID-------RFED

HDG-PVRGIDFHRT-QPLFVSGGDDYKIKVWSYQ--SR-------------RCL-FTLNG

HLDYVRTVFFHHE---LPWIISASDDQTIRIWNWQ--NRSQICTMTGHNHYVMCAQFHPK

-DDLVVSASLDQSVRVWDI--------S---------GL-RKKH-S-A-P------T---

-----SMS--------TY-D--------D-RMSQ--------------SN-A--------

-QQTDMF----------------GNTD--AMVKFVLEGHDRGVNWVAFHPTAPHIVSAGD

DRLVKLWRF------------S----------------ETKAWEVDTCRGHFQNSSGCLF

HPHQ-DLILSCGEDKTIRVWDSLKRTAVQSFKR-EN----------DRFWVIAAHPTINL

FAAGHDNGVMVFKLE

>UniRef90_A0A507CP85_11_332 | Coatomer subunit alpha n=1 Tax=Synchytrium endobioticum TaxID=286115 RepID=A0A507CP85_9FUNG | E_val=1.3e-168

--LLSKFETKSNRVKGLCFH--PKRPWILASLHNGTIQLWDYKMGSLVD-------RFDE

HEG-PVRGIAFHPT-QPLFVSGGDDYKIKLWNWK--LR-------------RCL-FTLNG

HLDYIRTVFFHHE---HPWIISASDDQTIRIWNWQ--SRNCISILTGHNHYVMCAQFHPK

-EDLVVSASLDQTVRVWDI--------S---------GL-RKKH-A-A-P------H---

-----QQ---------SL-D----------ELH---------------RT-P--------

-HQADLF----------------GGTD--AIVKYVLEGHDRGVNWASFHPNLPLIVSGAD

DRQVKLWRM------------N----------------ETKAWEVDTCRGHFNNVSCVIF

HPRQ-ELIVSDSEDRTIRIWDMTKRVVLQTFRR-EN----------DRFWIMAAHPELNL

FAAGHDSGLMVFKLE

>UniRef90_A0A6P8HXX5_1_322 | Coatomer subunit alpha n=3 Tax=Actiniaria TaxID=6103 RepID=A0A6P8HXX5_ACTTE | E_val=2.5e-168

--MLTKFETKSARVKGLCFH--PKRPWVLASLHNGVIQLWDYRMCTLLE-------RFDE

HDG-PVRGIHFHDV-QPLFVSGGDDYKIKVWNYK--QK-------------RCL-FTLLG

HLDYIRTTFFHHE---YPWILSCSDDQTIRIWNWQ--SRTCICVLTGHNHYVMCAQFHPS

-EDLVVSASLDQTVRVWDM--------S---------GL-RKKT-V-A-P------G---

-----AS---------GL-D--------D-HL----------------KS-P--------

-GHTDLF----------------GQSD--AIVKHVLEGHDRGVNWVAFHPTMPLIVSGAD

DRQVKLWRM------------N----------------DSKAWEVDTCRGHYNNVSCVLF

HPRQ-ELMLSNSEDKSIRVWDLSKRTGVQTFRR-EH----------DRFWAIAAHPTLNL

FAAGHDSGMIVFKLE

>UniRef90_A0A139AT68_1_324 | Coatomer subunit alpha n=1 Tax=Gonapodya prolifera (strain JEL478) TaxID=1344416 RepID=A0A139AT6 | E_val=3.5e-168

--MLSKFETKSNRVKGLSFH--PKRPWILASLHNGSVQLWDYKMGTLID-------RFDE

HEG-PVRGICFHPT-QPLFVSGGDDYKVKVWNWK--LR-------------RCL-FTLNG

HLDYIRTVFFHHE---LPWIISASDDQTIRIWNWQ--SRNCISILTGHNHYVMCAQFHPK

-DDLVVSASLDQTVRVWDI--------S---------GL-RKKH-A-AGP------Q---

-----TS---------QT-E--------D-SAI---------------RS-G--------

-QQADIF----------------GGTD--AVVKYVLEGHDRGVNWASFHPTLPLIVSGAD

DRQVKLWRM------------N----------------DTKAWEVDTCRGHYNNVSCVLF

HPRQ-ELILSDCEDKTIRVWDMTKRTCLQTFRR-EH----------DRFWILAAHPELNL

FAAGHDSGLIVFKLE

>UniRef90_UPI0006416C7A_1_323 | coatomer subunit alpha-like n=1 Tax=Hydra vulgaris TaxID=6087 RepID=UPI0006416C7A | E_val=8.8e-168

--MLTKFETKSARVKGLAFH--SKRPWVLASLHNGVIQLWDYRMCTLLD-------RFDE

HDG-PVRGIDFHEN-QPLFVSGGDDYKIKVWNYK--QK-------------KCI-FTLLG

HLDYIRTTFFHHE---YPWIVSCSDDQTIRIWNWQ--SRNCINVLTGHNHYVMCAQFHKT

-EDYIVSASLDQTVRVWDI--------S---------GL-RKKF-A-S-P------G---

-----TK---------DR-D--------DTSV----------------KN-P--------

-GQIDLF----------------GHAD--AVVKHVLEGHDRGVNWVTFHPTMPLIVSAAD

DRQVKLWRM------------N----------------ESKAWEVDTCRGHYNNVSSVIF

HPRQ-ELILSNSEDKSIRVWDMSKRTGVQTFRR-EN----------DRYWILAAHPTLNL

FAAGHDSGMVVFKLE

>UniRef90_A0A6M2DKH2_1_322 | Coatomer subunit alpha n=2 Tax=Pulicidae TaxID=7511 RepID=A0A6M2DKH2_9NEOP | E_val=2.5e-167

--MLTKFETKSARVKGISFH--PKRPWILTSLHNGIIQLWDYRMCTLLD-------KFDE

HDG-PVRGICFHNQ-QPLFVSGGDDYKIKVWNYK--QR-------------RCI-FTLLG

HLDYIRTTLFHHE---YPWILSASDDQTIRIWNWQ--SRTCICVLTGHNHYVMCAMFHPS

-EDTLVSASLDQTVRVWDI--------S---------GL-RKKN-V-A-P------G---

-----PG---------GL-E--------D-HL----------------KN-P--------

-GATDLF----------------GQAD--AVVKYVLEGHDRGVNWAHFHPTLPLIVSGAD

DRLIKLWRM------------N----------------EYKAWEVDICRGHYNNVSCVLF

HPRQ-ELIISNSEDKSIRVWDMTKRTCLHTFRR-EH----------ERFWVLTAHPTLNL

FAAGHDTGMIVFKLE

>UniRef90_UPI0006B0E6B7_1_322 | coatomer subunit alpha-like n=1 Tax=Limulus polyphemus TaxID=6850 RepID=UPI0006B0E6B7 | E_val=4.7e-167

--MLTKFETKSARVKGLSFH--PKRPWVLTSLHNGVIQLWDYRMCTLLD-------KFDE

HDG-PVRGICFHNQ-QPLFVSGGDDYKIKVWNYK--HR-------------RCI-FTLLG

HLDYIRTTTFHHE---YPWILSASDDQTIRIWNWQ--SRTCIYVLTGHNHYVMCAQFHPS

-EDLVVSASLDQTVRVWDI--------S---------GL-RKKN-V-A-P------G---

-----PT---------GL-D--------E-HL----------------KN-P--------

-SHTDLF----------------GQSG--AVVKHVLEGHDRGVNWAAFHPTMPLIVSGAD

DRQIKLWRM------------N----------------DSKAWEVDTCRGHYNNVSCVTF

HPRQ-ELILSNSEDKSIRVWDMSKRTCLHTFRR-DH----------DRFWILTAHPSLNI

FAAGHDSGMIIFKLE

>UniRef90_A0A6J8CPW3_1_322 | Coatomer subunit alpha n=3 Tax=Mytilus coruscus TaxID=42192 RepID=A0A6J8CPW3_MYTCO | E_val=1e-166

--MLTKFETKSARVKGLTFH--PKRPWVLTSLFNGVIQLWDYRMCTLID-------KFDE

HDG-PVRGICFHSQ-QPLFVSGGDDYKIKVWNYK--QK-------------RCL-FTLLG

HQDYIRTTFFHHE---YPWILSASDDQTIRVWNWQ--SRTCVCVLAGHSHFVMCAQFHPS

-EDLIVSASLDQTVRVWDI--------S---------GL-RKKN-V-S-P------G---

-----PS---------SI-E--------D-RL----------------RA-T--------

-GQTDLF----------------GMSD--AVVKHVLEGHDRGVNWAAFHPTLPLIVSGAD

DRQVKLWRM------------N----------------DAKAWEVDTCRGHYNNVSCCTF

HPRH-ELILSNSEDKSIRVWDMSKRTGVQTFRR-EH----------DRFWIMAAHPTLNL

FAAGHDSGMIVFKLE

>UniRef90_A0A316VFL0_1_326 | Coatomer subunit alpha n=1 Tax=Meira miltonrushii TaxID=1280837 RepID=A0A316VFL0_9BASI | E_val=1.9e-166

MQMLTKFESKSNRVKGIAFH--PRLPLLASALHNGSIQLWNYQMGTIYD-------RLDE

HEG-PVRGVDFHPT-QPLLVSGGDDYRIKVWNHK--TR-------------RCL-FTLNG

HLDYVRTTYFHHT---QPWILSASDDQTIRIWNWQ--SRSCIAILTGHNHYIMCAQFHPR

-EDLIVSASMDQTVRVWDI--------S---------GL-RKKN-T-S-A------Q---

-----PM---------SF-E--------E-QIAR--------------AN-T--------

-GQADLF----------------GNTD--AMVKYVLEGHERGVNWATFHPTLPLIVSGGD

DRQVKLWRM------------S----------------ETKAWEVDTCRGHFNNISSCLF

HPRH-ELIVSDGEDKTIRVWDMGKRTAVQTFRR-EH----------DRFWVLTAHPNLNL

FAAGHDNGLIVFKLE

>UniRef90_UPI001402FFD1_1_321 | coatomer subunit alpha n=1 Tax=Petromyzon marinus TaxID=7757 RepID=UPI001402FFD1 | E_val=5.9e-166

--MLTKFETKSARVKGLCFH--SKRPWVLASLHSGVIQLWDYRMCTLID-------KYDE

HDG-PVRGVDFHKQ-QPLFVSGGDDYKIKVWNYK--LR-------------RCL-FTLLG

HLDYIRTTFFHHE---YPWIISCSDDQTIRIWNWQ--SRTCVCVLTGHNHYVMCAQFHPM

-EDLVVSASLDQTVRVWDI--------S---------GL-RKKN-L-N-P------G---

-----SS---------G--D--------A-EL----------------RS-I--------

-SGVDLF----------------GAAD--AVVKHVLEGHDRGVNWAAFHPTMPLIVSGAD

DRQVKLWRM------------N----------------DSKAWEVDTCRGHYNNVSCCLF

HPRH-ELILSNSEDKSIRVWDMSKRTGVQTFRR-DH----------DRFWVLAAHPSLNL

FAAGHDSGMIVFKLE

>UniRef90_A0A7M5XL95_1_323 | Coatomer subunit alpha n=1 Tax=Clytia hemisphaerica TaxID=252671 RepID=A0A7M5XL95_9CNID | E_val=1.2e-165

--MLTKFETKSARVKGICFH--SKRPWVLASLHNGVIQLWDYRMCTLLE-------RFDE

HDG-PVRGIHFHEH-QPLFVSGGDDYKIKVWNYK--QK-------------KCI-FTLLG

HLDYIRTTFFHHE---YPWILSCSDDQTIRIWNWQ--SRNCISVLTGHNHYVMCAQFHPT

-EDLIVSASLDQTVRVWDF--------G---------GL-RRKF-V-S-P------G---

-----AK---------AK-E--------ETGP----------------KT-P--------

-GQMDLF----------------GQAD--AIVKHVLEGHDRGVNWVAFHPSMPFIVSAAD

DRQVKLWRM------------N----------------DAKAWEVDTCRGHYNNVSSALF

HPRQ-ELIISNSEDKSIRVWDMSKRTGVQTFRR-EH----------DRFWTLAAHPTLNL

FAAGHDSGMLVFKLE

>UniRef90_A0A0L0HKE1_1_325 | Coatomer subunit alpha n=2 Tax=Spizellomyces TaxID=4815 RepID=A0A0L0HKE1_SPIPD | E_val=1.7e-165

--MLSKFETKSNRVKGLAFH--PKRPWILAALHNGAIQLWDYKMGTLID-------RFDE

HEG-PVRGIAFHHT-QPLFVSGGDDYKIKVWNWK--TR-------------RCL-FTLNG

HIDYIRTVFFHHE---SPWIISASDDQTIRIWNWQ--SRNCISILTGHNHYVMCAQFHPK

-DDLVVSASLDQTIRVWDI--------S---------GL-RKKH-A-AAP------S---

-----SI---------TT-D----------YEHR-------------LGA-A--------

-GQADIF----------------GNTD--AMVKYVLEGHDRGVNWATFHPNLPLIVSSSD

DRTVKLWRM------------N----------------ETKAWEVDTCRGHYNNVSSVLF

HPRQ-DLIISDSEDKTIRIWDMTKRTALQTFRR-EH----------DRFWILAAHPELNL

FAAGHDSGLIVFKLE

>UniRef90_UPI0009950A54_1_322 | coatomer subunit alpha n=1 Tax=Pseudomyrmex gracilis TaxID=219809 RepID=UPI0009950A54 | E_val=3.6e-165

--MLTKFETKSARVKGLSFH--PKRPWVLASLHNGVIQLWDYRMCTLLD-------KFDE

HDG-PVRGICFHNQ-QPLFVSGGDDYKIKVWNYK--QR-------------RCI-FTLLG

HLDYIRATVFHHE---YPWILSASDDQTIRIWNWQ--SRTCICVVTGHNHYVMCAQFHPT

-EDLIISASLDQTIRIWDI--------S---------GL-RKKN-V-A-P------G---

-----PG---------GL-E--------D-HL----------------KN-P--------

-GATDLF----------------GQAD--AALKYMLEGHDRGVNWACFHPTLPLILSGAD

DRQIKLWRM------------N----------------DAKAWEVDTCRGHYNNVSCVLF

HPRQ-DLILSNSEDKSIRVWDMTKRTCLHTFRR-EH----------ERFWVLAAHPTLNL

FAAGHDSGMIIFKLE

>UniRef90_A0A7I8V772_1_320 | Coatomer subunit alpha n=1 Tax=Dimorphilus gyrociliatus TaxID=2664684 RepID=A0A7I8V772_9ANNE | E_val=5.1e-165

--MLTKFESKSARVKGLCFH--PKRPWILASLHNGVIQLWDYRMATLID-------KYDE

HDG-PVRGLCFHNQ-QPLFVSGGDDYKIKIWNYK--LK-------------RCL-FTLTG

HLDYIRTTTFHHE---YPWILSCSDDQTIRIWNWQ--NRSCVTVITGHSHYVMCAAFHPT

-EDLIVSCSLDQTIRVWDT--------S---------AI-RKKN-V-S-P------G---

-----IM------------S--------D-YG----------------RR-Q--------

-DQANLF----------------GVTD--AVVKHVLEGHDRGVNWVAFHPTLPLIVSGAD

DRSIKLWRM------------N----------------DSRAWEVDTCRGHFNNVSCVLF

HPRM-DLILSNAEDKTIRVWDMSKRTAIQSFRR-EN----------DRFWVLAAHPTLNL

FAAGHDAGMVLFKLE

>UniRef90_UPI000719B932_1_323 | coatomer subunit alpha-like n=1 Tax=Priapulus caudatus TaxID=37621 RepID=UPI000719B932 | E_val=6.5e-165

--MLTKFETKSARVKGLSFH--PKRPWILSTLHTGVIQLWDYRMCTLLD-------KYDE

HDG-PVRGISFHSQ-QPLFVSGGDDYKIKVWNYK--QK-------------RCI-FTLLG

HLDYIRTTFFHHE---YPWVLSASDDQTIRIWNWQ--SRTCICVLTGHNHYVMCANFHPS

-DDLVVSASLDQTVRVWDI--------S---------GL-RKKN-V-A-P------G---

-----PG---------GL-D--------D-HV----------------KNQA--------

-GHTDLF----------------GTAD--AIVKHVLEGHDRGVNWAAFHATMPLIVSGAD

DRQVKLWRM------------N----------------DAKAWEVDTCRGHYNNVSCALF

HPRQ-DLIISNSEDKSIRVWDMAKRTCLQTFRR-EG----------DRFWVITSHPALNL

FAAGHDSGMIVFKLE

>UniRef90_A0A0K3CS52_1_326 | Coatomer subunit alpha n=5 Tax=Rhodosporidium toruloides TaxID=5286 RepID=A0A0K3CS52_RHOTO | E_val=1.4e-164

MQMLTKFESRSNRVKGISFH--PRLTLLAASLHSGSIQLWNFQMGVLVD-------RFEE

HDG-PVRGIDFHPT-QPLFVSGGDDYKIKVWNYK--TR-------------RCL-FTLHG

HLDYVRTVFFHPE---QPWIISASDDQTIRIWNWQ--SRTCIAILTGHNHYIMCAQFHPK

-EDYVVSASMDQTVRVWDI--------S---------GL-RKKS-T-T-A------A---

-----PL---------SF-E--------E-QIQR--------------AN-A--------

-GQADLF----------------GNTD--AVVKYVLEGHDRGVNWASFHPTLPLIVSCGD

DRQIKLWRM------------S----------------ETKAWEVDTCRGHYNNVSQVLF

HPKH-ELIISDSEDKTIRVWDMTKRTAVQTFRR-EN----------DRFWVLAAHPKLNL

FAAGHDSGLIVFKLD

>UniRef90_J9K3N8_1_322 | Coatomer subunit alpha n=12 Tax=Aphididae TaxID=27482 RepID=J9K3N8_ACYPI | E_val=4.3e-164

--MLTKFETKSARIKGLTFH--SKRPWILASLHTGVIQLWDYRMCTLLD-------KFDE

HDG-PVRGISFHSQ-QPIFVSGGDDYKIKVWNYT--QR-------------RCI-FTLLG

HLDYIRSTMFHHE---YPWILSASDDQTIRIWNWQ--SRACICVLTGHNHYVMCAQFHPS

-EDLVVSASLDQTVRVWDI--------S---------GL-RKKN-V-A-P------G---

-----PG---------GL-D--------D-HL----------------KN-P--------

-NATDLF----------------GQAD--AVVKHVLEGHDRGVNWCSFHPTLPLIVSGAD

DRQIKLWRM------------N----------------DSKAWEVDTCRGHYNNVSCVVF

HPKQ-ELILSNSEDKSIRVWDMTKRTCLNTFRR-EH----------ERFWVLAAHPTSNL

FAAGHDSGMIIFKLE

>UniRef90_A0A6P7GHR3_1_322 | Coatomer subunit alpha n=2 Tax=Diabrotica virgifera virgifera TaxID=50390 RepID=A0A6P7GHR3_DIAVI | E_val=6.8e-164

--MLTKFETKSARVKGLSFH--PKRPWILTSLHNGCIQLWDYRMCTLLE-------KFDE

HDG-PVRGICFHNQ-QPLFVSGGDDYKIKVWNYK--QK-------------RCI-FTLLG

HLDYIRTTLFHHE---YPWIVSASDDQTIRIWNWQ--SRTCICVLTGHNHYVMCVSFHPS

-EDLLVSASLDSTVRVWDI--------S---------GL-RKKN-V-A-P------G---

-----PA---------GL-E--------E-HL----------------KN-P--------

-GATDLF----------------GQAD--AVVKHVLEGHDRGVNWASFHPTLPLIASGAD

DRQVKLWRM------------N----------------DSKAWEVDTCRGHWHNVSCVLF

HPRQ-ELILSNSEDKTIRVWDTTKRTCLHTFKR-EN----------ERFWIMASHPNLNL

FAAGHDGGMVIFKLE

>UniRef90_A0A131ZSG2_63_384 | Coatomer subunit alpha-like protein n=1 Tax=Sarcoptes scabiei TaxID=52283 RepID=A0A131ZSG2_SARSC | E_val=1.1e-163

--MLTKFESKSSRVKGIAFH--PKRQWVLASLHSGLIQLWDYRVGTLID-------KFDE

HDG-PVRGIDFHNQ-QPLFVSGGDDYKIKMWNYK--SR-------------RCI-FTLLG

HLDYIRVVKFHHE---YPWILSCSDDQTIRIWNWQ--SRTNISILTGHNHYVMCAQFHPT

-EDLVVSCSLDQTVRVWDI--------S---------GL-RKKN-V-S-P------G---

-----PG---------GL-E--------E-HL----------------KN-P--------

-HSTELF----------------GPSD--VYVKHVLEGHDRGVNWASFHPTMPLIVSGAD

DRLIKLWRM------------N----------------ESKAWEVDSCRGHYNNVSCVMF

HPKQ-DLIISDSEDKSIRVWDLTKRTCLYTFRR-EH----------ERFWILAAHPNQNL

FAAGHDGGIIVFKLE

>UniRef90_A0A7R9ADV1_1_322 | Hypothetical protein n=1 Tax=Darwinula stevensoni TaxID=69355 RepID=A0A7R9ADV1_9CRUS | E_val=1.3e-163

--MLTKFETKSARVKGLSFH--QKRPWILASCHNGNIQLWDYRMCVLLD-------KFEE

HDG-PVRGICFHQQ-QPLFVSGGDDYKIKVWNYR--KR-------------RCL-FTLLG

HLDYIRTTFFHHE---YPWILSASDDQTLRIWNWQ--SRSCICVMTGHNHYVMCAQFHPS

-DDLVVSASLDQTVRVWDI--------S---------GL-RKKN-V-A-P------G---

-----PG---------GL-E--------E-HL----------------RN-P--------

-ASTDLF----------------GQAD--AIVRHLLEGHDRGVNWVAFHPTLPLIVSAAD

DRQIKLWRM------------N----------------DTRAWEVDTCRGHYNNVSCVLF

HPRQ-ELLLSNSEDKSIRVWDMTKRTCLHTFRR-EH----------DRFWVLAAHPSLNL

FAAGHDGGMLVFKLE

>UniRef90_A0A0A9ZAE9_1_322 | Coatomer subunit alpha n=3 Tax=Miridae TaxID=30083 RepID=A0A0A9ZAE9_LYGHE | E_val=1.6e-163

--MLTKFETKSARVKGLSFH--PKRPWILASLHSGVIQLWDYRMCTLLD-------KFDE

HDG-PVRGICFHTQ-QPLFVSGGDDYKIKVWNYK--HR-------------RCI-FTLLG

HLDYIRTTVFHQE---YPWILSASDDQTIRIWNWQ--SRTCICVLTGHNHYVMCAQFHPT

-DDIVVSASLDMTVRVWDI--------S---------GL-RKKN-V-A-P------G---

-----PG---------GL-E--------E-HL----------------KN-P--------

-TATDLF----------------GQAD--AVVKHVLEGHDRGVNWAAFHPTLPLIVSGAD

DRHIKLWRM------------N----------------ESKAWEVDTCRGHYYNVSCVLF

HPRQ-DLILSNSEDRSIRVWDMTKRTCLHTFRR-EN----------DRFWVLTAHPTLNL

FAAGHDLGMIIFKLE

>UniRef90_A0A3M7S875_1_322 | Coatomer subunit alpha n=1 Tax=Brachionus plicatilis TaxID=10195 RepID=A0A3M7S875_BRAPC | E_val=4.6e-163

--MLIKFETKSARVKGLSFH--PKRPWVLASLHNGIIQLWDYRMSTMID-------KFDE

HDG-PVRGIAFHSQ-QPLFVSGGDDYKIKVWNYK--QR-------------RCL-FTLLG

HLDYIRTTFFHHE---YPWIVSASDDQTIRIWNWQ--SRNCVSVLTGHNHYVMCAQFHPS

-EDFIVSASLDQSVRVWDI--------S---------GL-RKKN-V-A-P------G---

-----PE---------GF-H--------S-HLQ---------------RS-T--------

-GTPDLF----------------GQTD--AVVKHVLEGHDRGVNWASFHPTLPLIVSAAD

DRQVKLWRM------------N----------------DTKAWEVDTCRGHYNNVSCAIF

HPKQ-ELMLSNSEDKSIRVWDMSKRTCLNTFRR-D-----------DRYWIITGHPTLSL

FAAGHDNGMIIFKLE

>UniRef90_A0A482WL14_1_321 | Coatomer subunit alpha n=3 Tax=Delphacidae TaxID=33362 RepID=A0A482WL14_LAOST | E_val=8.1e-163

--MLTKFETKSARVKGLSFH--PKRPWVLSSLHSGIIQLWDYRMCTLLD-------KFDE

HDG-PVRGICFHNQ-QPIFVSGGDDYKIKVWNYK--QR-------------RCI-FTLLG

HLDYIRTTVFHHE---YPWILSASDDQTIRIWNWQ--SRNCISVLTGHNHYVMCAQFHPS

-EDIIVSSSLDQTVRVWDI--------S---------GL-RKK--V-A-P------G---

-----PG---------GF-D--------D-HL----------------KN-S--------

-TTTDLF----------------VPPD--AVVKHVLEGHDRGVNWACFHPSLPFIASGAD

DRQIKLWRM------------N----------------EAKAWEVDTCRGHYNNVSCVLF

HPRQ-ELILSNSEDKSIRVWDMNKRVCLHTFRR-EH----------DRFWALTAHPTLNL

FAAGHDAGMVIFKLE

>UniRef90_A0A433PVX9_1_336 | WD40-repeat-containing domain protein n=1 Tax=Endogone sp. FLAS-F59071 TaxID=2340872 RepID=A0A43 | E_val=9.3e-163

MQMLTKFESKSNRVKGIAFH--AKRPWILASLHNGCIQLWDYRMGTLLE-------RFDE

HDG-PVRGISFHAT-QPLFVSGGDDYKIKVWNHK--TR-------------RCL-FTLNG

HLDYVRSVFFHHE---YPWIISASDDQTIRIWNWQ--SRNCIAILTGHNHYVMCAQFHPK

-DDLVVSASLDTTVRVWDISGMYSDTGS---------GL-RKKN-Q-A-P------G---

-----SN---------SY-E--------D-RLQ---------------AQ-S--------

-GQPDLF----------------GSTD--AIVKYVLEGHERGVNWATFHPSLPLIVSCGD

DRQAYLLSLND---------CS----------------DTKAWEVDTCRGHFNNVSCSIF

HPRQ-DLIISDAEDKSIRVWDMSKRTAVATFRR-DH----------DRYWVLTAHPELNL

FAAGHDNGLIVFKLE

>UniRef90_A0A066VNJ2_1_343 | Putative COP1-coatomer complex alpha chain of secretory pathway vesicles n=1 Tax=Tilletiaria ano | E_val=1.1e-162

MQMLTKFESKSNRVKGIAFH--PSLPLLAATLHNGSIQLWNYQTGTIYD-------RLDE

HDG-PVRGICFHPT-QPLLVSGGDDYKIKVWNHK--TR-------------TCL-FTLNG

HLDYVRTVFFHHT---QPWILSASDDQTIRIWNWQ--SRQCIAILTGHNHYIMCAQFHPT

-EDLIVSASMDQTVRVWDI--------S---------GL-RKKS-A-A-S------SSNG

TASSMPM---------SI-E--------E-QIAR--------------AA-SGSGLPGAL

GGQADLF----------------GSTD--AMVKYVLEGHDRGVNWASFHPTLPLIVSAGD

DRQVKLWRM------------S----------------ETKAWEVDTCRGHFNNVSSALF

HPRH-ELIISDAEDKTIRVWDMGKRTAVQTFRR-EA----------DRFWVLTAHPKLNL

FAAGHDSGLIVFKLE

>UniRef90_A0A7R9TAQ9_1_323 | Hypothetical protein n=1 Tax=Prasinoderma coloniale TaxID=156133 RepID=A0A7R9TAQ9_9VIRI | E_val=1.9e-162

--MLTKFETKSNRVKGLSFH--PKRPWILASLHSGVVQLWDYRMGTLID-------RFDE

HDG-PVRGVHFHKS-QPLFVSGGDDYKVKVWNYK--NR-------------RCL-FTLLG

HLDYIRTVQFHEE---HPWIVSASDDQTIRVWNWQ--SRSCIAVLTGHNHYVMCACFHLK

-EDLVVSASLDQTVRVWDI--------S---------GL-RKRS-A-A-P------G---

-----SQ---------GP-D--------E-LMQ---------------LG-Q--------

-MNSDLF----------------GGGD--AVVKYVLEGHDRGVNWAAFHPSLPLIVSGAD

DRQVKLWRM------------N----------------DTKAWEVDTLRGHLNNVSCVMF

HARQ-DIIVSNSEDKSIRVWDMSKRTGVQTYRR-EH----------DRFWILAAHPELNL

LAAGHDSGMIVFKLE

>UniRef90_A0A2G8KYM4_1_322 | Coatomer subunit alpha n=1 Tax=Stichopus japonicus TaxID=307972 RepID=A0A2G8KYM4_STIJA | E_val=2.7e-162

--MLTKFETKSARVKGLSFH--KNRPWILASLHSGVIQLWDYRMCTLLD-------KFDE

HDG-PVRGICFHDQ-QPLFVSGGDDYKIKLWNYK--LK-------------RCL-FTLLG

HLDYIRTTFFHHE---YPWILSASDDQTIRIWNWQ--SRNCICVLSGHNHYVMCANFHPS

-DDMVVSASLDQTVRVWDI--------S---------GL-RKKN-V-Q-P------G---

-----PS---------GL-D--------D-HL----------------KN-T--------

-MTPELF----------------GTAD--ATVKHQLEGHDRGVNWAAFHPTMPLIVSAAD

DRYVKLWRM------------N----------------DAKAWEVDTCRGHYNNVSCAIF

HPRQ-ELLISNSEDKSVRVWDMSKRMCLHTFRR-DH----------DRFWVMTAHPSLNL

FAAGHDSGMIVFKLE

>UniRef90_A0A1E4RQB8_1_328 | Coatomer subunit alpha n=1 Tax=Hyphopichia burtonii NRRL Y-1933 TaxID=984485 RepID=A0A1E4RQB8_9A | E_val=3.7e-162

MKMLTKFESKSSRAKGVAFH--PKRPWVLVALHSSTIQLWDYRMGTLID-------RFED

HDG-PVRCVDFHPT-QPLFVSGSDDYSIKVWSLN--TR-------------KCI-FTLNG

HLDYLRTVSFHHD---LPWILSCSDDQTIRIWNWQ--NRQEIACLTGHNHYVMSAQFHPS

-DDLIVSASLDQTVRVWDI--------S---------GL-RKKH-S-A-P------T---

-----SSM-------RSF-E--------E-QIQR--------------QQ-L--------

-PQQDIF----------------GNVN--AIVKYVLEGHDKGVNWAAFHPTLPLIVSAAD

DRLVKLWRM------------S----------------DTKAWEVDTCRGHTGNVLSAVF

HPHQ-DLILSVSDDKTIRVWDLNKRTPVKQFRR-EH----------DRFWLVASHPTINL

FATCHDSGVMVFKLE

>UniRef90_A0A2N1JEM6_1_326 | Coatomer subunit alpha n=1 Tax=Malassezia vespertilionis TaxID=2020962 RepID=A0A2N1JEM6_9BASI | E_val=6.4e-162

MQMLTKFESKSNRVKGIAFH--PKLALLASSLHNGSIQLWNYQTGTIYE-------RLED

HEG-PVRGICFHPT-QPLLVSGGDDYKIKVWNHK--TG-------------KVL-FTLHG

HLDYVRSVFFHHE---HPWIISASDDQTIRIWNWQ--SRTCIAILTGHNHYVMCAQFHPY

-EDLIVSASMDQTVRVWDY--------T---------AL-KQKS-T-T-A------Q---

-----PM---------SL-D--------E-QMAR--------------AS-S--------

-AQMDLF----------------GHMD--VMVKYVLEGHERGVNWAAFHPALPLIVSASD

DRTVKLWRM------------S----------------ETKAWEVDTCRGHYNNVSAALF

HPHA-ELILSVSEDKTIRVWDMGKRTAVQTFRR-EH----------DRFWVLTAHPQLNL

FAAGHDSGLIVFKLE

>UniRef90_G8YDX3_1_328 | Coatomer subunit alpha n=2 Tax=Pichia sorbitophila (strain ATCC MYA-4447 / BCRC 22081 / CBS 7064 / N | E_val=9.5e-162

MKMLTKFESKSSRAKGVAFH--PTRPWILVALHSSTIQLWDYRMGTLID-------RFEE

HIG-PVRTVNFHPT-QPLFVSGGDDFTIKVWSLQ--SR-------------KCI-FTLNG

HLDYIRTVSFHRD---LPWIISASDDQTIRIWNWQ--NRQEIACLTGHNHYVMSAEFHPT

-EDLIVSASLDQTVRVWDI--------S---------GL-RKKH-S-A-P------T---

-----SSM-------RSF-E--------D-QLQR--------------QQ-L--------

-PQQDIF----------------GNVN--AIVKYVLEGHDRGVNWATFHPTLPLIVSAGD

DRLVKLWRM------------S----------------ETKAWEVDTCRGHTGNVLCAVF

HPNQ-DLIISIADDKTVRVWDLNKRTPVKQFRR-EH----------DRFWLIACHPHINL

FAACHDSGVMVFKLE

>UniRef90_A5DXZ8_1_328 | Coatomer subunit alpha n=1 Tax=Lodderomyces elongisporus (strain ATCC 11503 / CBS 2605 / JCM 1781 / | E_val=1.4e-161

MKMLTKFESKSSRAKGVAFH--PKRPWCLVSLHSSTIQLWDYRMGTLID-------RFED

HVG-PVRCVNFHPT-QPLFVSGGDDYSIKVWSLN--TR-------------KCI-FTLNG

HLDYVRGVSFHHD---LPWIISCSDDQTIRIWNWQ--NRQEIACLTGHNHYVMSAQFHPT

-EDLIVSASLDQTVRVWDI--------S---------GL-RKKH-S-A-P------T---

-----SSV-------RSF-E--------D-QLQR--------------QQ-L--------

-PQQDIF----------------GNVN--AIVKYVLEGHDKGVNYAAFHPTLPLIVSAGD

DRLVKLWRM------------S----------------DTKAWEVDTCRGHTGNVLSAIF

HPHQ-DMILSVSDDKTIRVWDLNKRVPIKQFRR-EN----------DRFWLIASHPTINL

FAACHDSGVMVFKLE

>UniRef90_A0A131XLR0_1_322 | Coatomer subunit alpha n=17 Tax=Ixodidae TaxID=6939 RepID=A0A131XLR0_9ACAR | E_val=2.2e-161

--MLTKFETKSARVKGLSFH--PKRPWILASLHNGVIQLWDYRMCTLLD-------KFDE

HDG-PVRGICFHNQ-QPLFVSGGDDYQIKVWNYK--QR-------------RCM-FTLLG

HLDYIRTTTFHHE---YPWILSSSDDQTIRVWNWQ--SRTCICVLTGHTHYVMCAQFHPS

-EDLMVSASLDQTIRVWDL--------G---------GL-RKKN-V-A-P------G---

-----PG---------GL-D--------D-HL----------------KN-P--------

-GHTDLF----------------GTSD--AVVRHFLDGHERGVNWAVFHPTAPLVVSGAD

DRQIKLWRM------------N----------------ESKAWEVDTCRGHYCNVSCVLF

HPRQ-ELILSNSEDKSIRVWDMSKRTCLHTFRR-EH----------DRFWILASHPSLNL

FAAGHDSGMILFKLE

>UniRef90_A0A3G2S3X6_1_326 | Coatomer subunit alpha n=1 Tax=Malassezia restricta CBS 7877 TaxID=425264 RepID=A0A3G2S3X6_9BASI | E_val=3.1e-161

MQMLTKFESKSNRVKGIAFH--QKLPLLAASLHNGSIQLWNYQTGTIYE-------RLED

HEG-PVRGVCFHPT-QPLLVSGGDDFKIKVWNHK--TG-------------KVL-FTLHG

HLDYVRSVFFHHE---HPWIISASDDQTIRIWNWQ--SRTCIAVLTGHNHYVMCAQFHPY

-EDLIVSASMDQTVRVWDF--------T---------TL-KQKS-T-T-A------Q---

-----PM---------SL-E--------D-QIAR--------------AN-S--------

-TQMDLF----------------ANMD--VVVKYVLEGHDRGVNWAAFHPALPLIVSASD

DRQIKLWRM------------S----------------DTKAWEVDTCRGHYNNVSASLF

HPHA-ELILSVSEDKTIRVWDMGKRIAVQTFRR-EN----------DRFWVLTAHPHLNL

FAAGHDSGLIVFKLE

>UniRef90_A0A7S2V4C4_1_323 | Hypothetical protein (Fragment) n=1 Tax=Fibrocapsa japonica TaxID=94617 RepID=A0A7S2V4C4_9STRA | E_val=5.5e-161

--MLTKFESKSNRVKGLSFH--PVRPWILASLHNGIIQLWDYRMGTLLD-------RFDE

HDG-PVRGVDFHAR-QPLIVSGGDDYKIKVWDYK--MR-------------RCL-FTLLG

HLDYIRTVRFHSE---YPWIVSASDDQTIRIWNWQ--SRGCVAVLTGHNHYVMCASFHPK

-DDLIVSASLDQSVRVWDI--------S---------GL-RKKT-V-R-G------A---

-----PS---------AD-D--------P-ANM---------------VA-R--------

-VNADLF----------------GGND--AVVKYVLEGHDRGVNWASFHPTLPLVISGAD

DRQVKLWRM------------N----------------ETKAWEVDTMRGHTNNVSCVIF

HPKH-ELIVSNSEDRSIRVWDISKRLGVQTFRR-EN----------DRFWILAAHPEQNL

LAAGHDSGMIVFKLE

>UniRef90_A0A507C8J5_15_343 | Coatomer subunit alpha n=1 Tax=Synchytrium microbalum TaxID=1806994 RepID=A0A507C8J5_9FUNG | E_val=7.2e-161

--LLSKFETKSNRVKGLTFH--PKRPWILASLHNGTLQLWDYKMGTLISVTLHTQLNPCT

NAG-PVRGIAFHPT-QPLFVSGGDDYKIKVWNWK--TR-------------RCL-FTLNG

HLDYIRTVFFHHE---HPWIISASDDQTIRIWNWQ--SRNCISILTGHNHYVMCAQFHPK

-EDLVVSASLDQTVRVWDI--------S---------GL-RKKH-A-A-P------Q---

-----QP---------TL-E--------D--LH---------------RG-P--------

-NQADLF----------------GGTD--AIVKYVLEGHDRGVNWASFHPNLPLIVSGAD

DRQVKLWRM------------N----------------ETKAWEVDTCRGHYNNVSCVIF

HPRQ-ELIISDSEDRTIRIWDMTKRTVLQTFKR-EN----------DRFWIMAAHPELNL

FAAGHDSGLIVFKLE

>UniRef90_A0A7S1TJL4_1_327 | Hypothetical protein n=1 Tax=Erythrolobus australicus TaxID=1077150 RepID=A0A7S1TJL4_9RHOD | E_val=1.2e-160

--MLTKFETKSNRVKGLAFH--PSRPWILSTLHSGSIQLWDYRMGTLID-------RFEE

HEG-PVRGVDFHVT-QPLFVSGGDDYKIKVWNYK--LR-------------RCL-FTLLG

HLDYIRTVYFHKE---HPWIVSASDDQTVRIWNWQ--NRSCLAVLTGHNHYVMCASFHPR

-DDLVVSASLDQTLRVWDI--------S---------AL-SQKS-VRA-P------T---

-----GE---------EM-M--------G-QMRG------------NFNA-A--------

-VNSDLF----------------GGTD--AIVKYVLEGHSRGVNWAHFHPTLPLIISGAD

DRLIKLWRM------------S----------------DVKAWEVDTFRGHLNNVSCVMF

HPRQ-ELVLSDSEDKTIRVWDLNRRTCIYSFRR-EN----------DRFWIMAAHPHVNL

IAAGHDSGMVVFKLE

>UniRef90_A0A7S3GXJ7_4_326 | Hypothetical protein n=1 Tax=Spumella elongata TaxID=89044 RepID=A0A7S3GXJ7_9STRA | E_val=1.7e-160

--MLTKFESKSNRVKGLSFH--PIRPWILASLHNGHIQLWDYRMGTLLD-------KFEE

HDG-PVRGVDFHRT-QPLIVSGGDDYKVKVWDYK--LR-------------RCL-FTLLG

HLDYIRTVQFHVE---YPWIVSASDDQTIRIWNWQ--SRHCISVLTGHNHYVMCAAFHPK

-EDMIVSASLDQTVRVWDT--------T---------GL-RKKT-V-R-G------A---

-----PP---------QM-D--------D-GNV---------------VS-R--------

-VNNELF----------------GGND--AVVKYVLEGHERGVNWASFHPTLPLVISGAD

DRQVKLWRM------------N----------------ETKAWEVDTMRGHTNNVSCVLF

HPKH-ELIVSNSEDRTIRVWDISKRLGVQTFRR-DG----------DRFWILAVHPEQNL

LAAGHDSGMTVFKLE

>UniRef90_U5EW24_1_322 | Coatomer subunit alpha n=1 Tax=Corethrella appendiculata TaxID=1370023 RepID=U5EW24_9DIPT | E_val=2.3e-160

--MLTNFETKSARVKGLSFH--PKRPWILVSLHSGIIQLWDYRISTLIE-------KFDE

HDG-PIRGIAFHAQ-QPLFVSGGDDFKIKVWNYK--QR-------------RCI-FTLHG

HLDYVRTTVFHHE---YPWILSASDDQTIRIWNWQ--SRSCISVLTGHNHYVMCAQFHAT

-DDIVVSASLDQTVRIWDI--------S---------GL-RKKN-V-A-P------G---

-----PN---------GL-D--------E-HL----------------KN-P--------

-TATDLF----------------GQAD--AVVRHVLEGHDRGVNWASFHPNLPLIVSGAD

DRQIKLWRM------------N----------------EYKAWEVDTCRGHYNNVSCVLF

HPRQ-ELILSNSEDKSIRVWDMTKRQCLHTFRR-EH----------ERFWILAAHPHLNL

FAAGHDSGMIVFKLE

>UniRef90_A0A421JR32_1_329 | Coatomer subunit alpha n=1 Tax=Spathaspora sp. JA1 TaxID=2028339 RepID=A0A421JR32_9ASCO | E_val=4.1e-160

MKMLTKFESKSSRAKGIAFH--PKRPWVLVSLHSSTIQLWDYRMGTLID-------RFED

HVG-PVRSVDFHPT-QPLFVSGGDDYSIKVWSLV--TR-------------KCI-FTLNG

HLDYIRQVSFHHD---LPWILSCSDDQTIRIWNWQ--NRQEIACLTGHNHYVMSSQFHPS

-EDLIVSASLDQTVRVWDI--------S---------GL-RKKH-S-AGP------G---

-----SST-------RSF-E--------D-QLQR--------------QQ-L--------

-PQQDIF----------------GNIN--AVVKYVLEGHDKGVNFASFHPTLPLIVSAGD

DRVVKLWRM------------S----------------DTKAWEVDTCRGHTGNVLCAIF

HPHQ-DLILSISDDKTIRVWDLNKRVPVKQFRR-DH----------DRFWLIGAHPNMNL

FGACHDSGVMVFKLE

>UniRef90_A0A1E4SPV1_1_328 | Coatomer subunit alpha n=1 Tax=Suhomyces tanzawaensis NRRL Y-17324 TaxID=984487 RepID=A0A1E4SPV1 | E_val=7.5e-160

MKMLTKFESKSSRAKGVAFH--PKRPWVLVSLHSSTIQLWDYRMGTLID-------RFEE

HTG-PVRCVDFHPT-QPLFVSGGDDYLIKVWSLK--DR-------------LCI-FTLQG

HLDYIRTVSFHNE---LPWIISCSDDQTIRIWNWQ--NQQEIACLTGHNHYVMSAQFHPT

-DDLIVSASLDQTVRVWDI--------S---------GL-RKKH-S-A-P------T---

-----SAA-------RSF-E--------D-QLQR--------------QQ-L--------

-PQQDIF----------------GNVN--AIVKYVLEGHDKGVNWATFHPTLPLIVSAGD

DRLVKLWRM------------S----------------DTKAWEVDTCRGHTGNVLSAVF

HPQQ-DLILSVSDDKTIRVWDLNKRVPIKQFRR-EH----------DRFWLITPHPTMNL

FATCHDSGVMVFKLE

>UniRef90_B3RTT6_1_322 | Coatomer subunit alpha n=2 Tax=Trichoplax TaxID=10227 RepID=B3RTT6_TRIAD | E_val=9.2e-160

--MLIKFETKSARVKGISFH--PMRPWVLASLHSGLIQLWDYRMCTLID-------KYDE

HDG-PVRGVDFHSQ-QPLFVSGGDDYKIKVWNYK--SK-------------KCL-FTLLG

HLDYIRTTFFHNE---YPWIVSSSDDQTIRIWNWQ--SRSCVSVLTGHNHYVMCANFHPT

-EDLIVSASLDQTVRVWDI--------T---------GL-RKKT-V-A-P------G---

-----AG---------GF-D--------D-RN----------------RG-P--------

-GSTDLF----------------GVQD--AVVKHVLEGHDRGVNWANFHHSMPLIVSGAD

DRQVKIWRM------------N----------------DSKAWEVDTCRGHYNNVSCVLF

HPRQ-DLIISNSEDKSIRVWDMSKRIGIQTFRR-ET----------DRFWVVTSHPSLNL

FAAGHDGGLIVFKLE

>UniRef90_G3AYT5_1_328 | Coatomer subunit alpha n=1 Tax=Candida tenuis (strain ATCC 10573 / BCRC 21748 / CBS 615 / JCM 9827 / | E_val=1.3e-159

MKMLTKFESKSSRAKGVALH--PKRPWVLVSLHSSTIQLWDYRMGTLID-------RFED

HSG-PVRCVSFHPT-QPLFVSGGDDYSIKVWSLN--SR-------------KCI-FTLNG

HLDYLRSVSFHHD---LPWILSCSDDQTIRIWNWQ--NRQEIACLTGHNHYVMSAQFHPK

-EDLIVSASLDQTVRVWDI--------S---------GL-RKKH-S-A-P------T---

-----SSI-------RSF-E--------D-QLQR--------------QQ-L--------

-PQQDIF----------------GNVN--AVVKFVLEGHDKGVNYAAFHPTLPLIVSGGD

DRLVKLWRM------------S----------------ETKAWEVDSCRGHTGTVLATIF

HPHQ-DLILSVGDDKTIRVWDLNKRTPVKQFRR-EH----------DRFWDIACHPTVNL

FAGCHDSGVMIFKLE

>UniRef90_F0YGR3_1_324 | Coatomer subunit alpha n=1 Tax=Aureococcus anophagefferens TaxID=44056 RepID=F0YGR3_AURAN | E_val=2.6e-159

--MLTKFESKSNRVKGLAFH--PHRPWILTSLHNGVIQLWDYRMGTLLD-------RFDE

HDG-PVRGVDFHQA-QPLIVSGGDDYRIKVWDYK--LR-------------RCL-FTLLG

HLDYIRTVCFHGD---YPWLVSASDDQTIRIWNWQ--SRSCVSVLTGHNHYVMCASFHTR

-DDLIVSASLDQTVRVWDI--------T---------GL-RKKN-V-R-G------A---

-----PT---------SGAG--------T-ASV---------------VS-R--------

-VNADLF----------------GGND--AVVKYVLEGHDRGVNWASFHPTLPLIISGAD

DRQVKLWRM------------N----------------ETKAWEVDTMRGHTNNVSCVVF

HPKH-ELIISNSEDRSIRVWDISKRLGVQTFRR-EN----------DRFWILAAHPEQNL

LAAGHDSGMIVFKLE

>UniRef90_A0A7S2RES0_1_324 | Hypothetical protein n=1 Tax=Rhizochromulina marina TaxID=1034831 RepID=A0A7S2RES0_9STRA | E_val=4.8e-159

--MLTKFESKSNRVKGLSFH--PVRPWVLTSLHNGVIQLWDYRMGVLLE-------RFDE

HDG-PVRGVDFHKI-QPLIVSGGDDYKIKVWDYK--LR-------------RCR-FTLLG

HLDYIRTVQFHNE---YPWIVSASDDQTIRIWNWQ--SRSCLSVLTGHNHYVMCAAFHPK

-NDLIVSASLDQTVRVWDT--------A---------NL-RKKN-V-R-G------A---

-----PS---------FESG--------P-DTV---------------VS-R--------

-VNADLF----------------GSAD--ALVKYVLEGHDRGVNWASFHPTLPLVVSGAD

DRQVKLWRM------------N----------------ETKAWEVDTLRGHTNNVSCVIF

HPKH-ELIISNSEDRSIRVWDISRRMGIQTFRR-EN----------DRFWILAAHPEQNL

LAAGHDTGMIVFKLE

>UniRef90_I0Z956_1_320 | Coatomer subunit alpha n=1 Tax=Coccomyxa subellipsoidea (strain C-169) TaxID=574566 RepID=I0Z956_COC | E_val=1e-158

--MLTKFETKSNRVKGLSFH--PKRPWILASLHSGVIQLWDYRMGTLID-------RFDE

HDG-PVRGVHFHKS-QPLFVSGGDDYKIKVWNYK--LR-------------RCL-FTLLG

HLDYIRTVQFHQE---YPWVVSASDDQTIRIWNWQ--SRTCISVLTGHNHYVMSACFHPK

-DDLVVSASLDQTVRVWDI--------S---------GL-RKKT-V-A-P----G-G---

-------------------E--------D-MLR---------------LP-Q--------

-MNSDLF----------------GGGD--AVVKYVLEGHDRGVNWAAFHPTLPLIVSGAD

DRQVKLWRM------------N----------------DTKAWEVDTLRGHVNNVSCVMF

HARQ-DIIVSNSEDKSIRVWDMSKRTGVQTFRR-EH----------DRFWIMAAHSEVNL

LAAGHDSGMIVFKLE

>UniRef90_A0A2P6MPH4_1_323 | Coatomer subunit alpha n=1 Tax=Planoprotostelium fungivorum TaxID=1890364 RepID=A0A2P6MPH4_9EUKA | E_val=1.6e-158

--MLIKLETKSNRVKGLSFH--PTRPWVLASLHNGVIQLYDYRIRTLVD-------KFEE

HDG-PVRGLDFHST-QPLFVSGGDDYKIKVWNYK--QR-------------RCL-YPLLG

HLDYIRTVQFHLE---HPWILSSSDDQTIRIWNWQ--SKTCVAVLTGHNHYVMCAAFHPT

-DDLVVSASLDQTVRVWDI--------S---------GL-RKKN-V-SGP------G---

-----TSS-------NSY-E--------D-------------------NR-L--------

-AQNDLF----------------GNAD--ALVKYVLEGHERGVNWVAFHKTAPLIVSGAD

DRQIKLWRM------------N----------------ESRAWEVDSFRGHFNNVSCVLF

HPKQ-DLILSDSEDKTVRVWDTNKRSGIQTFRR-EH----------DRFWILAAHPENNF

FAAGHDSGMLVFKLE

>UniRef90_A0A182F496_1_323 | Coatomer subunit alpha n=5 Tax=Nyssorhynchus TaxID=44543 RepID=A0A182F496_ANOAL | E_val=2.9e-158

--MLTNFETKSARVKGLSFH--PKRPWILASLHSGVIQLWDYRISTLIE-------KFDE

HDG-PVRGIAFHNQ-QPLFVSGGDDFKIKVWNYK--QR-------------RCI-FTLLG

HLDYVRTTVFHHE---YPWILSASDDQTIRIWNWQ--SRSCICVLTGHNHYVMCAQFHMS

DEDIIVSASLDQTVRIWDI--------S---------GL-RKKN-V-A-P------G---

-----PA---------GL-E--------D-HL----------------KN-P--------

-GTADLF----------------GQAD--AVVKHVLEGHDRGVNWACFHPSMPLIASGAD

DRQVKLWRM------------N----------------EYKAWEVDTCRGHYHNVSCVLF

HPRA-DFIISNSEDKSIRVWDMTKRQCIHTFRR-EN----------ERFWILAAHPNLNL

FAAGHDSGTIVFKLE

>UniRef90_A0A6A5PKY2_1_319 | Coatomer subunit alpha n=3 Tax=Lupinus albus TaxID=3870 RepID=A0A6A5PKY2_LUPAL | E_val=4.3e-158

--MLTKFETKSNRVKGLSFH--IKRPWILASLHSGVIQLWDYRMGTLID-------RFDE

HDG-PVRGVHFHNS-QPLFVSGGDDYKIKVWNYK--TH-------------RCL-FTLLG

HLDYIRTVQFHHE---SPWIVSASDDQTIRIWNWQ--SRTCISVLTGHNHYVMCALFHPK

-EDLVVSASLDQTVRVWDI--------S---------SL-KRKN-A-S-P------A---

-------------------D--------D-VLR---------------LS-Q--------

-MNTDLF----------------GGVD--AVVKYVLEGHDRGVNWASFHPTLPLIVSGAD

DRQVKLWRM------------N----------------DTKAWEVDTLRGHMNNVSCVMF

HAKQ-DIIVSNSEDKSIRVWDATKRTGIQTFRR-EH----------DRFWILVAHPELNL

LAAGHDSGMIVFKLE

>UniRef90_A0A1E3NR24_1_335 | Coatomer subunit alpha n=2 Tax=Pichia membranifaciens TaxID=4926 RepID=A0A1E3NR24_9ASCO | E_val=5.9e-158

MKMLTKFESKSSRAKGVAFH--PRRPWLLVSLHSSTIQLWDYRMGTLID-------RFED

HDG-PVRCVAFHPT-QPIFVSGGDDYTIKVWSTQ--TR-------------KCM-FTLTG

HLDYVRTVFFHHE---LPWIISASDDQTIRIWNWQ--NRKEIACLTGHNHYVMCAQFHPK

-QDLIVSASLDQTVRVWDI--------S---------GL-VKKH-S-A-P------S---

-----GAS--------SF-GGHSSMNQFD-QYNR--------------NQ-P--------

-PQQDIF----------------GNTD--AVVKYVLEGHDKGVNWASFHPELPLIVSGGD

DRAVKLWRM------------S----------------ETRAWEVDGCRGHTNNVPCVLF

HPTE-DLIISVGEDKTIRTWDLNSRSPVKQFKR-EN----------DRFWMIVAHPNINL

FAACHDSGVMVFKLD

>UniRef90_A0A7S1XAY7_1_331 | Hypothetical protein n=1 Tax=Compsopogon caeruleus TaxID=31354 RepID=A0A7S1XAY7_9RHOD | E_val=1.2e-157

--MLTKFETKSNRVKGLSFH--HFRPWVLASLHNGSIQMWDYRMGTLLD-------RYDE

HDG-PVRGVEFHRS-QPLFVSGGDDYKIKLWNYK--LR-------------RCV-FTLLG

HLDYIRTVSFHHE---SPWIVSASDDQTVRIWNWQ--NRQCLAVMTGHNHYVMSAAFHPA

-EDLVVSASLDQTIRVWDI--------S---------EL-RTGG-N-R-P------F---

-----PSS-------KGP-Q--------D-VLSK---------VKGKLPP-S--------

-VNADLF----------------GTSD--AVVKYVLEGHSRGVNWATFHPTLPLIISGAD

DRQVKLWRM------------S----------------DTKAWEVDTFRGHFNNVSSAIF

HPHR-ELVISCSEDKTIKVWDLARRICVHTFRR-EN----------DRFWISAAHPRINL

LAAGHDSGMVVFKLE

>UniRef90_A0A2K1JUI4_1_319 | Coatomer subunit alpha n=2 Tax=Physcomitrium patens TaxID=3218 RepID=A0A2K1JUI4_PHYPA | E_val=1.4e-157

--MLTKFETKSNRVKGLSFH--PRRPWILASLHSGIIQLWDYRMGTLID-------RFDE

HDG-PVRGVHFHKS-QPLFVSGGDDYKIKVWNYK--MR-------------RCL-FTLLG

HLDYIRTVQFHHE---NPWIVSASDDQTIRIWNWQ--SRTCISVLTGHNHYVMCASFHIK

-EDLVVSASLDQTVRVWDI--------G---------AL-RKKS-V-A-P------A---

-------------------V--------D-MLR---------------LT-Q--------

-MNTDLF----------------GGGD--SVVKYVLEGHDRGVNWASFHPSLPLIVSGAD

DRQVKLWRM------------N----------------DTKAWEVDTLRGHVNNVSCVMF

HARQ-DIIVSNSEDKSIRVWDMSKRTGVQTFRR-EQ----------DRFWILSAHPEMNL

LAAGHDSGMIVFKLE

>UniRef90_A0A166GFB3_1_319 | Coatomer subunit alpha n=1 Tax=Daucus carota subsp. sativus TaxID=79200 RepID=A0A166GFB3_DAUCS | E_val=3.5e-157

--MLTKFETKSNRVKGLSFH--SKRPWILASLHSGVIQLWDYRMGTLID-------RFDE

HDG-PVRGVHFHKS-QPLFVSGGDDYKIKVWNYK--LH-------------RCL-FTLLG

HLDYIRTVQFHHE---HPWIVSASDDQTIRIWNWQ--SRTCISVLTGHNHYVMCALFHPK

-DDLVVSASLDQTVRVWDI--------G---------AL-RKKS-A-S-P------A---

-------------------D--------D-ILR---------------LS-Q--------

-MNTDFF----------------GGVD--AVVKYVLEGHDRGVNWAAFHPSLPLIVSGAD

DRQVKLWRM------------N----------------DTKAWEVDTLRGHMNNVSCVLF

HAKQ-DIIVSNSEDKSIRVWDATKRTAIQTFRR-EQ----------DRFWILASHPEMNL

LAAGHDSGVIVFKLE

>UniRef90_A0A1X6NW10_1_324 | WD_REPEATS_REGION domain-containing protein n=1 Tax=Porphyra umbilicalis TaxID=2786 RepID=A0A1X6 | E_val=6.7e-157

--MLTKFETKSNRVKGLAFH--PQRPWVLSSLHNGAIQLWDYRMGTLID-------RYDE

HEG-PVRSVDFHPT-QPLFVSGGDDYKIKVWNYK--LR-------------RCV-FTLYG

HLDYIRTVSFHPE---SPWIVSASDDQTVRIWNWQ--NRSCLAVLSGHNHYVMSASFHPA

-EDLVVSASLDLTVRVWDI--------S---------AL-RQKA-V-R-P------S---

-----DMI-------SSV-R--------G-QL----------------PP-S--------

-VNADLF----------------GTMD--ATVRHVLEGHTRGANWASFHPTLPLIVSGGD

DRHVKLWRM------------S----------------DTKAWEMDTLRGHLNNVSCTLF

HPRA-DLVLTASEDKTIRVWDLNRRSCIHTFRR-EA----------DRFWILAAHPTVNL

LAAGHDNGMLVFKLQ

>UniRef90_A0A218XY20_1_319 | Coatomer subunit alpha n=2 Tax=Punica granatum TaxID=22663 RepID=A0A218XY20_PUNGR | E_val=1.6e-156

--MLTKFETKSNRVKGLSFH--SKRPWILASLHSGVIQLWDYRMGTLID-------RFDE

HDG-PVRGVHFHKS-QPLFVSGGDDYKIKVWNYK--TC-------------RCL-FTLLG

HLDYIRTVQFHHE---HPWIVSASDDQTIRLWNWQ--SRTCISVLTGHNHYVMCASFHPK

-EDLVVSASLDQTVRVWDI--------S---------SL-KKKT-A-S-R------G---

-------------------D--------D-ILR---------------LS-Q--------

-MNTDLF----------------GGVD--VVVKYVLEGHDRGVNWAAFHPNLPLVVSGAD

DRQVKLWRM------------N----------------DAKAWEVDTLRGHTNNVSCVMF

HARQ-DIIVSNSEDKSIRVWDATKRTGLQTFRR-EH----------DRFWILTAHPEMNL

FAAGHDSGMIVFKLE

>UniRef90_A0A4P9ZEH1_1_328 | Coatomer subunit alpha n=1 Tax=Metschnikowia bicuspidata TaxID=27322 RepID=A0A4P9ZEH1_9ASCO | E_val=2.9e-156

MKMLTKFESKSLRAKGVAFH--PTRPWVLVALHLSIIQLWDYRMGTLLD-------RFED

HEG-PVRCVNFHPT-QPLFVSGSDDYFIKVWSLT--TR-------------KCI-FTLTG

HLDYLRNVSFHHD---LPWVLSCSDDQTIRIWNWQ--NRQEIACLTGHNHYVLLAQFHPS

-DDLIVSASLDLTVRVWDI--------L---------GL-RKKH-S-A-P------L---

-----SQQ-------HLF-E--------D-QMLR--------------RQ-L--------

-PQQDIF----------------GNVN--AVVKYVLEGHDKGVNWAAFHPTLPLIVSAAD

DRLVKLWRM------------S----------------DTKAWEVDTCRGHTGNVICAVF

HPRQ-DLILSVSDDRTIRVWDLNKRTPVKQFKR-EN----------DRFWMVAPHRSINL

FAACHDSGVIVFKLE

>UniRef90_A0A1I8BXQ7_1_332 | Coatomer subunit alpha n=2 Tax=Meloidogyne TaxID=189290 RepID=A0A1I8BXQ7_MELHA | E_val=4.7e-156

MALLKKFESKSARVKGISFH--PTRPWVLASLHSGVIQLWDYRMCVLID-------KFDE

HDG-PVRGVDFHSQ-QPIFVSGGDDYKIKVWNYK--QR-------------KCI-FTLLG

HLDYIRTTYFHKQ---HPWIISASDDQTVRIWNWQ--SRISIAILTGHNHYVMCAQFHPT

-EDLVASASLDQTVRIWDI--------S---------GL-RKKN-S-A-P------G---

-----GGS--------SA-P--------S-GISR---------FGSTSVS-S--------

-AQTELF----------------GQPD--VVVKHVLEGHDRGVNWVSFHPSMPLLVSGAD

DRQVKLWRY------------N----------------DSKAWEVDSCRGHYNNVSCVIF

HPKA-ELILSNSEDKSIRVWDMQKRTCLQNFRH-EN----------DRFWVMSSHPSLNL

FAAGHDNGMIVFKTE

>UniRef90_F6GZQ1_1_319 | Coatomer subunit alpha n=4 Tax=Vitis TaxID=3603 RepID=F6GZQ1_VITVI | E_val=7.4e-156

--MLTKFETKSNRVKGLSFH--TKRPWILASLHSGVIQLWDYRMGTLID-------RFDE

HDG-PVRGVHFHKS-QPLFVSGGDDYKIKVWNYK--LH-------------RCL-FTLFG

HLDYIRTVQFHHE---YPWIVSASDDQTIRIWNWQ--SRTLMSVLTGHNHYVMCASFHPK

-EDLVVSASLDQTVRVWDI--------G---------AL-RKKT-S-S-P------A---

-------------------D--------D-ILR---------------LS-Q--------

-MNTDFF----------------GGVD--AVVKYVLEGHDRGVNWASFHPTLPLIVSGAD

DRQVKLWRM------------N----------------DTKAWEVDTLRGHMNNVSCVFF

HARQ-DVIVSNSEDKSIRVWDATKRTGIQTFRR-EH----------DRFWILTAHPEMNL

LAAGHDSGMIVFKLE

>UniRef90_A0A1L8DVX3_1_331 | Coatomer subunit alpha n=3 Tax=Phlebotominae TaxID=7198 RepID=A0A1L8DVX3_9DIPT | E_val=1e-155

--MLTNFETKSARVKGLSFH--PKRPWVLASLHNGVIQLWDYRMHTLLE-------KFDE

HDG-PVRGICFHSQ-QPLFVSGGDDFKIKVWNYK--QR-------------RCT-FTLLG

HLDYVRTTVFHEE---YPWILSASDDQTIRIWNWQ--SRQCISVLTGHNHYVMCALFHPT

-EDQLVSASLDQTVRVWDI--------SGTANRDGMKGL-RKKN-V-A-P------G---

-----PS---------GL-E--------E-HL----------------KN-P--------

-GATDLF----------------GQAD--AVVKHVLEGHDRGVNWASFHPHMPLIVSGAD

DRQIKLWRM------------N----------------EYKAWEVDTCRGHYNNVSCVLF

HPRQ-ELIISNSEDKSIRVWDMTKRQCLHTFRR-EH----------ERFWVLAAHPTLNL

FAAGHDAGMIVFKLE

>UniRef90_A0A642UM92_1_334 | Coatomer subunit alpha n=1 Tax=Diutina rugosa TaxID=5481 RepID=A0A642UM92_DIURU | E_val=1.1e-155

MKMLTKFESKTSRAKGVAFH--PKRPWALVSLHSSTIQLWDYRMGTLID-------RFDD

HTG-PVRCVDFHPT-QPLFVSGGDDYTVKVWSLN--TR-------------KCI-FTLNA

HLDYVRTVFFHKT---LPWIISCSDDLTIKIWNWQ--NRQEIACLTGHNHYVMSAQFHPT

-EDLIVSASLDQTVRVWDI--------S---------GL-RKKH-S-A-P------GGGM

GPG--SGP-------RSF-E--------D-QLTR--------------SQ-L--------

-PQQDIF----------------GNAN--AVVKYVLEGHDKGVNWAAFHPTLPLIVSAAD

DRLVKLWRM------------S----------------DTKAWEVDTCRGHTGNVLCSIF

HPTQ-DLILSVSDDKTVRVWDLNKRTPIKQFRR-EH----------DRFWLIANHPSINL

FAACHDSGVMIFKLE

>UniRef90_A0A5J4N8Q7_14_352 | Coatomer subunit alpha (Fragment) n=1 Tax=Paragonimus westermani TaxID=34504 RepID=A0A5J4N8Q7_9T | E_val=1.7e-155

MSYLSKFDCKSARVKGLTFH--HKRPWVLSSLHTGIIQLWDYRSCTLID-------KFEG

HEG-PVRGIDFHNN-QPLFVSGGDDYKIKVWNYK--QR-------------KCL-FNLLG

HLDYIRTTFFHKE---YPWILSASDDQTIRIWNWQ--SRTIASVLTGHSHYVMCAQFHPK

-EDLVVSASLDQTVRVWDI--------S---------GL-RKKN-V-A-P------S---

-----GIS--------GI-E--------D-HMRQLTGSSR--HGNTGVGG-P--------

-GHTELF----------------GTGD--VVVRHVMEGHDRGVNWVAFHPTLPIVVSAAD

DRLVKLWRM------------T----------------ETKAWELDTLRGHYNNVSCVLF

HPRQ-DLLLSDSEDKSIRIWDLAKRTCVTTIRR-DS----------DRFWVVAAHPKLNL

FAAGHDTGFVVFKLE

>UniRef90_A0A1Y1WGI0_1_322 | Coatomer subunit alpha n=1 Tax=Linderina pennispora TaxID=61395 RepID=A0A1Y1WGI0_9FUNG | E_val=2.3e-155

MQMLTKFETKSSRVKGVAFH--PKRPWILASLHNGSIQLWDYRMGTLLE-------RFEE

HEG-PVRGISFHPS-QELFVSGGDDYKVKVWNYR--TR-------------RCL-FTLQG

HLDYVRTAR---------WIITASDDQTIRIWNWQ--SRQCIAVLTGHNHYVMSAEFHPT

-EDLVVSACLDQTVRVWDI--------S---------GL-RQKS-A-A-G------A---

-----QPV--------MM-A--------D-PLAS-------------QRM-G--------

-AQGDFF----------------NATD--VVVKFVLEGHTRGVNWATFHPTMPLILSCGD

DRQIKIWRM------------N----------------DSRAWEVDTCRGHYNNVNSALF

HPKR-EFILSDSEDKTIRVWDTTRRTLLATFRR-EQ----------DRFWCLTAHPELNL

FAAGHDNGLIVFKLE

>UniRef90_A0A1Q3ATJ7_1_320 | Coatomer subunit alpha n=1 Tax=Cephalotus follicularis TaxID=3775 RepID=A0A1Q3ATJ7_CEPFO | E_val=2.9e-155

--MLTKFETKSNRVKGLSFH--SKRPWILTSLHSGVIQLWDYRMGTLID-------RFDE

HDG-PVRGVHFHNS-QPLFVSGGDDYKIKVWNYK--LH-------------RCL-FTLLG

HLDYIRTVQFHHE---SPWIVSASDDQTIRLWNWQ--SRTCISVLTGHNHYVMCASFHPK

-EDLVVSASLDQTVRVWDI--------G---------PLKKKKT-A-S-P------A---

-------------------D--------D-ILR---------------MS-Q--------

-MNTDFF----------------GGVD--AVVKYVLEGHDRGVNWAAFHPTLPLIVSGAD

DRQLKLWRM------------N----------------DTKAWEVDTLRGHTNNVSCVMF

HAKQ-DIIVSNSEDKSIRVWDATKRTGIQTFRR-EH----------DRFWILACHPEMNL

LAAGHDSGMIVFKLE

>UniRef90_S8CG48_1_319 | Coatomer subunit alpha n=1 Tax=Genlisea aurea TaxID=192259 RepID=S8CG48_9LAMI | E_val=3.8e-155

--MLTKFETKSNRVKGLSFH--AKRPWILASLHSGVIQLWDYRMGTLID-------RFDE

HDG-PVRGVHFHKS-QPLFVSGGDDYKIKVWNYK--LH-------------RCL-FTLLG

HLDYVRTVQFHHE---YPWIVSASDDQTIRIWNWQ--SRTCISVLTGHNHYVMSASFHPK

-EDLVVSASLDQTVRVWDI--------G---------AL-RKKT-V-S-P------A---

-------------------D--------E-VIR---------------LP-Q--------

-MNTDFF----------------GGVD--AVVKYVLEGHERGVNWASFHPTLPLIVSGAD

DRHVKIWRM------------N----------------DTKAWEVDTLRGHMNNVSCVLF

HAKQ-DVIVSNSEDKSIRIWDSTKRTGLQTFRR-EH----------DRFWILSAHPEMNL

LAAGHDNGIMVFKLE

>UniRef90_W0TI75_1_324 | Coatomer subunit alpha n=1 Tax=Kluyveromyces marxianus (strain DMKU3-1042 / BCC 29191 / NBRC 104275) | E_val=7e-155

MKMLTKFESKSTRAKGIAFH--PSRPWVLVALFSSTIQLWDYRMGVLLH-------RFED

HEG-PVRGIDFHPT-QPLFVSAGDDYTIKVWSLE--TN-------------KCL-FTLDG

HLDYVRTVFFHHE---LPWIISSSDDQTIRIWNWQ--NRKEIACLTGHNHFVMCAQFHPV

-EDLVVSASLDETVRVWDI--------S---------GL-RKRH-S-A-P------G---

-----SQ---------SF-D--------E-QM-------------------R--------

-QQQNLLD--------------GGFGD--CVVKFILEGHTRGVNWASFHPTLPLIVSGSD

DRQVKLWRM------------S----------------ATKAWEVDTCRGHTNNVDSVIF

HPYQ-NLIISVGEDKTLRVWDLDKRTPVKQFKR-EN----------DRFWLVRAHPNLNL

FGAAHDSGIMIFKLD

>UniRef90_A0A1W0WQG3_1_321 | Coatomer subunit alpha n=1 Tax=Hypsibius dujardini TaxID=232323 RepID=A0A1W0WQG3_HYPDU | E_val=1e-154

--MLTKFECKSARVKGLAFH--PKRPWILSSLHTGVIQLWDYRLNTMIE-------KYEE

HEG-PVRGVHFHSQ-QPLFVSGGDDYKIKVWNYK--LR-------------RCI-FTLLG

HLDYIRTTFFHHE---YPWIITCSDDQTVRIWNWQ--SRNCINILTGHNHYVMCAQFHPT

-EDLVISASLDQTIRVWDI--------S---------GL-RKKN-V-A-P------G---

-----PG---------GM-D----------SV----------------RT-P--------

-GQTELF----------------GQTD--VVVKFVLEGHDRGVNWAMFHPTAKLIVSGSD

DRQVKVWRY------------S----------------QEKAWEIDSCRGHYNNVSCVSF

SNSQ-DVILSNSEDKSIRIWDLNKRTCLYTFRR-EH----------DRFWVIGAHPTLNL

YAAGHDSGCIVFKLE

>UniRef90_A0A2J6K5C6_1_319 | Coatomer subunit alpha n=2 Tax=Lactuca TaxID=4235 RepID=A0A2J6K5C6_LACSA | E_val=1.7e-154

--MLTKFETKSNRVKGLSFH--NKRPWILASLHSGVIQLWDYRMGTLID-------RFEE

HDG-PVRGVHFHQS-QPLFVSGGDDYKIKVWNYK--LH-------------RCL-FTLLG

HLDYIRTVQFHHE---YPWIVSASDDQTIRIWNWQ--SRACISVLTGHNHYVMCALFHPK

-EDLVVSASLDQTIRVWDI--------G---------AL-KKKT-V-S-P------A---

-------------------D--------D-ILR---------------LS-Q--------

-MNADFF----------------GGVD--AVVKYVLEGHDRGVNWASFHPTLPLIVSGAD

DRQVKIWRM------------N----------------DSKAWEVDTLRGHMNNVSSVLF

HSKQ-DIIVSNSEDKSIRVWDATKRTALQTFRR-EH----------DRFWILTCHPELNL

LAAGHDNGMIVFKFE

>UniRef90_A0A016SBB5_39_362 | WD_REPEATS_REGION domain-containing protein n=5 Tax=Ancylostoma TaxID=29169 RepID=A0A016SBB5_9BI | E_val=2.8e-154

MSLLIKFESKSARVKGISFH--PTRPWVLASLHSGVIQLWDYRMCVLLD-------KFDE

HDG-PVRGICFHQD-QPIFVSGGDDYKIKVWNYK--QR-------------RCI-FTLLG

HLDYIRSTFFHHK---YPWIISSSDDQTVRIWNWQ--SRNSIAILTGHNHYVMCAQFHPS

-EDLVASASLDQTVRIWDI--------S---------GL-RKKQ-M-P-G------G---

-----GS---------AV-P--------R----------------SGTGP-Q--------

-SQADLF----------------GQPD--VVVKHVLEGHDRGVNWVAFHHTMPILVSGSD

DRQVKMWRY------------N----------------ESKAWEVDSCRGHYNNVSSVLF

HPNA-ELILSNSEDKSIRVWDMQKRTSLHVFRH-EN----------ERFWVLSAHPNLNM

FAAGHDNGMIVFKI-

>UniRef90_A0A2Z7B4Q9_1_319 | Coatomer subunit alpha n=1 Tax=Dorcoceras hygrometricum TaxID=472368 RepID=A0A2Z7B4Q9_9LAMI | E_val=4.8e-154

--MLTKFETKSNRVKGLSFH--SKRPWILASLHSGVIQLWDYRMGTLID-------RFDE

HEG-PVRGVHFHKS-QPLFVSGGDDYKIKVWNYK--LH-------------RCL-FTLLG

HLDYIRTVQFHHE---CPWIVSASDDQTIRIWNWQ--SRTCISVLTGHNHYVMCASFHPK

-EDLVVSASLDQTVRVWDI--------G---------AL-RKKT-V-S-P------A---

-------------------D--------D-ILR---------------LS-Q--------

-MNADFF----------------GGVD--AVVKYVLEGHDRGVNWASFHPTLPLIVSGAD

DRQVKIWRM------------N----------------DTKAWEVDTLRGHVNNVSCALF

HARQ-DIIVSNSEDRSIRVWDATKRTGLHTFRR-EV----------DRFWILSVHPEINL

LAAGHDSGMIVFKLE

>UniRef90_UPI00187D88BE_1_329 | coatomer subunit alpha isoform X1 n=2 Tax=Bradysia coprophila TaxID=38358 RepID=UPI00187D88BE | E_val=5.5e-154

--MLINFESKSARVKGLSFH--PKRPWILVSLHSGVIQLWDYRMCTLLE-------KFDE

HDG-PVRGICFHNQ-QPLFVSGGDDFKIKFWNYK--KR-------------RCI-FTLHG

HLDYVRTTFFHHE---YPWILSASDDQTIRIWNWQ--SRTCISVLTGHNHYVMCAQFHPT

-EDQIVSASLDQTVRVWDI--------AGIANRDGMKGL-RKKN-V-A-P------G---

-----PG------------D--------D-LP----------------KL-Q--------

-GTTDLF----------------GAAD--AVVKHVLEGHDRGVNWANFHPTLPLIVSGAD

DRQIKLWRM------------N----------------EYKAWEVDTCRGHYNNVSSVMF

HPRQ-ELIISNSEDKSIRVWDMTKRQCLHTFRR-EN----------ERYWILTGHPTLNL

FAAGHDAGMVVFKLE

>UniRef90_UPI000C1D69A6_1_319 | coatomer subunit alpha-1-like isoform X1 n=4 Tax=Olea europaea var. sylvestris TaxID=158386 R | E_val=1e-153

--MLTKFETKSNRVKGLSFH--SKRPWILASLHNGVIQLWDYRMGTLVD-------RFDK

HVG-PVRGVHFHKS-QPLFVSGGDDYKIMVWNYK--LH-------------SCL-FTLLG

HLDYIRTVQFHHE---YPWIVSASDDQTIRIWNWQ--SRTCISVLTGHNHYVMCASFHPK

-EDLVVSASLDQTVRVWDI--------G---------AL-RKKT-V-S-P------G---

-------------------D--------D-ILR---------------LS-Q--------

-MNTDFF----------------GGVD--AVVKYVLEGHDRGVNWVSFHPTLPLIVSGAD

DRQVKIWRM------------N----------------DTKAWEVDTLSGHLNNVSCVSF

HARQ-DIIVSNSEDKSIRIWDATKRTGLQTFRR-EH----------DRFWILSAHPEMNL

LAAGHDSGMIVFKLE

>UniRef90_UPI001263749E_1_322 | coatomer subunit alpha-2-like n=1 Tax=Pistacia vera TaxID=55513 RepID=UPI001263749E | E_val=2.3e-153

--MLTKFETKSNRVKGLSFH--SKRPWILASLHNGVIQLWDYRMGTLVD-------QFDE

HDG-PVRSVQFHRS-QPLFVSGSDDYKIKVWNYK--LH-------------RCL-FSLIG

HLDYIRTVEFHQE---CPWIVSSSDDQTIRIWNWQ--SRTCVSILTGHNHYVTCASFHPK

-DDLIVSASLDQTIRVWDI--------S---------TL-RKNT-V-A-P------S---

-------------------D--------D-VLRS------------KQLS-Q--------

-INSDLF----------------GGVD--AVVKHVLEGHDRGVNWASFHPTLPLIVSAAD

DRQVKIWRM------------N----------------DSKAWEVDTLRGHMNNVSCVLF

HARQ-DIIVSNSEDRSIRVWDAIKRTGTQTFRR-EH----------DRFWILASHPEMNL

LAAGHDSGLIVFKLE

>UniRef90_M0SSF7_1_318 | Coatomer subunit alpha n=1 Tax=Musa acuminata subsp. malaccensis TaxID=214687 RepID=M0SSF7_MUSAM | E_val=3.8e-153

--MLTKFETKSNRVKGLSFH--SKRPWILASLHSGVIQLWDYSMGTLID-------RFDE

HDG-PVRGVHFHKS-QPLFVSGGDDYKIKVWNYK--TR-------------RCL-FTLLG

HLDYIRTVQFHDE---HPWIVSASDDQTIRIWNWQ--SRTCISVLTGHNHYVMCASFHPK

-EDLVVSASLDQTIRVWDI--------G---------SL-RQKV-V---P------A---

-------------------D--------D-ILR---------------LS-Q--------

-MNTDLF----------------GGID--AVVKYVLEGHDRGVNWASFHPTLPLIVSGAD

DRQVKLWRM------------N----------------DSKAWEVDTLRGHTNNVSSVMF

HTKM-DIIVSNSEDKSIRVWDATKRTGVQTFRR-EH----------DRFWILAVHPAMNL

LAAGHDSGMIIFKLE

>UniRef90_A0A1R1YFE8_1_325 | Coatomer subunit alpha n=3 Tax=Smittium culicis TaxID=133412 RepID=A0A1R1YFE8_9FUNG | E_val=1.7e-152

MQMLTKFESKSSRVKSAAFH--PKRPWVLSSLHNGSIQLWDYRMGSLIE-------KFDD

HEG-PVRGVAFHPS-QDLFVSGGDDYKIRVWSYK--SK-------------KCL-FVLEG

HMDYIRTVSFHAE---QPWILSASDDQTVRIWNWQ--SRQCIAVLSGHNHYVMSAQFHPE

-HDLIVSASLDQTVRVWDF--------S---------GL-KKKN-S-A-G------A---

-----PP---------IQ-Q--------N-SM---------------NFN-S--------

-LSQDDL----------------NQSD--VYARFVLEGHTRGVNWVSFHPTKPLIISCGD

DRQIKLWRM------------S----------------ETKAWEVEAFRGHYNNVSAAVF

QSNR-DFIVSCSEDKSVRVWDTNKQSLVQSFKR-EN----------DRFWCITLHPNINL

FAAGHDSGLIVFKLE

>UniRef90_I2H5W3_1_325 | Coatomer subunit alpha n=1 Tax=Tetrapisispora blattae (strain ATCC 34711 / CBS 6284 / DSM 70876 / NB | E_val=3.3e-152

MKMLTKFESKSTRAKGIAFH--PSRPWALVALFSSTIQLWDYRMGTLLH-------RFED

HEG-PVRSVDFHPT-QPLFVSAGDDCTIKVWSLE--TN-------------KCL-YTLTG

HLDYVRTVFFHHE---LPWIISASDDQTIRIWNWQ--NRKEIANLIGHNHFVMCAQFHPT

-EDLVVSASLDETVRVWDI--------S---------GL-RKKH-S-A-P------A---

-----QSA--------SF-E--------E-QM-------------------S--------

-TQQNILD--------------GGFGD--CVVKFILEGHTRGVNWVSFHPTLPLIVSGSD

DRQVKLWRM------------S----------------ATKAWEVDTCRGHTNNVDSVIF

HPHQ-NLIISVGEDKTLRVWDLDKRTPVKQFKR-EN----------DRFWLIAAHPNINL

FGAAHDSGVMIFKLD

>UniRef90_A0A0C3QTC3_1_315 | WD_REPEATS_REGION domain-containing protein n=1 Tax=Tulasnella calospora MUT 4182 TaxID=1051891 | E_val=8.9e-152

MQMLTKFESKSNRVKGLAFH--PTRPLLAASLHNGSVQLWNYQMGTLVD-------RFDE

HDG-PVRGVAIHPT-RPLLVTGGDDYKVKVWDIRPQGR-------------KCL-FTLHG

HLDYVRTVQFHHE---MPWIISASDDQTIRIWNST--SRNCIAILTGHSHYIMSAMFHPK

-EDLVVSASMDQTVRVWDI--------S---------GL-RKGT-P-Q-A----------

-------------------------------------------------Q-P--------

-GAFDTF-----------------DSF--STVKYVLEGHDRGVNYAAFHPSLPLIVSAGD

DRQVKLWRM------------S----------------ETKAWEVDTCRGHFNNVSSALF

HPRH-ELILSVGEDKTIRVWDMSRRTAVQTFRR-EH----------DRFWVLTAHPELNL

FAAGHDNGLIVFKLE

>UniRef90_A0A7J7MN17_1_335 | Coatomer subunit alpha n=1 Tax=Kingdonia uniflora TaxID=39325 RepID=A0A7J7MN17_9MAGN | E_val=2e-151

--MLTKFETKSNRVKGLSFH--TKRPWILASLHSGVIQLWDYRMGTLID-------RFDE

HDG-PVRGVHFHKT-QPLFVSGGDDYKIKVWNYK--MH-------------RCL-FTLLG

HLDYIRTVQFHHE---YPWIVSASDDQTIRIWNWQ--SRNCISVLTGHNHYVMCATFHPK

-EDLVVSASLDQTVRVWDI--------G---------AL-RKKT-V-S-P------A---

-------------------D--------D-ILR---------------LS-Q--------

-MNTDLF----------------GGVD--AIVKYVLEGHDRGVNWASFHPSLPLIVSGAD

DRQVKLWRM------------NGISEKPVSCVIWQYPIDTKAWEVDTLRGHMNNVSCVMF

HAKQ-DIIVSNSEDKSIRVWDVTKRTGVQTFRR-EH----------DRFWVLASHPEMNL

LAAGHDSGMIVFKLE

>UniRef90_Q6C5A1_9_355 | Coatomer subunit alpha n=4 Tax=Yarrowia lipolytica TaxID=4952 RepID=Q6C5A1_YARLI | E_val=6.4e-151

--ILTQFESKSSRAKGLAFH--STRPWVLVSLHSSTIQLWDYRMGTLVD-------RFED

HDG-PVRGVDFHKT-QPLFVSCGDDYKIKVWSLQ--TR-------------KCL-FTLVG

HLDYVRTVFFHHE---LPWIISCSDDQTIRIWNWQ--NRQEIACLTGHSHYIMSAQFHPS

-EDLVVSACLDQTVRVWDI--------S---------GL-RKKH-S-A-G------G---

-----GVSAGGAGSSMSF-E--------E-QMMMAARNSGGPGGPGGHPQ-Q--------

-GGQDMF----------------GNQD--CIVKYVLEGHDGGVNWATFHPTLPLIVSGGD

DRVLKIWRM------------S----------------DTKAWEVDTCRGHTNNILSCCF

HPYQ-DVIVSVSEDKTIRTWDLHKRTLIKQFKR-EN----------DKFWALTAHPNINL

FAAGHESGIMVFKME

>UniRef90_A0A0K0EKX4_19_349 | Coatomer subunit alpha n=4 Tax=Strongyloides TaxID=6247 RepID=A0A0K0EKX4_STRER | E_val=2.2e-150

MALLKRFETQSARVKGISFH--ATRPWILAALHSGVIQLWDYRMCTMVD-------KFEE

HDG-PVRGIDFHDQ-QPIFVSGGDDYKIKVWNYK--QR-------------KCL-FTLLG

HLDYIRTTFFHKN---YPWIISASDDQTIRIWNWQ--SRNSIAILTGHNHYVMCAQFHPT

-EDIVASASLDQTIRIWDI--------S---------GL-KKKN-A-P-P------G---

-----MGG--------SM-P--------S-RVGS----------GAGPLS-A--------

-AQGDLF----------------GQPD--ILVKHVLEGHDRGVNWVSFHQSLPLLVSGAD

DKQVKLWRY------------N----------------DVRAWEVDSCRGHYNNISSVVF

HPKA-ELILSNSEDKSIRVWDMQKKTCLHTFRH-DN----------DRFWILANHPSLNI

FAAGHDNGLIVFKIE

>UniRef90_S8EE05_10_330 | WD_REPEATS_REGION domain-containing protein (Fragment) n=1 Tax=Genlisea aurea TaxID=192259 RepID=S8E | E_val=6.2e-150

-EMLTKFETKSNRVKGLSFH--SKRPWILASLHSGVIQLWDYRMGTRID-------QFDE

HAGVPVRGVHFHKS-QPLFVSGGDDFKIKVWNYK--LH-------------RCL-FTLLG

HLDYVRTVQFHHE---YPWIVSASDDQTIRIWNWQ--SRACISVLTGHNHYVMSASFHPK

-EDLVVSASLDQTVRVWDI--------G---------AL-RKKT-V-S-P------S---

-------------------E--------D-FIR---------------LP-Q--------

-MNTDFF----------------GGID--AVVKYVLEGHDRGVNWASFHPTLPLIVSGAD

DRQVKIWRM------------N----------------DTKAWEVDTLRGHMNNVSCVLF

HAKQ-DIIVSNSEDKSIRVWDSTKRTGLQNFRR-EH----------DRFWILAAHPEMNL

LAAGHDSGMIVFKLE

>UniRef90_Q75A42_1_324 | Coatomer subunit alpha n=2 Tax=Saccharomycetaceae TaxID=4893 RepID=Q75A42_ASHGO | E_val=1.3e-149

MKMLTKFESKSTRAKAIAFH--PSRPWVLVALFSSTIQLWDYRMGVLLH-------RFEE

HEG-PVRGVDFHPT-QPLFVSAGDDYSIKVWSLS--TH-------------KCL-FTLNG

HLDYVRTVFFHTE---LPWIISASDDQTIRIWNWQ--NRREIACLTGHNHFVMCAQFHPT

-EDLVVSASLDETVRIWDI--------S---------GL-RKRH-S-A-P------G---

-----SQ---------SF-E--------E-QM-------------------I--------

-TQQNLFD--------------GGFGD--CVVKFILEGHTRGVNWASFHPTLPLIVSGSD

DRQVKLWRM------------S----------------STKAWEVDTCRGHTNNVDSVIF

HPFQ-NLIISVGEDSTIRVWDLDKRTPVKQFKR-EQ----------DRFWSIRAHPNVNL

FGAAHDSGIMVFKLD

>UniRef90_L1JYZ3_8_333 | Coatomer subunit alpha n=1 Tax=Guillardia theta (strain CCMP2712) TaxID=905079 RepID=L1JYZ3_GUITC | E_val=1.9e-149

--FLTKFETKSNRVKGLSFH--PRRPWIVASLHNGVIQLWDYRMGTLLD-------RFEE

HEG-PVRGVNFHQT-QPLLVSGGDDYKIKIWNYK--LR-------------RCLKPDLLG

HLDYIRTVQFHHE---YPWVVSSSDDQTVRIWDWQQSGRPCLVVLTGHNHYVMSVQFHPR

-EDLIVSASLDQTVRVWDI--------S---------GL-KKKY-V-G-H------G---

-----VD---------DM-Q--------G-QTS---------------VP-K--------

-IGADLF----------------GSTD--TTVKYLLEGHDRGVNWASFHHTLPLIVSGAD

DRQVKLWRM------------N----------------DSKAWEVDTLRGHVNNVSCVLF

HPKQ-ELIISNSEDKSIRVWDMSKRTGVQTFRR-EQ----------DRFWILAVHKDQGL

LAAGHDSGMLVFKLD

>UniRef90_A7TGU6_1_328 | Coatomer subunit alpha n=1 Tax=Vanderwaltozyma polyspora (strain ATCC 22028 / DSM 70294 / BCRC 21397 | E_val=3.4e-149

MKLLTKFESKSTRAKGIAFH--PSRPWVLVALFSSTIQLWDYRMGTLLH-------RYED

HEG-PVRGIDFHPT-QPLFVSAGDDYTIKVWSLE--TN-------------KCL-YTLDG

HLDYVRTVFFHKE---LPWIISASDDQTIRIWNWQ--NRKEIACITGHNHFVMCAQFHPT

-EDLIVSASLDETVRVWDI--------S---------KL-REKH-S-A-P------G---

-----RSA-----MPTSF-E--------E-KI-------------------A--------

-AQQNLLD--------------GGFGD--CTVKFILEGHTRGVNWASFHPTLPLIVSGGD

DRQVKLWKM------------S----------------ATKAWELDSCRGHTNNVDSVIF

HPTQ-NLILSVGEDKTLRVWDLDKRTPVKQFKR-EN----------DRFWLIAAHPNINL

FGAAHDSGIMIFKLD

>UniRef90_D8QH78_3_336 | Coatomer subunit alpha n=2 Tax=Schizophyllaceae TaxID=5332 RepID=D8QH78_SCHCM | E_val=1.1e-148

V-MLTKFESKSNRVKGLAFH--PTQPLLAAALHNGSVQLWNYRMGVLVD-------RFEE

HEG-PVRGVAFHPS-RPLLVTGGDDYKVRVWDIRPQNR-------------RCL-FTLHG

HLDYVRTVQFHHE---MPWIISTGDDQTIRIWNST--SRNCIAILTGHSHYIMSAFFHPK

-DDLVVSASMDQTVRVWDI--------S---------GL-RKGA-P-N-STPGGGMG---

-----GPGGPG----GGG-G--------G------------------ASG-A--------

-GGFEAF-----------------DSF--STVKYVLEGHDRGVNFASFHPTLPLIVSAAD

DRVIKIWRM------------S----------------ETKAWEVDSCRGHFNNVSCAIF

HPKH-ELILSCGEDKTIRVWDLAKRTAIQTFRR-EH----------DRFWVLAAHPNLNL

FAAGHDSGLIVFKLE

>UniRef90_D7LBD0_1_319 | Coatomer subunit alpha n=2 Tax=Arabidopsis lyrata subsp. lyrata TaxID=81972 RepID=D7LBD0_ARALL | E_val=5.7e-148

--MLTKFKTKSNRVKGLSFH--PKRPWILASLHSGVIQLWDYRVGTLID-------KFDG

HQG-PVRGVHFHTS-QPLFVSGGDDCKIKVWNYK--TH-------------WCL-FTLLG

HLDYIRTVQFHHE---YPWILSASDDQTIRIWNWQ--SRTCVSVLAAHNHYVMCASFHPK

-DDLVVSASLDQTVRVWEI--------G---------AL-KKKT-V-S-P------S---

-------------------D--------D-IMR---------------LA-E--------

-INSDLF----------------DSVD--VTVKYVLEGHERGVIWAAFHPNLPLIVSGSD

DRQVKLWRM------------N----------------ETKAWEVDTLRGHMNNVSSVMF

HAKQ-DIIVSNSEDNSIRVWDATKRTEIQTFRR-EH----------DRFWSLAVHPEINL

LAAGHDNGMIVFKLE

>UniRef90_UPI00071174B0_1_318 | coatomer subunit alpha-1-like protein n=1 Tax=Blastocystis sp. subtype 4 TaxID=944170 RepID=U | E_val=1.3e-147

--MLAKFETKSSRVKGISFH--PTRPWLLCSLHDGQIQLWDYRVGTLLE-------TFDE

HDG-PVRSVDFHST-QPLFVSGGDDYKIRVWNYN--NK-------------RSL-FTLMG

HLDYIRTVQFHPE---NPWIVSCSDDQNIRIWNWQ--SRECIAVLTGHNHYVMSAQFHPK

-EDLIVSASLDQTIRVWDI--------S---------GL-KQKG-S-K-M------S---

-----SGP-------KTA-R----------SV----------------IG-H--------

-LGTDIV----------------G------TVKYVLEGHERGVNWASFHPELPLIVSGSD

DRMIKIWRT------------N----------------ETKAWEVDTLRGHTNNVSCVMF

HPRE-DLILSNSEDHSIRVWDSTKRIGIQSFVR-AH----------DRFWIITVHKKQNL

LAAGHDSGTVVFKL-

>UniRef90_A0A1E7F6J1_1_333 | Coatomer subunit alpha n=3 Tax=Bacillariaceae TaxID=33852 RepID=A0A1E7F6J1_9STRA | E_val=2.1e-147

--MLTKFESKSARVKGLAFH--PYRPWVCASLHNGIIQLWDYRVGTVID-------RFEE

HDG-PVRGVDFHKT-EPYIVSGGDDYKIKVWDYK--LR-------------RCL-FTLLG

HLDYIRTVQFHPDPAKFPWILSASDDQTLRLWDFH--KRTCLSVLTGHNHYVMCAAFHPS

-EDLIVSASLDQTVRVWDT--------T---------GL-RKKQ-T-G-E------G---

-----AM---------------------D-PMRG---PPV--SATTNAAM-N--------

-VQAELF----------------GTND--VVVKYVLEGHDRGVNWACFHPTLPLLASAAD

DRQVKLWRM------------S----------------ETKAWEVDTLRGHANNVSSCLF

HPKH-DLVVSNSEDRSIRVWDVSKRVGVQTFRR-EG----------DRFWILAAHANQNL

LAAGHDSGMIVFKLE

>UniRef90_A0A067MD04_1_315 | Coatomer subunit alpha n=1 Tax=Botryobasidium botryosum FD-172 SS1 TaxID=930990 RepID=A0A067MD04 | E_val=3.8e-147

MQMLTKFESKSNRVKGLAFH--PTRPLLAASLHNGSVQLWNYQMGTLVD-------RFDE

HDG-PVRGVAIHPS-RPLLVTGGDDYKIKVWDIRPQNR-------------RCL-FTLHG

HLDYVRTVQFHHE---MPWILSASDDQTIRIWNST--SRNCIAILTGHSHYIMSAQFHPK

-EDLIVSASMDQTVRVWDI--------S---------GL-RKST-P-N-T----------

-------------------------------------------------Q-P--------

-GSFDNF-----------------DTF--STVKYVLEGHDRGVNFASFHPTLPLIISAAD

DRQIKLWRM------------S----------------DTKAWEVDTLRGHFNNVSSALF

HTRH-ELLLSVGEDKTIRVWDLTRRTAVQTFRR-EH----------DRFWVLTAHPELNL

FAAGHDNGLIVFKLE

>UniRef90_D8M8H5_1_318 | Coatomer subunit alpha n=1 Tax=Blastocystis hominis TaxID=12968 RepID=D8M8H5_BLAHO | E_val=9.4e-147

--MLAKFETKSSRVKGIAFH--PTRPWILCSLHDGCIQLWDYRVGTLLE-------TFSE

HDG-PVRSVDFHPS-QPLFVSGGDDYKIRVWNYN--NK-------------RSL-FTLMG

HLDYIRTVQFHHE---NPWIVSCSDDQNIRIWNWQ--SRECIAVLTGHNHYVMCAQFHPK

-EDLVVSASLDQTIRVWDI--------S---------GL-KQKG-K-K-I------P---

-----GKT-------GGP-S----------TM----------------LG-R--------

-LSTDLV----------------G------TVKYVLEGHERGVNWASFHPELPLIVSGSD

DRMIKIWRT------------N----------------ETKAWEVDTLRGHTNNVSCVMF

HPRE-DLILSDGEDHSIRVWDSTKRIGIQSFVR-AH----------DRFWIIIAHKTQNL

LAAGHDSGAVVFKL-

>UniRef90_A0A167X8G2_1_286 | Coatomer subunit alpha n=1 Tax=Ascosphaera apis ARSEF 7405 TaxID=392613 RepID=A0A167X8G2_9EURO | E_val=2.5e-146

-------------------------------------------MGTLID-------RFEE

HDG-PVRGIDFHPT-QPLFVSGGDDYKIKVWSLQ--TR-------------RCL-FTLNG

HLDYVRTVFFHHE---LPWILSASDDQTIRIWNWQ--NRSLICTMTGHNHYVMCAQFHPK

-EDLIVSASLDQTVRVWDI--------S---------GL-RKKH-S-A-P------TS--

-----SL---------TF-E--------D-QMNR--------------AN-A--------

-AQADMF----------------GNTD--AIVKFVLEGHDRGVNWVSFHPTLPLIVSAGD

DRLIKLWRM------------G----------------DTRAWEVDTCRGHFQNSGCALF

HPHQ-DLILSVGEDKTIRVWDLNKRTSVQSFKR-DL----------DRFWVIAAHPEINL

FAAGHDTGVMVFKLE

>UniRef90_UPI0004417AF9_3_316 | coatomer subunit alpha-2 n=1 Tax=Punctularia strigosozonata (strain HHB-11173) TaxID=741275 R | E_val=8.9e-146

V-MLTKFESKSNRVKGLCFH--PTQPLLAAALHNGSVQLWNYRMGVLVD-------RFEE

HEG-PVRAIAIHPS-RPLLATGGDDYKIKVWDLRPQSR-------------RCL-FTLHG

HLDYIRTVHFHHE---MPWIISCSDDQTIRIWNST--SRNCIAILTGHSHYVMSAFFHPK

-EDLVVSASMDQTVRVWDI--------S---------GL-RKGT-P-N-T----------

-------------------------------------------------Q-P--------

-GAFDTF-----------------DNF--STVKYVLEGHDRGVNWASFHPTLPLIVSASD

DRQVKIWRM------------S----------------ETKAWEVDACRGHFNNVLCALF

HPMR-ELIVSCGEDKTIRVWDLQKRAAIQTFRR-EQ----------DRFWGLAAHPHLNL

FAAAHDSGLIVFKLE

>UniRef90_U4LH34_1_285 | Coatomer subunit alpha n=1 Tax=Pyronema omphalodes (strain CBS 100304) TaxID=1076935 RepID=U4LH34_PY | E_val=2e-145

-------------------------------------------MGTLID-------RFEE

HDG-PVRGVDFHRT-QPLFVSGGDDYKIKVWSLT--TR-------------RCL-FTLNG

HLDYVRTVFFHHE---LPWIVSSSDDQTIRIWNWQ--NRQLICNMTGHNHYTMCAQFHPK

-EDLIVSASLDQSVRVWDI--------S---------GL-RKKH-S-A-P------T---

-----TM---------TF-E--------D-QMSR--------------AN-G--------

-NQPDMF----------------GNTD--AVVKFVLEGHDRGVNWVSFHPTLPLIISAGD

DRNVKLWRM------------S----------------ETKAWEVDTCRGHFQNASACLF

HPNQ-DLILSVGEDKTIRVWDLNKRTSVQSFKR-EN----------DRFWVIAAHPEINL

FAAGHDNGVMVFKLE

>UniRef90_A0A166DSM6_3_316 | Coatomer subunit alpha n=2 Tax=unclassified Peniophora TaxID=2635284 RepID=A0A166DSM6_9AGAM | E_val=2.4e-145

V-MLTKFESKSNRVKGLAFH--PTQPLLAASLHNGSVQLWNYRMGVLVD-------RFEE

HEG-PVRAVAIHPS-RPLLASGGDDYKVKVWDLRPQSR-------------KCL-FTLHG

HLDYIRTVQFHHE---MPWILSCSDDQTIRIWNST--SRQCIAILTGHSHYVMSASFHPK

-DDLVVSASMDQTVRVWDI--------S---------GL-RKST-P-N-T----------

-------------------------------------------------A-P--------

-GTFDTY-----------------DSF--STVKYVLEGHDRGVNFAQFHPTLPLIVSAGD

DRQVKIWRM------------S----------------ETKAWEVDACRGHFNNVLSAVF

HPKH-ELIVSCGEDKTVRVWDLAKRTAVQTFRR-EH----------DRFWTLAAHPELNL

FAAGHDSGLIVFKLE

>UniRef90_A0A409XWB8_3_325 | Coatomer subunit alpha n=2 Tax=Agaricales TaxID=5338 RepID=A0A409XWB8_PSICY | E_val=5.1e-145

V-MLTKFESKSNRVKGLAFH--PTQPLLAASLHNGCVQLWNYRMGVLVD-------RFEE

HEG-PVRAVAIHPS-RALLCTGGDDYKIKVWDIRPQNR-------------RCL-FTLHG

HLDYVRTVQFHHE---MPWIISASDDQTIRIWNST--SRQCIAVLTGHSHYVMSVQFHPK

-EDLIVSASMDQTVRVWDI--------S---------GL-RKGS-P-N-Q------G---

-----GP---------GS-S--------N------------------SNG-P--------

-GNFETF-----------------DTF--STVKHVLEGHDRGVNFATFHPTLPLILSAGD

DRVIKIWRM------------S----------------ETKAWEVDSCRGHFNNVSTALF

HPKH-ELIVSCGEDKTVRVWDLGKRTAIQTFRR-EQ----------DRFWVLAAHPNLNL

FAAGHDSGLIVFKLE

>UniRef90_A0A0C9T7Z0_3_318 | Coatomer subunit alpha n=1 Tax=Plicaturopsis crispa FD-325 SS-3 TaxID=944288 RepID=A0A0C9T7Z0_PL | E_val=7.4e-145

V-MLTKFESKSNRVKGLAFH--PTQPLLAASLYNGSVQLWNYRMGVLVD-------RFEE

HEG-PVRAVAIHPS-RALLATGGDDFKIKVWDIRPQNR-------------RCL-FTLHG

HLDYVRTVQFHHE---MPWILSCADDQTIRIWNST--SRNCIAILTGHSHYVMSAQFHPK

-DDLIVSASMDQTVRVWDI--------S---------GL-RKAGGPNN-G----------

-------------------------------------------------G-P--------

-GNFETF-----------------DTF--STVKYVLEGHDRGVNYAMFHPTLPLIVSAGD

DRVIKIWRM------------S----------------ETKAWEVDSCRGHFNNVLCALF

HPKH-ELIVSCGEDKTVRVWDLAKRTAIQTFRR-EQ----------DRFWALAAHPNLNL

FAAAHDNGLIVFKLE

>UniRef90_A0A2W1DSH4_8_281 | Coatomer subunit alpha n=2 Tax=Pyrenophora tritici-repentis TaxID=45151 RepID=A0A2W1DSH4_9PLEO | E_val=4.2e-144

---VSEFESKSSRAKGIAFH--PKRPWILVSLHSSTIQLWDYRMGTLID-------RFEE

HDG-PVRAVDFHKT-QPLFVSGGDDYKIKVWSYQ--TR-------------RCL-FTLNG

HLDYVRTAYFHHE---LPWILSCSDDQTIRIWNWQ--NRSLICTMTGHNHYVMAASFHPK

-EDLVVSASLDQSVRVWDI--------S---------GL-RKKH-S-A-P------Q---

-----AM---------SF-E--------D-QMAR--------------AN-Q--------

-NQADMF----------------GNTD--AVVKFVLEGHDRGVNFVAFHPTLPLIVSAGD

DRLVKLWRM------------S----------------ETKAWEVDTCRGHFQNASACLF

HPHQ-DLILSVGEDS---------------------------------------------

---------------

>UniRef90_A0A409YCM7_3_325 | Coatomer subunit alpha n=1 Tax=Panaeolus cyanescens TaxID=181874 RepID=A0A409YCM7_9AGAR | E_val=7.3e-144

V-MLTKFESKSNRVKGLAFH--PTQPLLAASLHNGSVQLWNYRMGVLVD-------RFEE

HEG-PVRAVAIHPT-RALLATGGDDYKIKVWDIRPQNR-------------RCL-FTLHG

HLDYVRTVQFHHE---MPWILSASDDQTIRIWNST--SRQCIAVLTGHSHYVMSAQFHPK

-EDLIVSASMDQTVRVWDI--------S---------GL-RKGS-P-N-T------G---

-----GP---------GS-S--------N------------------QSG-P--------

-GSFETF-----------------DSF--STVKHVLEGHDRGVNYAMFHPTLPLIISAAD

DRAIKIWRM------------S----------------ETKAWEVDSCRGHFNNVSSALF

HPKH-ELIVSCGEDKTVRVWDLAKRSSIQTFRR-EH----------DRFWVLASHPNLNL

FAAGHDNGLIVFKLE

>UniRef90_A0A0D0BBD5_3_316 | Coatomer subunit alpha n=65 Tax=Boletales TaxID=68889 RepID=A0A0D0BBD5_9AGAM | E_val=1.3e-143

V-MLTKFESKSNRVKGLAFH--PTQPLLAAALHNGSVQLWNYRMGVLVD-------RFEE

HEG-PVRGVAIHPS-RALLVTGGDDYKIKVWDLKPQSR-------------RCL-FTLHG

HLDYVRTVQFHHE---MPWILSASDDQTIRIWNST--SRHCVAILTGHSHYVMSAQFHPK

-DDLIVSTSMDQTVRVWDI--------S---------GL-RKNT-P-N-N----------

-------------------------------------------------G-P--------

-NNFETF-----------------DTF--STVKYVLEGHDRGVNWASFHPTLPLIVSAAD

DRTIKIWRM------------S----------------ETRAWEVDSCRGHFNSVSSAIF

HPKH-ELIVSCAEDKTVRVWDLAKRTAIQTFRR-EN----------DRFWILAAHPNLNL

FAAGHDGGLIVFKLE

>UniRef90_A0A2C6KKY8_4_331 | Coatomer subunit alpha (Fragment) n=1 Tax=Cystoisospora suis TaxID=483139 RepID=A0A2C6KKY8_9APIC | E_val=2.2e-143

--MLVKCETKSSRVKGLAFH--PSLQWILAALHNGTIQLWDYRIGSLID-------KFEE

HEG-PVRGIDFHSS-QQLFVSGGDDYKVKLWSLS--TR-------------KCI-FTFVG

HLDYLRTAFFHHI---YPWILSASDDQTVRIWNWQ--SRSCIAVLTGHNHYVMSALFHPY

-EDLVVSASLDQTIRVWDT--------S---------GL-REKT-G-G-A------A---

-----GS---------SFGR--------GGSL-----------GGSGNNR-R--------

-PHADMF----------------TAND--AVCKFVLEGHERGVNWAAFHPSMPLIASAAD

DRTIKLWRY------------N----------------DSKAWEVDTLRGHFNNVSCLVF

HPQR-ELLISNSEDRTIRVWDVSKRVGVHTFRR-ES----------DRFWIIAAHRSSSA

LAVGHDSGMVVFKL-

>UniRef90_A0A7R8CIR8_1_289 | COPA n=1 Tax=Lepeophtheirus salmonis TaxID=72036 RepID=A0A7R8CIR8_LEPSM | E_val=1.8e-142

--MLTKFETKSPRVKGLAFH--PKRPWILASLHNGVIQLWDYRMCTLLE-------KFDE

HEG-PVRGIDFHKQ-QPLFVSGGDDYKIKVWNYK--LK-------------RCL-FQLLG

HLDYIRTTFFHHE---YPWVLSASDDQTIRIWNWQ--SRSCVSVLTGHNHYVMSASFHPS

-EDLLVSASLDQSVRVWDI--------S---------GL-RKKN-V-A-L------G---

-----PD---------DV-H--------H-SL----------------KS-T-----GSA

ANNTDLF----------------GVAD--CVVKHVLEGHDRGVNWASFHPTMPLIVSGAD

DRQIKLWRM------------N----------------ESKAWEVDTCRGHYNNVSCVLF

HPRQ-DLIISNSEDRTIRVWDTAKRTS---------------------------------

---------------

>UniRef90_A0A2A9MED7_4_334 | Putative coatomer protein complex, subunit alpha n=1 Tax=Besnoitia besnoiti TaxID=94643 RepID=A0 | E_val=6.4e-142

--MLVKCETKSSRVKGLAFH--PSLQWILAALHNGTIQLWDYRIGSLID-------KFEE

HEG-PVRGIDFHSS-QPLFVSGGDDYKVKLWSLT--TR-------------KCI-FTFLG

HLDYLRTVFFHHI---YPWILSASDDQTVRIWNWQ--SRQCIAVLTGHNHYVMSALFHPY

-EDLVVSASLDQTIRVWDT--------S---------GL-REKT-G-G-A------G---

-----GAH--------AFGR--------SGSTAA---------GVPGARR-H--------

-HPSEMF----------------TAND--AVCKFVLEGHERGVNWAAFHPSMPLIASAAD

DRTIKLWRY------------N----------------DAKAWEVDTLRGHFNNVSCLVF

HPQR-ELLISNSEDRTIRVWDVSKRVGVHTFRR-ES----------DRFWIIAAHRTCSA

LAVGHDSGMVVFKL-

>UniRef90_B6AAR9_1_323 | Coatomer alpha subunit protein, putative n=2 Tax=Cryptosporidium TaxID=5806 RepID=B6AAR9_CRYMR | E_val=2.2e-141

--MLIRCESKSTRVKGLSFH--PKLPWILVSLHNGIIQFWDYRLGSLLD-------TYEE

HEG-PVRSVDFHES-QPIFVSGGDDYRVKVWNYK--ER-------------RCL-FTLIG

HLDYIRTVEFHKE---YPWILSSSDDQTMRLWNWQ--SRACIAVITGHNHYVMCSKFHPH

-QDLIVSASMDQSIRIWDF--------T---------GL-REKT-V-K-G------H---

-----SSL-----------S--------T-SISN--------------TM-P--------

-AHSDMF----------------GAND--VICKFVLEGHERGVNWVTFHPTLSLIASASD

DRTIKLWRY------------S----------------ETKAWEIDTLRGHFNNVSSVIF

HTSK-DWLLSDSEDRTIRIWDLTKRIPLHTYKR-EG----------DRFWAIVSHPTSSL

FAAGHDSGMMIFKLE

>UniRef90_D3B3V5_1_315 | Coatomer subunit alpha n=1 Tax=Polysphondylium pallidum (strain ATCC 26659 / Pp 5 / PN500) TaxID=670 | E_val=6.3e-141

--MLYKYETKSNRVKGLSFH--PTRPWILASLHSGAIHLYDYRIKTLLE-------KFEE

HDG-PVRGVNFHMT-QPLFVSGGDDYKIKVWNYK--QR-------------RCL-FTLKG

HKDYIRTVEFHRE---APWILSASDDQVIRIWNWQ--SRTCIAELNGHNHYVMCASFHPK

-DDLIVSASLDQTIRIWDI--------S---------GL-KKKT-T-T-I------K---

-----PY---------PQ-N--------D-TM-------------------R--------

-LQEEIF----------------G-TD--VVVKLSLEGHDRGVNWAAFHPTQPYIVSASD

DHQVKLWKM------------N----------------DN----VDSFRGHFNNVSCALF

HPRQ-DLIISDSEDKTIRVWDMAKKTTIQTIRR-EH----------DRFWTLASHPNANL

FAAGHDSGMIVFKLE

>UniRef90_A0A6A7FQ53_1_319 | Coatomer subunit alpha n=1 Tax=Hirondellea gigas TaxID=1518452 RepID=A0A6A7FQ53_9CRUS | E_val=4.2e-140

--MLIKFETKSNRVKGLAFH--HKRPWILASLHNGVIQLYDYRMEVLVE-------KFEE

HDG-PVRGVDFHFS-QPLFVSGGDDYKIKVWNYN--LH-------------RCL-FQLVG

HTDYIRSVKFHHE---APWILSASDDQTLRIWNWQ--NRTCLSVLTGHNHYVMCGEFHPT

-EDLVLSASLDESVRVWDI--------S---------GL-RKRS-S-Q-N------L---

-----FH------------Q--------D-------------------FR-P--------

-GAQELF---------------GGSHD--VTVKHILEGHSRGVNWACFHKTMPLIVSGAD

DREVKLWRM------------S----------------ESKAFIVDTMRGHTNNVSCVIF

HPKR-ELIVSNSEDKSVRVWDISKQNRTQTFRR-VN----------DRYWILAAHPKLNL

LAAGHDNGFIVFKLE

>UniRef90_A0A1Y1HU83_1_280 | Coatomer subunit alpha n=1 Tax=Klebsormidium nitens TaxID=105231 RepID=A0A1Y1HU83_KLENI | E_val=1.5e-138

-------------------------------------------MGTLID-------RFDE

HDG-PVRGVHFHKS-QPLFVSGGDDYKIKVWNYK--MR-------------RCL-FTLLG

HLDYIRTVQFHQE---YPWIVSASDDQTIRIWNWQ--SRNCISVLTGHNHYVMCAAFHIK

-EDLVVSASLDQTVRVWDI--------S---------AL-RKKT-V-S-P------A---

-------------------E--------D-ILR---------------LP-Q--------

-MNTDLF----------------GGGD--AIVKYVLEGHDRGVNWAAFHPTLPLIVSGAD

DRQVKLWRM------------N----------------DTKAWEVDTLRGHVNNVSCVMF

HARQ-DIIVSNSEDKSIRVWDMSKRTGVQTFRR-EH----------DRFWILAAHPEMNL

LAAGHDSGMIVFKLE

>UniRef90_A0A6A4XBX0_1_331 | Coatomer subunit alpha n=3 Tax=Thoracica TaxID=6676 RepID=A0A6A4XBX0_AMPAM | E_val=5.1e-138

--MLTKFETKSARVKGLSFH--PKRPWVLASLHSGVIQLWDYRMCTLLE-------KFDE

HDG-PVRGICFHMQ-QPLFVSGGDDYKIKYVTDT--SRRRCGTTSRSGACSRCS-ATWTT

SGPQSSTTSTRGS---SP----ASDDQTIRIWNWQ--RRSCICILTGHNHYVMCARFHPT

-EDLVVSASLDQTVRVWDI--------S---------GL-RQKH-V-A-P------G---

-----PS---------GL-D--------D-HL----------------KS-T--------

-ASPDLF----------------GQAD--AVVKHVLEGHDRGVNWAAFHPSLPLIVSGAD

DRQIKFWRM------------N----------------DSKAWEVDTCRGHYNNVSCVLF

HPRQ-ELILSNSEDKSIRVWDMSKRTCLHTFRR-EH----------DRFWVMAAHPQMNV

FAAGHDSGMIIFKLE

>UniRef90_F0ZEM4_1_316 | Coatomer subunit alpha n=1 Tax=Dictyostelium purpureum TaxID=5786 RepID=F0ZEM4_DICPU | E_val=3.1e-136

--MLYKFETKTNRVKGLSFH--PTRPWILTSLHSGSIHLYDYRIKTLLE-------KFEE

HEG-PVRGVNFHMT-QPLFVSGGDDYKIKVWNYK--QR-------------RCL-FTLKG

HKDYVRTVEFHRE---APWIVSSSDDMVIRIWNWQ--SRTCISELSGHNHYVMSALFHPK

-DDLVVSASLDQFIRVWDI--------S---------GL-KKKM-T-T-V------K---

-----PY---------RE-N--------D-PM-------------------R--------

-IQEEIF----------------G-TD--VIVKLSLEGHDRGVNWAAFHPTQPYIVSASD

DHHVKLWRM------------N----------------DPI---VDTFRGHYNNVSCALF

HPRQ-ELIISNSEDKTIRVWDIVKKQTVHMIRR-DN----------DRFWTLASHPNQNL

FAAGHDSGMIVFKLE

>UniRef90_A0A6L5CXI1_1_285 | WD repeat-containing protein 55 homolog n=1 Tax=Ephemera danica TaxID=1049336 RepID=A0A6L5CXI1_9 | E_val=3.9e-135

-------------------------------------------MCTLLD-------KFDD

HDG-PVRGICFHSQ-QPLFVSGGDDYKI-----K--QR-------------RCI-FTLLG

HLDYIRTTMFHHE---YPWILSASDDQTIRIWNWQ--SRTCICVLTGHNHYVMCAQFHPT

-DDIVVSASLDQTVRVWDI--------SGIASQ--RQGL-RKKN-V-A-P------G---

-----PG---------GL-E--------D-HM----------------RN-P--------

-GSTDLF----------------GQAD--AVVKHVLEGHDRGVNWACFHPTLPLIVSGAD

DRQIKLWRM------------N----------------EAKAWEVDTCRGHYNNVSCVLF

HPRQ-ELILSNSEDKSIRVWDMTKRTCLHTFRR-EN----------ERFWVLAAHPTLNL

FAAGHDNGMIVFKLE

>UniRef90_A0A1Q3DX44_3_307 | Coatomer subunit alpha n=2 Tax=Omphalotaceae TaxID=72117 RepID=A0A1Q3DX44_LENED | E_val=3.4e-134

V-MLTKFESKSNRVKGLAFH--PTQPLLAAGLHNGSIQLWNYRMGVLVD-------RFEE

HEG----------T-RALLVSGGDDYKIKVWDIRPQNR-------------RCL-FTLHG

HLDYLRTTQFHHE---MPWIISCSDDQTIRIWNST--SRNCIAVLTGHSHYVMSAFFHPK

-DDLVVSASMDQTVRVWDI--------S---------GL-RKGS-P-N-S----------

-------------------------------------------------G-P--------

-GNFETF-----------------DTF--STVKYVLEGHDRGVNYATFHPTLPLILSAAD

DRTIKIWRM------------S----------------ETKAWEVDSCRGHFNNVSAAIF

HPKH-ELIVSCGEDKTVRVWDLAKRTAIQTFRR-EH----------DRFWDLAAHPNLNL

FAAGHDSGLIVFKLE

>UniRef90_A0A5A8DF99_1_346 | WD_REPEATS_REGION domain-containing protein n=6 Tax=Cafeteria roenbergensis TaxID=33653 RepID=A0 | E_val=7.8e-133

--MLTKFDTKSSRVKGLAFH--PKRPWILASLHDGIVQLWDYRNGTCIE-------RFAG

HEG-PVRGLDVHPSGQPLFVSGGDDYLIKVWSLA--QR-------------KCL-FTLIG

HLDYIRTVQFHHE---YPWIVSASDDQTIRIWNWQ--SRACIATMPGHSHYVMCARFHPR

-DDLVVSASLDQTVRVWDL--------T---------PL-RKRS-V-R-G------A---

-----GGIPPA---SAGA-T--------T-TMAAL----------------R--------

-AQAQRFASAGAGGPGARPAGVQDSGE--PFVRDIFDSHDRGVNWATFHPTLPLIISAAD

DRTMKVWRM------------S----------------DGRAWEVDTLRGHNNNVSCVLF

HPHC-DLIVSNSEDRSIRIWDGTKRMALSTYRR-ET----------ERFWVLAAHPNENL

LAAGHDRGLVVFKLE

>UniRef90_A0A0S4IPS1_4_314 | Coatomer subunit alpha n=1 Tax=Bodo saltans TaxID=75058 RepID=A0A0S4IPS1_BODSA | E_val=2e-131

--LLVKFETRSFRVKGISFH--PKRPWVMIGMHNGVVQIVDYRMCATID-------RYEE

HDG-PVRGVDFHST-QPLFVTGGDDYKIKVWNYK--FR-------------RCL-FTLTG

HLDYIRTVQFHKE---QPWILSASDDQTLRIWNWQ--SRNCMSVLTGHNHYVMSAQFHPR

-EDLIVSASLDLTIRVWDI--------S---------GL-KARR-N-D-N------N---

------------------------------------------------SG-A--------

-ISQDLF----------------GSSD--VMVKFNLEGHEKGVNWASFHPTKPLIVSAAD

DRSIRIWKM------------D----------------DTRAYEINQLRQHTNNVSSVMY

F--K-DYIISNSEDRTIRVWDPAQNTAINTYRR-DT----------DRFWTLAAHPESNL

IAAGHDNGMMVFKL-

>UniRef90_A0A0V0J4C6_1_286 | WD_REPEATS_REGION domain-containing protein (Fragment) n=4 Tax=Diphyllobothriidae TaxID=28843 Re | E_val=5.8e-129

------------------------------------------------------------

--G-PVRGIDFHKN-QPLFVSGGDDYKIKVWNYK--HR-------------KCL-FTLLG

HLDYIRTTYFHHE---HPWILSASDDQTIRIWNWQ--SRTPACVLTGHSHYVMCAQFHPK

-EDLVVSASLDQTVRIWDI--------S---------GF-RKKN-V-A-P------T---

-----SMT--------GL-E--------E-QMRYLTGGSG--HGGSSSPI-G--------

-GNAELF----------------GTAD--AVVKHVLEGHDRGVNWVAFHPTLPIVVSASD

DRHIRLWRV------------T----------------ETKAWTLDTLRGHVNNVSCVLF

HPKQ-DLLLSNSEDKSIRVWDLAKRTCVSTIRR-DN----------DRFWVLAAHPTLSL

FAAGHDEGMVVFKLE

>UniRef90_A0A1R2BB29_1_311 | Coatomer subunit alpha n=1 Tax=Stentor coeruleus TaxID=5963 RepID=A0A1R2BB29_9CILI | E_val=3.2e-128

--MLVKFETSSNRVKGISFH--PTRYWILSSLHNGAIQLWDYNAKTLID-------SYEE

HEG-PVRGICFHNT-QPLFASGGDDNKVKIWNFK--QK-------------KCL-FTLEG

HLDYIRTVSFHHE---LAWLMSSSDDQTIRIWNWQ--SRACIAILTGHNHYIMSASFHAS

-EDMIISASLDQTVRVWDF--------K---------SL-RQKY-Y-A-S------G---

------------------------------------------------------------

-KSADVM----------------LGND--VVVKFVLEGHDRGVNWASFQNSSQYIVTAAD

DRSVRLWRY------------T----------------DTKAWEVEVMRGHSNNVSSAIF

HPNL-DVIVSNSEDKSLRIWDVTRRTCIYTHKK-DD----------DRFWVLASHPSLNI

IAAGTDSGMVIFKLE

>UniRef90_A0A409YWR3_1_283 | Coatomer subunit alpha n=1 Tax=Gymnopilus dilepis TaxID=231916 RepID=A0A409YWR3_9AGAR | E_val=3.1e-126

-------------------------------------------MGVLVD-------RFEE

HEG-PVRGIAIHPT-RALLVSGGDDYKIKVWDIRPQNR-------------RCL-FTLHG

HLDYVRTVQFHHE---MPWILSCSDDQTIRIWNST--SRQCIAVLTGHSHYVMSAQFHPK

-EDLIVSASMDQTVRVWDI--------S---------GL-RKGS-P-N-Q------G---

-----GP---------GS-S--------G------------------PNG-S--------

-SNFETF-----------------DTF--STVKHVLEGHDRGVNWATFHPTLPLIISAGD

DRVIKIWRM------------S----------------ETRAWEVDSCRGHFHNVLTALF

HPKH-ELIVSCGEDKTIRVWDLAKRSAIQTFRR-EQ----------DRFWILACHPNLNL

FAAGHDSGLIVFKLE

>UniRef90_A0A023BCE3_1_319 | Putative coatomer subunit alpha n=1 Tax=Gregarina niphandrodes TaxID=110365 RepID=A0A023BCE3_GRE | E_val=2e-123

--MLVKCESKSARVKAVAFH--PKLPWVLVGLHSGPIHLWDYKRGCILH-------KFED

HVG-PVRALHFHPN-QPLFCSAGDDHDIRVWNYS--TH-------------QKL-FSLSG

HLDYVRAVEFHYE---YPWIVSCSDDQTVRIWNWQ--SRQCLSVLAGHHHFVMYVRFHPS

-LDLIASASMDQTIRVWDV--------S---------GL-KEKT-V-A-R------L---

-----HSL-----------N-------------------------PGTNA-R--------

-NSTEMF----------------QPQD--AVCKYIIEGHDRGVNWIDFHPHVNMLVSGSD

DRMVKLWRF------------D----------------ETRWWEVDTFRGHFNNVSAVKF

HPIL-DVIISNSEDHTIRIWDVNQRKLLNSVRR-ES----------DRFWVLDCQ--QNS

IAVGHDSGMIVFKLQ

>UniRef90_K2N6X5_21_329 | Coatomer subunit alpha n=6 Tax=Trypanosoma cruzi TaxID=5693 RepID=K2N6X5_TRYCR | E_val=1.2e-120

-DMLTKFDVRSCRVKGISFH--KSRPWVLCGLHNGTVQIWDYRTNTSID-------TYTE

HSG-SVRGVDFHIS-QPLFVSGADDYLIKVWNYK--LR-------------RCL-FTLRG

HMDYIRVTFFHHE---QPWILSCSDDFTVRIWNWQ--SRSSVACLPGHNHYVMCAQFHPR

-EDLVVSASLDRTIRVWDI--------S---------SL-RLRK-Q-E-V------G---

------------------------------------------------------------

-IAQDLL----------------GTTD--VTLKYLLEGHEKGVNWVCFHPTKPYIASAAD

DRTVRVWRM------------M----------------ESSCHEELQLRGHTNNVCCVTY

--LK-DFLISDSEDRTIRVWDVKSRSPIMVFRR-DT----------DRYWILATLPEKNL

IAAGHDSGMQVFKL-

>UniRef90_A0A553NY15_1_241 | Coatomer subunit alpha n=1 Tax=Tigriopus californicus TaxID=6832 RepID=A0A553NY15_TIGCA | E_val=4.7e-118

--MLTKFETKSPRVKGLAFH--PKRPWILASLHNGVIQLWDYRMCTLLE-------KFDE

HEG-PVRGIGFHAQ-QPLFVSGGDDYKIKVWNYK--LK-------------RCL-FHLLG

HLDYIRTTVFHHE---YPWILSASDDQTIRIWNWQ--SRACISVLTGHNHYVMCALFHPS

-DDLVCSASLDQTVRVWDI--------S---------GL-RKKN-V-A-P------G---

-----RG---------GL-D--------D-HM----------------RN-P--------

-NSTDLF----------------GQAD--CVVKHVLEGHDRGVNWASFHPSMPLIVSGAD

DRQIKLWRM------------N----------------DSKV------------------

------------------------------------------------------------

---------------

>UniRef90_A0A484KB78_13_265 | Coatomer subunit alpha n=3 Tax=Cuscuta sect. Cleistogrammica TaxID=1824901 RepID=A0A484KB78_9AST | E_val=8.9e-117

------------------------------------------------------------

-------------------IIAGDDYKIKVWNYK--QH-------------RCL-FTLLG

HLDYIRTVQFHHE---SPWIVSSSDDQTIRIWNWQ--SRTCISVLTGHNHYVMCASFHPK

-EDLVVSASLDQTIRVWDI--------G---------AL-KKKS-V-S-P------A---

-------------------D--------D-ILR---------------LS-Q--------

-MNTDFF----------------GGVD--AVVKYVLEGHDRGVNWASFHPTLPLIVSGAD

DRQVKIWRM------------N----------------DTKAWEVDTLRGHMNNVSCVLF

HPRQ-DIIVSNSEDKSVHVWDATKRTSLQTFRR-EH----------DRYWILASHPEINL

LAAGHDSGMIVFKLE

>UniRef90_A0A0M9G536_1_309 | Coatomer subunit alpha n=2 Tax=Leptomonas TaxID=5683 RepID=A0A0M9G536_9TRYP | E_val=6.6e-115

--MLTKFEARSSRVKAVALH--SSATWVLCGLHNGAVQIWDYRMSTCVD-------TFTE

HVG-AVRGADFHVN-QPLFVTGGDDYTVKVWNYK--LR-------------RCL-FTLNG

HMDYVRTTFFHHE---QPWIVSCSDDFTIRIWNWQ--SRKSIACLPGHNHYVMCAQFHPF

-NDLVVSGSLDKTVRVWDI--------S---------AL-RHRK-E-E-V------G---

------------------------------------------------------------

-ITQDLL----------------GTTD--VVVRYELEGHDKGVNWVAFHPCGELLLSAAD

DRTVRLWTM------------S----------------GSSCYVSRTFTGHSSNVCCAVF

Y-KN-DYVVSCGEDRTVRVVHMSSGAAVQTFRR-EV----------ERYWIMTCDPVHNL

LATGHDAGLQVFKL-

>UniRef90_A0A448WS03_5_248 | WD_REPEATS_REGION domain-containing protein (Fragment) n=1 Tax=Protopolystoma xenopodis TaxID=11 | E_val=1.7e-110

----------STSILGLAFH--PKRPWILASLHTGAIQLWDYRTCTLID-------KFEE

HEG-PVRGIDFHLN-QPLFVSGGDDFKIRIWNYK--QR-------------KCL-FTLLG

HLDYIRTTFFHKE---YPWIISCSDDQTIRIWNWQ--SRTSVSVLTGHSHYVMCSQFHPK

-EDLVVSASLDQTVRVWDI--------S---------GL-RKKN-V-A-P------S---

-----GLS--------GI-E--------E-QIRYLTG-AG--NSPSNSSG-P--------

-GNTDLF----------------GGSD--VIVKHVLEGHDRGVNWVSFHPTLPIVVSAAD

DKQIKLWRM------------T----------------A---------------------

------------------------------------------------------------

---------------

>UniRef90_A0A671KAG7_36_265 | Coatomer subunit alpha n=2 Tax=Sinocyclocheilus TaxID=75365 RepID=A0A671KAG7_9TELE | E_val=8.5e-109

------------------------------------------------------------

-----------------------------------------------------------K

HFDLLTYRYHFQE---YPWILSASDDQTIRIWNWQ--SRTCVCVLTGHNHYVMCAQFHPS

-EDLVVSASLDQTVRVWDI--------S---------GL-RKKN-L-S-P------G---

-----AA------------D--------T-DV----------------RG-I--------

-SGVDLF----------------GASD--AVVKHVLEGHDRGVNWAAFHPSMPLIVSGAD

DRQVKIWRM------------N----------------ESKAWELDTCRGHYNNVSCAVF

HPRQ-ELILSNSEDKSIRVWDMSKRTGVQTFRR-DH----------DRFWVLGAHPNLNL

FAAGHDSGMLVFKLE

>UniRef90_A0A061D4P6_1_341 | WD domain, G-beta repeat domain containing protein, putative n=1 Tax=Babesia bigemina TaxID=5866 | E_val=6.8e-105

--MLIKCKTKSPRVKGITFH--PSLHFLLASMHSGEIQLWNYLNSSLVD-------VFEY

HEG-PVRGIDFHPL-QPLFASGGDDTNVVVWDFK--QK-------------KML-FAMSG

HNDYVRTVQFHPN---YPWIVSSSDDQTVRIWNWQ--GRSCISVLQGHMHYVMCARFHPS

-KDLLVSASLDQTARIWDI--------T---------QL-REKN-C-A-I------Q---

-----GVVGQS----GSN-L----LS--NVHMIS-----------CGNGH-S---AQSDL

GRKTDRV----------------PFTD--VTCLYTLVGHEKGVNWAVFHDVLPCAITASD

DKTIRVWRF------------N----------------GPNIWQTNILKGHTNNICSLIM

HPNNINYLLSVSEDKSIKVWDTKKWTLAHTFML-EK----------DRFWIVQSAKDSNY

IAAGHDSGLIVFKL-

>UniRef90_A0A2C9JV68_698_905 | Uncharacterized protein n=1 Tax=Biomphalaria glabrata TaxID=6526 RepID=A0A2C9JV68_BIOGL | E_val=6.3e-102

------------------------------------------------------------

------------------------------------------------------------

------------E---YPWILSASDDQTIRIWNWQ--SRTCISVLTGHSHWVMCAQFHPS

-EDMVVSASSDQTVRVWEI--------S---------GL-RKKN-V-A-P------G---

-----PG---------GI-E--------D-RL----------------LN-P--------

-GQTDLF----------------GIPD--AIVKHVLKGHDRGVNWAAFHPTLPLIVSGAD

DRQIKMWRM------------N----------------ESKAWEVDTCRGHYNNVSCCLF

HPHQ-ELILSNSEDKSIRVWDMSKRTIVQTFKR-EH----------DRFWVMVAHSTLNL

FAAV-----------

>UniRef90_L8WU49_23_269 | Coatomer subunit alpha n=1 Tax=Thanatephorus cucumeris (strain AG1-IA) TaxID=983506 RepID=L8WU49_THA | E_val=2.3e-100

------------------------------------------------------------

-TG-PVRGVAIHPT-RPLLVTGGDDYKVKVWVYN--MRI-----------GTVL-NTLK-

--------------------ISASDDQTIRIWNST--SRNCIAILTGHSHYIMSAQFHPK

-DDLVVSASMDQTVRVWDI--------S---------GL-RKTG-G-T-P------A---

-----TH---------AA-A--------Q-AAGS--------------MG-T--------

-TGFDAF-----------------DTF--STVKYVLEGHDRGVNYAAFHPTLPLIVSAAD

DRQIKLWRM------------S----------------ETKAWEVDTCRGHFNNVASALF

HPRH-ELILSVGEDKTIRVWDMGKRTAVQTFRR-EH----------DRFWTLTAHPELNL

FAAGM----------

>UniRef90_A0CXK8_1_309 | WD_REPEATS_REGION domain-containing protein n=1 Tax=Paramecium tetraurelia TaxID=5888 RepID=A0CXK8_P | E_val=5.7e-97

--MIVKFHKTTERIKGLSFH--PKQPWLLVGLHSGAIQMIDYRLGRTIE-------EFVQ

HEG-PVRSVQFHQS-LCLFISGSDDFTVRVWNYK--TK-------------KCQ-FVLRG

HLDFIRCVHFHPE---LPWCVSASDDQTSRVWNYQ--SRQMLAIVTGHSHYVMHCEFHPT

-KDFLITCSLDQTIRLWSI--------A---------QL-KKRF-T-Q-K------N---

------------------------------------------------------------

-LQ--ND----------------QQNE--LELIQILEGHNQGVNWCTFSPTENLILSASD

DKKVKVWKF------------S----------------DSRGFEIDSYQGHINNVSSAMF

HPFG-DYFISNSEDNTIRLWDMKKKVEIDCFTNYEL----------DRFWVSAVHQNNNY

FAGGSDSALYIFTL-

>UniRef90_A0CRE9_1_310 | WD_REPEATS_REGION domain-containing protein n=1 Tax=Paramecium tetraurelia TaxID=5888 RepID=A0CRE9_P | E_val=4e-96

--MIVKFHKKTERIKGLSFH--PKQPWLLVGLHSGEIQMIDYRFGRTIN-------EFYE

HEG-PVRSVQFHQS-LCLFISGSDDFTVRVWNYK--TK-------------KCQ-FVLRG

HLDFVRCVNFHPE---LPWCVSGSDDQTSRIWNYQ--SRQMIATVTGHSHYVMHCEFHPS

-KDFMITCSLDQTIRLWSI--------A---------QL-KKKF-T-S-K------S---

------------------------------------------------------------

-IQ-LGE----------------QASE--LELVQILEGHSQGVNWCSFNPKDNTILSSSD

DKKIKVWKY------------F----------------DTRGYEVDQYCGHTNNVSCAMF

HPFG-EYFISNSEDKTLRLWDMKKKVEVDCFTNHEL----------DRFWICAVHQSNNY

FAGGSDSALYIFTL-

>UniRef90_A0A3B0MR36_1_327 | Coatomer alpha subunit, putative n=2 Tax=Theileria annulata TaxID=5874 RepID=A0A3B0MR36_THEAN | E_val=6.8e-95

--MLIKCKTKGTRVKGVVFH--PRLHFLLASMHSGNIQMWDYLNSTLVE-------VFSE

HEG-PVRGIDFHQE-QPLFVSGGDDTTVIVWDFT--QR-------------KKL-FVLAG

HLDYVRTVQFHTS---YPWVMSSSDDQTIRIWNWQ--SRSCITVISGHNHYVMSSLFHPT

-ENLIISSSLDHTARIWDI--------T---------YL-VEKK-C-S-I------K---

-----PPIQNQ----SNF-Y----MAEPN-SM----------------GN-A--------

-FEIEVT----------------GVSD--VICLHTLVGHSSGVNYAIFFGT-NLAITAGD

DCTVRIWRY------------S----------------QYSFYQTNILRDHEDNVTCLLL

--VK-DYLLSTSEDHSIRIWDLNTYALVHTYLM-DD----------DRFWTISKSKHNNY

ITAGHDAGLIVFKL-

>UniRef90_X6MRT7_4_219 | WD_REPEATS_REGION domain-containing protein n=1 Tax=Reticulomyxa filosa TaxID=46433 RepID=X6MRT7_RET | E_val=3.6e-93

------------------------------------------------------------

------------------------------------------------------------

----------NSE---HPWILSASDDQTVRIWNWQ--SRNSLVVLTGHNHYVMCAQWHPT

-EDLILSASLDQTIRIWDI--------S---------GL-KKRS-T-S------------

----------------VL-E--------E-SN----------------TN-K--------

-ASQDVF----------------GNVD--AVVKFVLEGHSRGVNWASFHPTMNLVVSGAD

DREIKVWRM------------S----------------GSRAWETTTYRGHLNNVSCVLF

HPKE-NVIVSNSEDKTLRVWDISRNYAPLSFRR-DS----------DRYWVLASHPRLNL

LAAGHDSGMLIFKL-

>UniRef90_A0A058Z8M4_4_214 | Coatomer protein complex, subunit alpha (Xenin) n=1 Tax=Fonticula alba TaxID=691883 RepID=A0A058 | E_val=5.6e-92

------------------------------------------------------------

------------------------------------------------------------

---------------------------TIRIWNWQ--SRTCIAVLTGHNHYVMSAEFHPK

-EDLIVSASLDQTVRVWDI--------S---------GL-RRKA-M-P-G------M---

-----GATPR-----PGS-D--------E-PT----------------KP-L--------

-LQNDLF----------------GSVD--AYVKFVLEDHDRGVNWASFHPELSLIVSGAD

DRQVKIWRY------------T----------------DTKAWHVDSCRGHYSHATCVIF

HPRQ-DLILSASEDKTLRVWDITKRTAVATFRR-EH----------DRFWIIKAHPTENL

FVTGHDAGLIVFKLE

>UniRef90_A0A165DBE3_118_333 | Coatomer subunit alpha n=3 Tax=Polyporales TaxID=5303 RepID=A0A165DBE3_9APHY | E_val=2.3e-90

------------------------------------------------------------

----------------------------------------------------CC------

------TFHLNTE---HGSQISASDDQTIRIWNST--SRNCIAILTGHSHYVMSAQFHPK

-EDLVVSASQDQTVRVWDI--------S---------GL-RKST-P-N-T----------

-------------------------------------------------A-P--------

-GTFDTF-----------------DTF--STVKYVLEGHDRGVNFATFHPTLPLIVSAAD

DRQIKIWRM------------S----------------DTKAWEVDSCRGHFNNVSVALF

HPKH-ELIVSCGEDKTVRVWDLTKRSAVQTFRR-EN----------DRFWTLAVHPELNL

FAAGHDSGLIVFKLE

>UniRef90_A0DUW0_1_309 | WD_REPEATS_REGION domain-containing protein n=1 Tax=Paramecium tetraurelia TaxID=5888 RepID=A0DUW0_P | E_val=1.7e-88

--MFVKFERHSDRVKSVSFH--PHRPWVLSALHSGIIELIDYRIKKRIA-------TYDD

HKG-AVRSVQFHPQ-LNLFCSGGDDFTVRVWNFK-----------------QCQ-FILKG

HLDYVRCVTFHPI---NPWVLSGSDDQTARVWNYQ--SRQTIGILTGHTHYIMACHFHPT

-QDFIITCSLDQTARLWNY--------G---------VL-KQRY-A-Q-K------K---

------------------------------------------------------------

-NQEYVL----------------SGAE--VQLISILDAHKDQLNWCAFHQTEPFVITSAD

DKNIKLWKY------------N----------------DTKAWEYDTLSGHTNNVCCSEF

HPKG-NVIISDSEDHTVRIWDFATRKQIGVYEN-KYF---------DRYWIVSCHQNNYY

FACGSDTMLQVFTL-

>UniRef90_A2EA68_4_309 | WD repeat protein, putative n=1 Tax=Trichomonas vaginalis TaxID=5722 RepID=A2EA68_TRIVA | E_val=2.1e-84

--MHVKLEIQSGRVKSLCFH--DSRPWLLASFHTGEIIIYDYEVGVEIQ-------RYNE

FTV-PVRTACFHPS-LPLFAAGADDTCIKIFNYD--EQ-------------RCI-ATFTE

HLDYIRTVQFHPT---KPFLVSASDDQTIRIWNYE--TNLCLTSISGHNHYVMSAFFHPT

-LPLVLSASLDDSVRVWDI--------S---------SL-FNDG-Q-S-S------G---

------------------------------------------------------------

----GIF----------------SITD--AVMKFTQEEHTAGVNWAAWHPNKPMAVSCSD

DESVKIWRI------------V----------------ETEMSLVATLRGHTGNISCACF

MPNM-DLVLSCSEDQTVRVWDSKRFVHLSKYKS-EG----------NRFWCVAAHPVKPI

FAAGHDNGLVIYS--

>UniRef90_A0A177B1W6_77_295 | Alpha-COP n=1 Tax=Intoshia linei TaxID=1819745 RepID=A0A177B1W6_9BILA | E_val=6.8e-78

------------------------------------------------------------

------------------------------------------------------------

------------E---HPWIISASDDQTCRIWNWQ--SRQCLGILAGHSNYVMCAIFHPV

-RNLVASCSLDQTIRIWDI--------S---------NL-FKKN-I-S-P------P---

-----QS---------AF-E--------E-KI----------------RA-V--------

-GRPELF----------------SIGD--FIVMHVLEGHDGAVNWVEFHKKLPILLSASD

DKLIRIWKI------------T----------------DTKATQFDTLRGHHNHVSHVIF

HPFK-DIVISNSVDQTIRFWDLNSRICIDSHRI-EN----------DRFWCLGVHPSNNL

IAAGHDSGLMVFKLE

>UniRef90_A0A7S3K9E4_2_207 | Hypothetical protein (Fragment) n=1 Tax=Euplotes crassus TaxID=5936 RepID=A0A7S3K9E4_EUPCR | E_val=3.9e-75

------------------------------------IQLFDYRVGVIIE-------KFSD

HEG-PVRSVDFHRT-QPLFTSAGDDQKIRVWDYN--KK-------------KCL-FILSG

HIDYIRMVKFHHE---LPWIISTSDDQTIRLWNWQ--NRSNIAVLTGHTHYVMSAEFHPE

-DNLILSSSLDNKVYVWDY--------S---------EL-KEKH-Q-K-Y------A---

-----GDS-------------------------------------------K--------

-KNDLMY----------------TGTE--VEVKHICEGHEKGVNFACFHPSRGLIASGAD

DKLIKLWRM------------S----------------GSRAWELD--------------

------------------------------------------------------------

---------------

>UniRef90_A0A4Q9LJL8_5_306 | WD40 domain-containing protein (Fragment) n=1 Tax=Hamiltosporidium magnivora TaxID=148818 RepID= | E_val=1.6e-73

---YTQMEKKTQRVKSVAFH--HTKPILLTAHHTGTIKAYDYQTSSLIH-------EFND

HEG-PVRTIIFHPF-INIFVSGGDDCIIRIWSYN--ER-------------KII-TKLKG

HTDYIRCLDFHPN---LPWILSSSDDQTIKIWNFQ--SYKLLATLTGHSHYVMSAKFLT-

-QDYIISASLDQSLRLWDC--------K---------SL-KTPK-N-K-K------S---

------------------------------------------------------------

---IDIL----------------KVPE--VIVRQILDGHEKGINWISVNPNTTMFATSSD

DRQIKIWEL------------V----------------GEIVKERDVFHHHQGNVSSVLF

--VN-NLLISNGEDGILCIYDLKNKKSKKIVIE-------------DRFWCITTNRDSTL

FCVGYDTGFKVFSV-

>UniRef90_V6LG68_6_338 | Coatomer alpha subunit n=1 Tax=Spironucleus salmonicida TaxID=348837 RepID=V6LG68_9EUKA | E_val=1.7e-68

------FNLQTTRVKSVAFH--PRRPWVLVACHDGFVELHDYVVGALVE-------RFMA

HSA-PVRACAFHES-QPIFATGSDDRQIKLWDLE--NR-------------KEI-ASLQG

HTDYIRFIEFHKI---RPWIVSASDDCSVKIWNWQ--SRSCVGSCTGHKEFAMCAKFHPR

-DDLLATAGLDSTVRVWDL--------S---------KI-SRSV---GLK------K---

-----NIV-----------E-------------------------------K--------

-TLQQVL----------------NLPQCTLSISINGESHTRGVNWVSWGENDSL-YSSSD

DYSVREWRLLRGKPEKDEILGI----------------DATLYTHSTLTGHSGNVNAVEV

G-KD-NVVVSASSDGTIRVYEQKSKTYIGNLTC-QDYGIKPQNSMPTRFWCLSEHPKQSL

WAAGHDQGLIIFNI-

>UniRef90_A0A1J4K2J0_3_312 | Coatomer subunit alpha-3 n=1 Tax=Tritrichomonas foetus TaxID=1144522 RepID=A0A1J4K2J0_9EUKA | E_val=2.1e-65

----VKLDLESSRVKGVAFH--PNRPWVLYCTHTGTVHLHDYEIGVELACY-----KVTN

DNI-PVRCVAFHPT-QPLFACGTDNYEVIVYNWA--RK-------------MKL-FTLTG

HLDYVRSVEFHAV---YPLLISSSDDSTARIWNWQ--SRCCIAILEEHTYFVMCARFNPQ

-KPLIATASLDDCVRVFNV--------N---------TL-FNSS-M-S-K------D---

------------------------------------------------------------

-VESSFF----------------SMED-TSVLTSELEEHPEGADCVAWDQSGNKLYSCGE

DSRIKIYNI------------V----------------NDEATLSNQIFTHNGAVTGIAV

HYPT-SNFVSVSEDSTLRVFDGSTNQQIAKFEV-PV----------SRFWCVACHPKDAL

IAAGHDRGLVILK--

>UniRef90_E0S5J9_10_298 | WD40 repeat-containing coatomer complex protein n=1 Tax=Encephalitozoon intestinalis (strain ATCC 50 | E_val=2e-63

------MEKEASRVKSLSFH--PTKPVIISGHHSGSIKAWDYQMGVCIH-------EFLD

HDG-SVRAVLFHPR-GDFFVSGGDDKVIRVWSYT--ER-------------RIT-NKLRG

HDDFIRSLDFHPT---KPWILSASDDQTIMVWNML--TGKLLATARGHCHYVMAAKFLGE

-ES-IVSGSLDQSIRIWDC--------K---------GL-KEGG-K-K-N------S---

------------------------------------------------------------

-----------------------LLPD--IVIKQIVDGHDRGINAIAAKD--GVFVSGGD

DRDIKCWEW------------S----------------ETSVWEKEVMYNHQGPVTGLLC

--DR-EYVLSSGEDGLFSIYNTETRKSIERRTE-------------GRYWCVANK--GNL

YAAGHDSGFEVY---

>UniRef90_Q8SSJ2_10_298 | COATOMER COMPLEX n=2 Tax=Encephalitozoon cuniculi TaxID=6035 RepID=Q8SSJ2_ENCCU | E_val=5.2e-61

------MEKETSRVKSLSFH--PSKPVIISGHHSGSIRAWDYQMNVCIH-------EFLE

HDG-SVRAVLFHPR-GDFFVSGGDDKIIRVWSYS--ER-------------RVT-NRLKG

HDDFVRSLDFHPT---KPWILSASDDQTIMVWNML--TGKLLATARGHCHYVMAARFLGE

-ES-IVSGSLDQSIRVWDC--------K---------GL-KEGS-K-K-N------S---

------------------------------------------------------------

-----------------------LLPD--IVIKQIVDGHDRGINAIAVRG--EVFVSGGD

DRDIKCWEW------------S----------------ETSVWEKEVMYNHQGPVTGLLC

--DG-NHVLSSGEDGLFSIYNMESRKSVECRTE-------------GRYWCVASR--GNL

YAAGHDSGFEVY---
