## Supplementary Figure S1 for "An Extended Motif in the SARS-CoV-2 Spike Modulates Binding and Release of Host Coatomer in Retrograde Trafficking"

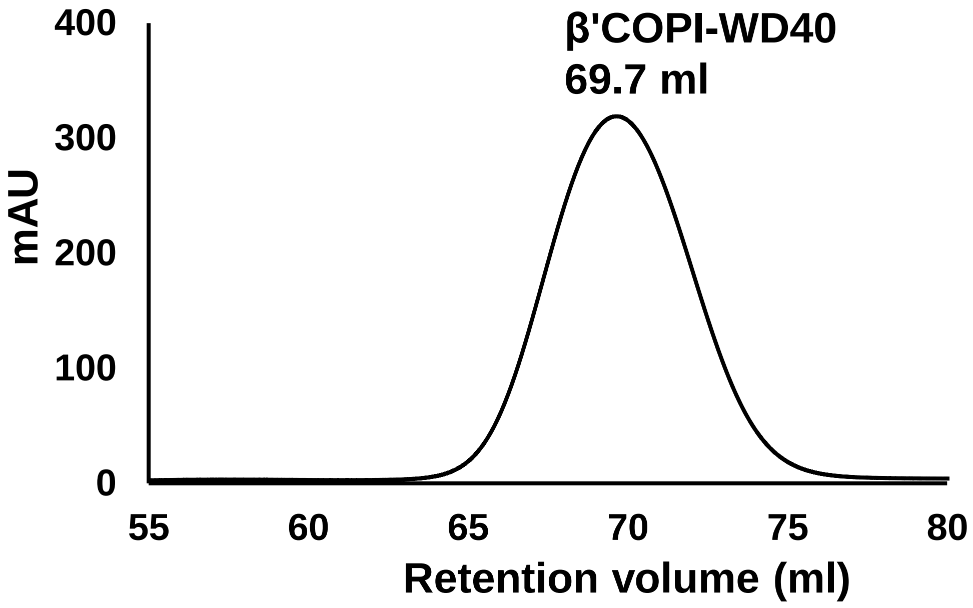


**SI Figure S1: Purification of β'-COPI-WD40 domain.** SEC analysis of β'COPI-WD40 protein showing a single monodisperse peak. A HiLoad Superdex 75 16/600 chromatography column was used for this analysis.
