## Supplementary Figure S2 for "An Extended Motif in the SARS-CoV-2 Spike Modulates Binding and Release of Host Coatomer in Retrograde Trafficking"

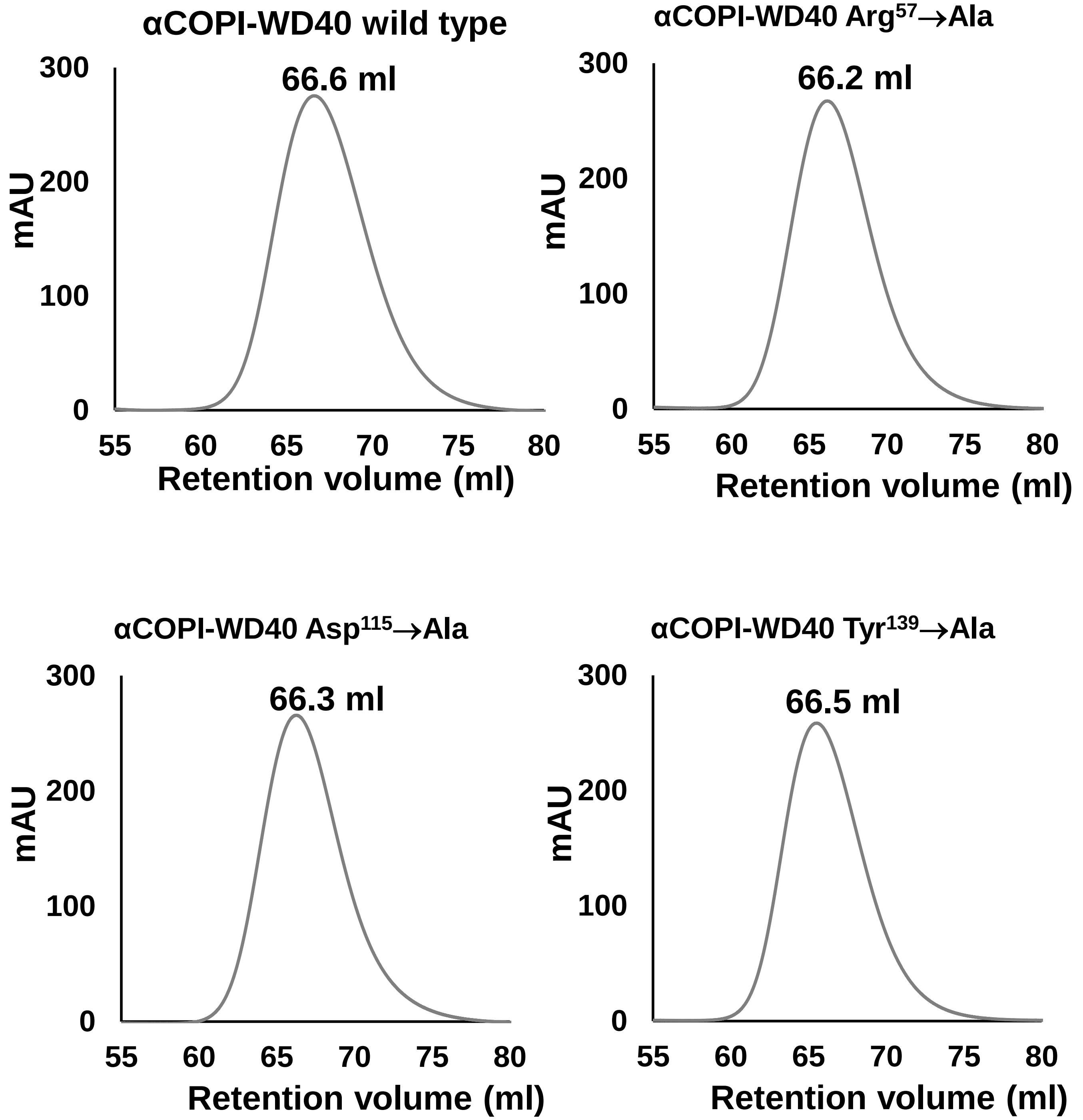


**SI Figure S2: Preparative SEC analysis of αCOPI-WD40 domain and mutants Arg$^{57}$→Ala, Asp$^{115}$→Ala, and Tyr$^{139}$ →Ala mutants.** A HiLoad Superdex 75 16/600 chromatography column was used for this analysis.
