## Supplementary Figure S3 for "An Extended Motif in the SARS-CoV-2 Spike Modulates Binding and Release of Host Coatomer in Retrograde Trafficking"

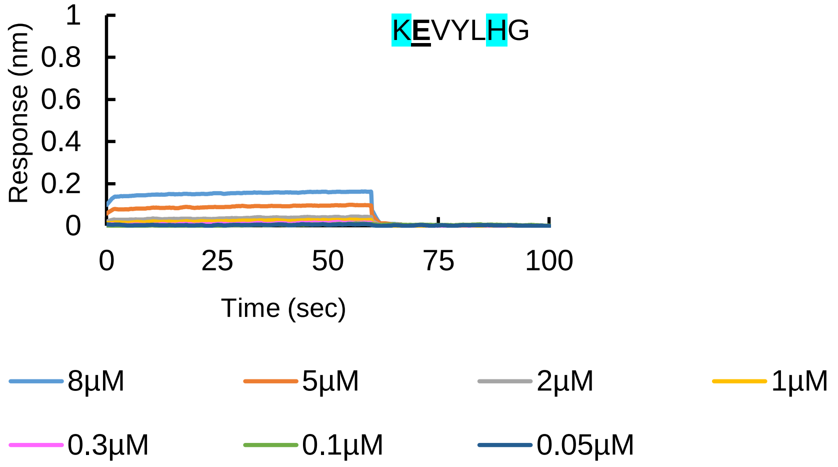


**SI Figure S3: BLI analysis of a spike hepta-peptide with a scrambled sequence containing the Thr$^{1273}$→Glu mutation.** This analysis shows weak binding of αCOPI-WD40 domain to this peptide. The color code corresponding to the αCOPI-WD40 concentrations is given at the bottom of the figure.
