## Supplementary Figure S4 for "An Extended Motif in the SARS-CoV-2 Spike Modulates Binding and Release of Host Coatomer in Retrograde Trafficking"

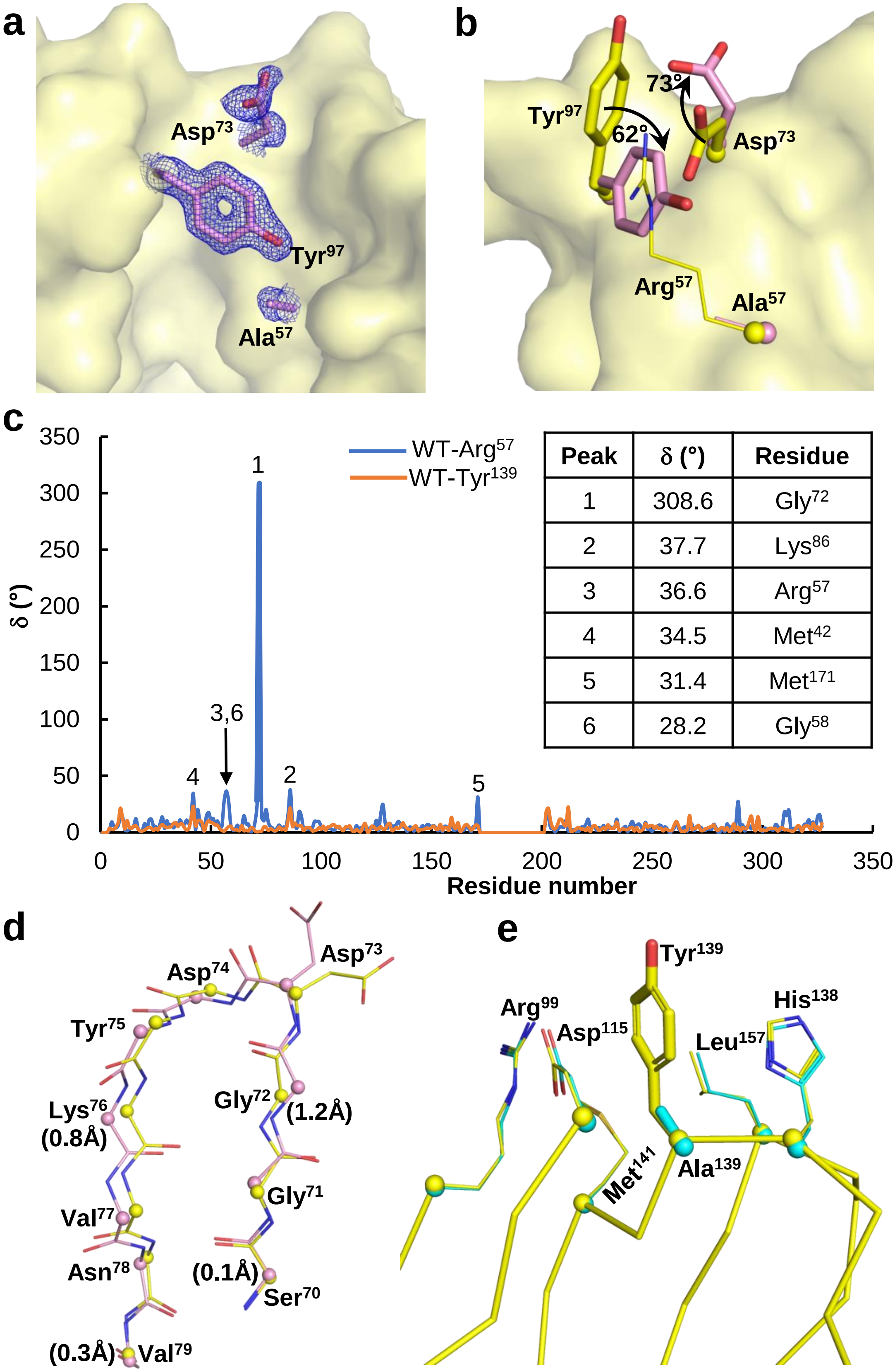


| Peak | δ (°) | Residue |
|---|---|---|
| 1 | 308.6 | Gly72 |
| 2 | 37.7 | Lys86 |
| 3 | 36.6 | Arg57 |
| 4 | 34.5 | Met42 |
| 5 | 31.4 | Met171 |
| 6 | 28.2 | Gly58 |

**SI Figure S4: Crystallographic analysis αCOPI-WD40 Arg$^{57}\rightarrow$Ala and Tyr$^{139}\rightarrow$Ala mutants. (a)** Electron density around the side chains of residues Ala$^{57}$, Asp$^{73}$, and Tyr$^{97}$ (blue mesh, 2Fo-Fc map contoured at 1.0 σ) in the structure of the Arg$^{57}\rightarrow$Ala mutant. For simplicity, other residues are shown as a yellow surface. **(b)** Conformational changes in αCOPI-WD40 caused by Arg$^{57}\rightarrow$Ala mutation. The mutant and wild type αCOPI-WD40 structures are shown in pink and yellow as the primary colors, respectively. The Arg$^{57}\rightarrow$Ala mutation generates a cavity in αCOPI-WD40. The nearby Tyr$^{97}$ residue side chain rotates into this cavity. This is accompanied by the outward rotation of the Asp$^{73}$ side chain, which is bonded to Arg$^{57}$ in the wild type structure. **(c)** An analysis of differences in main chain conformation

**SI Figure S4 (contd):** between the wild type and mutant αCOPI-WD40 crystal structures. The difference in Ramachandran angles was calculated for each residue, ($\delta=\sqrt{((\psi_{WT}-\psi_m)^2+(\phi_{WT}-\phi_m)^2)}$), where ($\psi_{WT}$, $\phi_{WT}$) and ($\psi_m$, $\phi_m$) are Ramachandran angles for wild type and each mutant crystal structure. This analysis shows larger conformational changes in the $Arg^{57}$→Ala mutant (blue) than in $Tyr^{139}$→Ala mutant (orange). The top six peaks are highlighted. Peak 1 corresponds to a main chain rearrangement coincident with an outward movement of $Asp^{73}$ as shown in panel **(d)**. Cα atoms are shown as spheres in panel **(d)**. Upto 1.2Å and 0.8Å shifts in the Cα atoms are observed for $Gly^{72}$ and $Lys^{76}$ respectively, between the wild type and mutant structures. The intervening residues demonstrate substantial conformational rearrangement of the main chain. **(e)** In contrast, the αCOPI-WD40 $Tyr^{139}$→Ala mutant structure (primary color cyan) shows limited changes from the wild type structure (primary color yellow).
